# Predicting children’s literacy from task-based fMRI: Neural heterogeneity and task-dependent performance

**DOI:** 10.64898/2026.04.17.719130

**Authors:** Gustavo S. P. Pamplona, Svenja Stettler, Bruno H. Vieira, Sarah V. Di Pietro, Nada Frei, Christina G. Lutz, Iliana I. Karipidis, Silvia Brem

## Abstract

Reading is a complex skill with a well-characterized neural basis. Multivariate fMRI analyses have deepened our neuroscientific understanding of literacy by linking neural patterns to behavioral traits. Although task-based fMRI often outperforms resting-state fMRI in predicting cognitive traits, few studies have applied it to continuous measures of children’s reading ability. To identify neural markers of literacy, we compared predictive performance across multiple fMRI tasks and reading-related measures. In this data-driven study, we predicted literacy skills in school-aged children (6.7–10.3 years) from eleven behavioral scores grouped into Reading (fluency and comprehension), Verbal (vocabulary knowledge and verbal intelligence), and Naming (object naming speed). Predictive performance was examined across four fMRI tasks completed by subgroups of children (n = 73–97): two active tasks – phonological-lexical decisions (PhonLex) and audiovisual character learning (Learn) – and two passive tasks – word and face viewing (Localizer) and character processing (CharProc). Individual activation contrast maps, categorized as simple (single condition) or subtractive (condition contrasts), were analyzed using a machine learning model with whole-brain predictors derived from principal component analysis. Results showed the highest predictive performance for Reading and Naming with PhonLex > Learn > Localizer = CharProc, and for Verbal with PhonLex = Learn > Localizer = CharProc. Simple contrasts generally outperformed subtractive contrasts in predicting behavioral scores. Key neural predictors, identified through whole-brain and region-of-interest analyses, included the left inferior frontal gyrus, supramarginal gyrus, ventral occipitotemporal cortex, insula, and default mode network regions. Together, these findings indicate that, for predicting literacy traits in children, active tasks and tasks that engage brain systems involved in multisensory learning tend to outperform both passive paradigms and simple subtractive task contrasts. This study provides a methodological benchmark for brain-based prediction of reading ability and highlights the value of activation heterogeneity across distributed regions as a potential marker for tracking literacy development over time.

## 1. Introduction

Reading acquisition involves the development and coordination of distributed brain networks that integrate visual, linguistic, and cognitive processes. Rather than relying on a single brain module, reading emerges from the dynamic interactions among multiple brain regions that gradually adapt and specialize as language and literacy skills develop (Benischek et al., 2020; Chyl et al., 2019; Dehaene and Cohen, 2007; Di Pietro et al., 2023; Fedorenko et al., 2024; Martin et al., 2015; Schlaggar and McCandliss, 2007; Tomasi and Volkow, 2012; Turker et al., 2025; Wise Younger et al., 2017). Within this broader architecture, a functionally specialized language network in left frontal and temporal cortex supports high-level linguistic processing, while interacting with adjacent and domain-general systems (Fedorenko et al., 2024). Among these brain regions, the inferior frontal gyrus (IFG), particularly its posterior portion often referred to as Broca’s area, contributes to phonological and semantic processing, as well as to aspects of language comprehension and production (Fedorenko et al., 2011; Sahin et al., 2009). The posterior insula, which is strongly connected to sensory and associative cortices (Namkung et al., 2017), has been implicated in phonological processing and audiovisual language integration (Łuniewska et al., 2019; Wang et al., 2020). The superior temporal gyrus (STG), including Wernicke’s area and the planum temporale, supports phonological processing of speech and print and is a key site for audiovisual integration during letter-speech sound processing (Blau et al., 2010; Humphries et al., 2005; Turker et al., 2025; van Atteveldt et al., 2004; Vigneau et al., 2006). Adjacent parietal regions, such as the supramarginal gyrus (SMG), are consistently linked to individual differences in reading and phonological skills (Beyer et al., 2022). The fusiform gyrus, particularly the sites referred to as visual word form area(s) (VWFA), plays a crucial role in the perceptual and lexical processing of visual input during reading (Brem et al., 2006; Caffarra et al., 2021; Dehaene and Cohen, 2011; Kronbichler et al., 2007; Lerma-Usabiaga et al., 2018; Li et al., 2024; McCandliss et al., 2003; Vinckier et al., 2007). In addition, the default mode network (DMN), including the posterior cingulate, medial prefrontal cortex, and angular gyrus is engaged during internally oriented cognition and higher-order aspects of language and narrative processing (Andrews-Hanna et al., 2014; Greicius et al., 2003; Spreng, 2012). Its self-generated cognitive processes have been proposed to support discourse-level language comprehension and writing, but can also contribute to task-unrelated, internally focused thoughts during reading (Andrews-Hanna et al., 2014; Brownsett and Wise, 2010; Musso et al., 1999; Zhang et al., 2022).

Many studies have used functional magnetic resonance imaging (fMRI) to investigate the relationship between reading processes and brain activity or functional connectivity. fMRI provides high spatial resolution and deep brain coverage, making it a powerful tool for brain mapping. Despite methodological challenges, acquiring fMRI data from children offers valuable insights into the development and acquisition of literacy skills. Most commonly, such data are analyzed using univariate and correlational approaches, which assess associations between neural activation and behavioral measures. However, this approach does not account for the complexity of voxel-wise activity patterns underlying reading processing and development. To address this limitation, predictive modeling using machine-learning algorithms has been applied to fMRI data (Varoquaux and Poldrack, 2019; Yarkoni and Westfall, 2017). These models then tend to be more sensitive in establishing associations between brain activity and behavioral aspects than mass univariate methods by aggregating information across the entire brain (Woo et al., 2017), allowing for a more nuanced understanding of brain mechanisms. Predictive studies can be classification-based, relying on categorical group labels, or they can use regression analysis to model continuous behavioral score distributions. These prediction-based approaches may thus offer a more comprehensive framework for understanding the neural basis of reading and literacy development.

Several studies have been conducted to predict individual literacy characteristics using neuroimaging. Research has demonstrated that literacy skill prediction is feasible through various imaging modalities, including structural MRI (Beyer et al., 2022; He et al., 2013; Kristanto et al., 2020), diffusion MRI (Cui et al., 2016; Kristanto et al., 2020), resting-state fMRI (Kristanto et al., 2020; Tomasi and Volkow, 2020; Xu et al., 2015; Yuan et al., 2023) and task-based fMRI (Bach et al., 2013; Feng et al., 2021; Hoeft et al., 2007; Tomasi and Volkow, 2020; Tomaz Da Silva et al., 2021; Zahia et al., 2020). These studies provide insights into the brain regions primarily responsible for successful prediction performance, based on individual variance in given neural features. A common finding across this line of prediction studies is the importance of reading-related brain regions, such as the left fusiform gyrus and the left superior temporal gyrus, in classifying individuals with and without dyslexia. Additionally, large-scale networks, including the frontoparietal and default-mode networks play a significant role in literacy-related predictions (see Nakai et al., 2024, for a review, and Koyama et al., 2011; Schurz et al., 2015).

Increasing evidence suggests that task-based fMRI yields higher predictive efficacy for cognitive skills than resting-state fMRI (Greene et al., 2018; Sripada et al., 2020; Tomasi and Volkow, 2020). In line with this, some studies have employed task-based fMRI to predict individual literacy characteristics, primarily through classification approaches that distinguish between groups based on reading or language skills, such as typical readers and individuals with dyslexia. However, reading skills develop and manifest along a continuum, and studies predicting continuous variables, rather than classification, could offer valuable insights into the full spectrum of literacy skills. According to a recent systematic review (Nakai et al., 2024), among task-based fMRI studies using machine learning to predict continuous measures of reading ability, rather than relying on group classifications such as poor vs typical readers, only two have been conducted in adults (Feng et al., 2021; Tomasi and Volkow, 2020) and just one seminal study has been conducted in children (Hoeft et al., 2007). Task-based fMRI studies often involve smaller sample sizes than structural MRI or resting-state fMRI studies (Nakai et al., 2024), which can limit statistical power and generalizability. Large-scale datasets, such as the Human Connectome Project (HCP) (Tomasi and Volkow, 2020) or the Adolescent Brain Cognitive Development study (ABCD) (Casey et al., 2018), help address this issue but were not specifically designed for literacy research. For instance, the HCP includes only two language– and reading-related measures (i.e., the Oral Reading Recognition Test and the Picture Vocabulary Test (Hodge et al., 2016)) which provide only a narrow view of literacy skills, and neither the HCP nor the ABCD study includes a functional MRI task related to reading. In the present study, we address this gap by working with a dataset that combines a decent sample size with a broad and detailed assessment of reading, enabling the prediction of multiple facets of reading ability from fMRI tasks beyond those available in the HCP or ABCD.

In this study, we aimed to determine whether fMRI tasks designed to assess neural activity related to different aspects of reading vary in their ability to predict literacy skills. Additionally, we used predictive maps to better characterize the brain regions supporting distinct components of literacy. To this end, we analyzed data from a comprehensive assessment of reading-related skills in 105 elementary-school-aged children and derived three summary scores indexing (i) reading fluency and comprehension, (ii) vocabulary knowledge and verbal intelligence, and (iii) object naming speed.

For subgroups of these children, we further analyzed fMRI data acquired during four tasks probing neural activity related to phonological–lexical decisions, artificial grapheme–phoneme correspondence learning, passive visual processing of words and faces, and passive visual processing of characters and false fonts with varying familiarity. Using individual condition-specific activation maps from these tasks, we applied a machine-learning-based approach to predict the behavioral summary scores for each child. Activation voxels served as predictors, and principal component analysis was used for dimensionality reduction before model fitting. We then compared predictive performance across fMRI tasks and identified key brain regions that predominantly drove these predictions. We hypothesized that more cognitively demanding tasks, and those more closely resembling natural reading, would yield superior predictions of literacy skills, and that regions traditionally implicated in reading would make prominent contributions to these models. The findings aim to advance methodological approaches for predicting children’s reading abilities from task-based neural activation and provide insights into the neural systems that underpin individual variability in literacy.

## 2. Materials and Methods

### Participants

We used data from a project that studied reading development, audiovisual learning, and the effects of app-based phonics training in school children with varying reading skills. Here, we used fMRI and behavioral data collected at the first time point of this longitudinal study, i.e., before the phonics training was administered. A total of 105 German-speaking children, aged 6.7 to 10.3 years and enrolled in grades one through four, participated in the study. The children underwent extensive behavioral assessments of reading and other cognitive abilities, along with an MRI session. MRI measurements were conducted at the MRI Center of the Psychiatric Hospital, University of Zurich, Switzerland. Six children did not complete the full battery of behavioral measures and were not considered for the calculation of summary scores (see Materials and Methods – “Behavioral measures”). The demographic information of the sample of 99 children included is presented in Table 1.

**Table 1.** Summary statistics for the demographic characteristics of the 99 children used for the calculation of summary scores.

| Phenotype | Mean ± standard deviation<br>(range)* |
| --- | --- |
| Gender: female / male | 44 / 55 |
| Age at acquisition in years | 8.7 ± 0.8 (6.7–10.3) |
| Handedness: right / left / ambidextrous** | 86 / 11 / 2 |
| ARHQ | 0.38 ± 0.13 (0.09–0.70) |
| RIAS Total IQ | 102.18 ± 9.17 (83–123) |
| RIAS Verbal IQ | 99.47 ± 10.71 (76–121) |
| RIAS Non-verbal IQ | 104.46 ± 7.48 (86–123) |
| SLRT-II words | 35.41 ± 31.87 (1–99) |
| SLRT-II pseudowords | 37.92 ± 30.36 (1–99) |
| SLS | 87.71 ± 19.19 (62–138) |
| ELFE-II | 36.99 ± 31.37 (0.6–99.4) |
| HRT | 46.69 ± 32.21 (2–99) |
| Schreib.On | 39.74 ± 32.58 (1–100) |
| PPVT | 54.47 ± 29.13 (8.1–98.9) |
| RAN simple | 0.83 ± 0.21 (0.42–1.50) |
| RAN complex | 0.62 ± 0.17 (0.20–1.02) |

Note: Values shown are standardized by age or school class and converted to percentiles, when applicable (i.e., for SLRT-II, ELFE-II, SLS, HRT, Schreib.On, and PPVT). \* = except for gender and handedness, where proportions are shown. \*\* = measured with the Edinburgh Handedness Questionnaire. Abbreviations: ARHQ = Adult Reading History Questionnaire (the higher score from both parents was used), RIAS = Reynolds Intellectual Assessment Scales, IQ = intelligence quotient, SLRT-II = Salzburger Lese- und Rechtschreibtest, SLS = Salzburger Lese-Screening, ELFE-II = Ein Leseverständnistest für Erst- bis Siebtklässler, HRT = Heidelberger Rechentest, PPVT = Peabody Picture Vocabulary Test, RAN = Rapid Automatized Naming.

### Behavioral measures

Several cognitive domains, including intelligence, reading, and verbal skills, were assessed in the children participating in our study using a series of behavioral tests. We included a total of eleven behavioral measurements (Table 1):

- Estimates of verbal (Verbal_IQ) and non-verbal intelligence (NonVerbal_IQ), based on the Reynolds Intellectual Assessment Scales (RIAS) (Hagmann-von Arx and Grob, 2014; Reynolds and Kamphaus, 2003);
- Measures of reading fluency for words (Word_Reading) and decoding fluency of pseudowords (Pseudoword_Reading), assessed using the Salzburg Reading and Spelling Test (*Salzburger Lese– und Rechtschreibtest – SLRT II*; Moll and Landerl, 2010). These measures reflect the number of correct words or pseudowords read within one minute;
- A measure of sentence reading fluency (Sentence_Reading), using the Salzburg Reading Screening (*Salzburger Lese-Screening – SLS*; Wimmer and Mayringer, 2016);
- A measure of reading comprehension (Reading_Comprehension), based on the Reading Comprehension Test for First to Seventh Graders (*Ein Leseverständnistest für Erst-bis Siebtklässler – ELFE-II*; Lenhard et al., 2018);
- A measure of arithmetic skills (Arithmetics), derived from the Heidelberg Arithmetic Test (*Heidelberger Rechentest – HRT;* Haffner, 2005);
- A measure of written spelling or orthography (Written_Spelling), based on the Schreib.On Test (May, 2008);
- A measure of vocabulary knowledge (Vocabulary), assessed with the Peabody Picture Vocabulary Test (PPVT-4; Dunn and Dunn, 2007);
- Measures of naming simple (RAN_Simple) and complex objects (RAN_Complex), based on the Rapid Automatized Naming Test (RAN; (Mayer, 2020)). The RAN_Simple subtest included images representing highly frequent mono– and bisyllabic words, while the RAN_Complex subtest consisted of images representing multisyllabic, more complex words. Children were instructed to name the objects from top to bottom as quickly as possible. For each subtest, the score was calculated as the number of items named per second. This test assesses the speed and accuracy of naming familiar stimuli.

When applicable, the behavioral measures (as shown in Table 1) were standardized by age or school class using percentiles. This standardization was performed for SLRT-II, ELFE-II, SLS, HRT, Schreib.On, and PPVT. Next, all standardized measures were converted to T-scores to normalize the scores across subjects within each measure, which are scaled to have a mean 50 and a standard deviation of 10, to maintain consistency. Handedness was assessed using the Edinburgh Handedness Questionnaire (Oldfield, 1971). The familial risk for dyslexia was assessed using the Adult Reading History Questionnaire (ARHQ) (Lefly and Pennington, 2000). Notably, the ARHQ revealed that 40 of the children were at risk for dyslexia, defined as scores from one of the parents greater than 0.4 (Maurer et al., 2003). Therefore, our study encompassed a broad spectrum of reading abilities and familial risk for dyslexia to account for variability.

### Tasks performed during fMRI acquisition

The data also included fMRI scans collected while children performed four tasks in the scanner: a phonological lexical decision task (hereafter referred to as the “PhonLex” task), a grapheme-phoneme correspondence learning task (“Learn” task), a passive viewing task of words and faces (“Localizer” task), and a passive visual processing of characters and false fonts (“CharProc” task). All tasks were implemented and administered using the Presentation® software (v. 20.1, www.neurobs.com).

*<u>PhonLex task (</u>Fig. 1A).* The PhonLex task is a phonological lexical decision task, during which children performed phonological-lexical decisions for letter and false-font strings (Di Pietro et al., 2023; Kronbichler et al., 2007). Children were presented with word strings in four types of experimental stimuli: (i) “word” condition, containing frequently used short German words with real meanings (e.g., *Zug*, meaning “train”); (ii) “ff” condition, where strings were written in a fictional alphabet (false fonts), as a non-lexical and visual control condition; (iii) “pseudoword” condition, showing pronounceable pseudowords in the Latin alphabet with no meaning (e.g., *Tebt*); and (iv) “pseudohomophone” condition, with pseudo-homophones that were orthographically wrong but sounded like real words (e.g., *Tsug*, which in German sounds like *Zug*). Children indicated via a response pad whether the string sounded like a real word (i.e., in the “word” and “pseudohomophone” conditions) or not (i.e., in the “ff” and “pseudoword” conditions), by responding ‘yes’ or ‘no’, respectively. Each presentation lasted 3500 ms, regardless of the child’s response, followed by a fixation cross for 1350 ms. Trials were presented in mini-blocks, consisting of one to three presentations per condition, with 5-second breaks every 15 iterations. In total, 60 trials were presented per run, leading to a total duration of 6min05s for each run of the PhonLex task.

**Figure 1.**
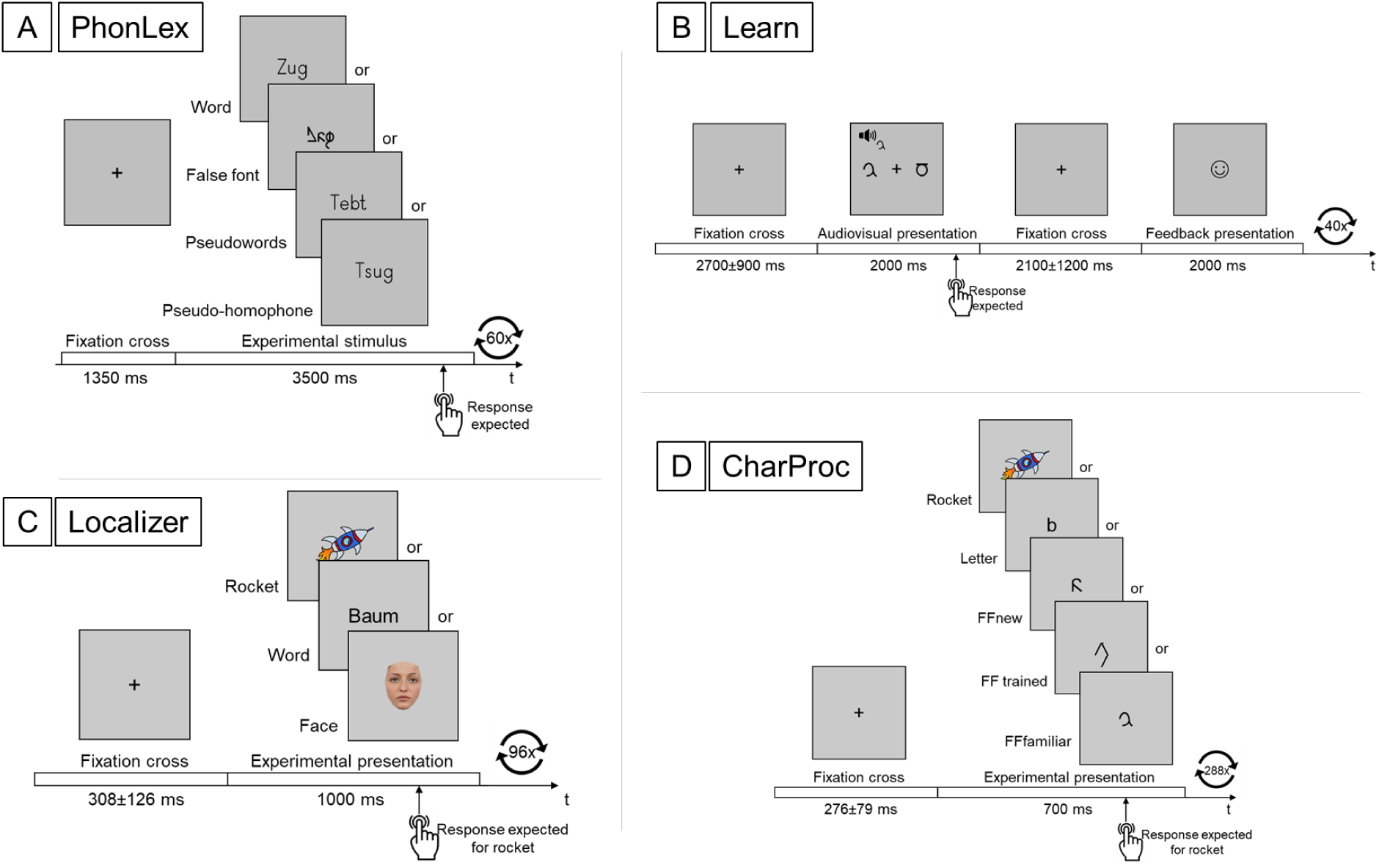
Task design and stimulus sequence for the (A) PhonLex, (B) Learn, (C) Localizer, and (D) CharProc tasks.

*<u>Learn task (</u>Fig. 1B).* In the Learn task, children were required to learn novel correspondences between unfamiliar false-font symbols and familiar German speech sounds, based on feedback received after each trial (Fraga-González et al., 2025; Frei et al., 2025). In each trial, children simultaneously saw two symbols while hearing one speech sound and were asked to match the speech sound to the correct symbol. The associations between symbols and speech sounds were unfamiliar to the children prior to the task. The children indicated the correct association by pressing either the left or right button on a response pad to select the corresponding symbol. Feedback after each trial showed whether the child’s response was correct, represented by a happy smiley, or incorrect, represented by a neutral smiley. Therefore, trials in this task could be classified based on the response as either correct or incorrect. In the context of our study, “correct” trials represent associations that have been learned, while “incorrect” trials represent associations that are still being learned. Children were instructed to respond in every trial and to guess if unsure about the association, to learn it via feedback. The feedback for slow responses was the German word “schneller” (faster) to prevent inactivity. Each run included four distinct symbols and four speech sounds, presented in all possible combinations, allowing children to learn the associations over time. Both the audiovisual and feedback presentations lasted 2000 ms each, regardless of the child’s response. Following the audiovisual presentation, a fixation cross appeared with an average jittered interval of 2500 ± 500 ms, and similarly after the feedback presentation, with an average jittered interval of 2000 ± 500 ms. Four symbol-sound associations could be learned, with each speech sound repeated across ten trials, totaling 40 associations per run. The total duration of each run of the Learn task was 5min54s.

*<u>Localizer task (</u>Fig. 1C).* In the Localizer task, children passively viewed either real words or neutral faces. Word stimuli comprised four-letter German words (condition “word”), while face stimuli consisted of neutral, frontal images of Caucasian male and female faces (condition “face”). The task included twelve blocks in total: six displaying 6-8 words and six blocks displaying 6-8 faces. In each block, a rocket figure appeared 0-2 times, prompting children to press a button to signalize task engagement. Each block contained eight trials, featuring either words and rockets or faces and rockets, randomly presented. Words were displayed for 1000 ms and faces were displayed for 1000 ms (in 14 runs) or 1200 ms (in the remaining runs). After each word or face presentation, a fixation cross appeared for an average jittered interval of 308 ms (range: 128-498 ms). Between blocks, a longer fixation cross was presented for 5000 ms. Each run consisted of 96 trials. The total duration of each run of the Localizer task was 3min28s.

*<u>CharProc task (</u>Fig. 1D).* In the CharProc task, children were presented with either real characters or false fonts with which they had different levels of familiarity (Pleisch et al., 2019a). Children were presented with one of four types of stimuli: (i) real letters (“letter” condition – high visual and auditory familiarity); (ii) false fonts previously introduced in the Learn task (“FFtrained” condition – i.e., visual and phonological knowledge acquired through audiovisual association learning during the Learn task), (iii) false fonts introduced visually during a pre-experiment practice phase (“FFfamiliar” condition – visual familiarity but no acquired phonological associations), or (iv) novel false fonts unfamiliar to the participant (“FFnew” condition new, unfamiliar false-font characters). For the FFfamiliar characters, the familiarization phase was conducted as a paper-and-pencil task immediately before the CharProc task, during a break outside the scanner. This familiarization consisted of 80 trials in which three symbols were presented, two of them being identical. The child was asked to mark the matching symbols with a pen. By the end of this phase, the child was visually familiar with eight symbols, each presented 10 times (the ones presented as FFfamiliar). The fMRI task consisted of 32 blocks – eight blocks per condition – with each block containing eight stimuli presentations. To ensure task engagement, a rocket figure appeared an average of once per block (range 0-2 times), prompting children to press a button. Each block included 8-10 trials, which featured either letters and rockets or false fonts and rockets, randomly presented. Stimuli (letters or false fonts) were presented for 711 ms, followed by a fixation cross for a jittered interval averaging 276 ms (range: 178-435 ms). This sequence was repeated a total of 288 times, derived from 4 conditions x 8 blocks per condition x 9 stimuli presentations per block (8 for experimental stimuli and an average of 1 rocket figure). Between blocks, a longer fixation cross appeared for an average of 6000 ms (range: 3000-9000 ms). The total duration of each run of the CharProc task was 8min09s.

Every task included a starting and a finishing screen for 5000 ms and 8000 ms, respectively. Button presses and response times were recorded for each task and participant for posterior data analysis.

### fMRI data acquisition

Neuroimaging data included both anatomical and functional images, acquired using a Philips Achieva 3 T MRI scanner equipped with a 32-channel head coil (Philips Healthcare, Best, The Netherlands). Visual stimuli were delivered through video goggles (VisualStimDigital, Resonance Technology, Northridge, CA), while auditory stimuli were presented via headphones (MR Confon). fMRI data were collected using a T2*-weighted fast echo-planar imaging (EPI) pulse sequence. Imaging parameters varied depending on the task: PhonLex Task – echo time/repetition time (TE/TR) = 30/1000 ms, voxel size = 3 × 3 × 3.5 mm³, slice gap = 0.5 mm, matrix size = 64 x 64, 32 slices, and 366 volumes; Learn Task – TE/TR = 35/1330 ms, voxel size = 3 × 3 × 3 mm³, slice gap = 0.3 mm, matrix size = 64 x 64, 42 slices, and 273 volumes; Localizer Task: TE/TR = 35/1250 ms, voxel size = 2.46 × 2.46 × 3 mm³, slice gap = 0.3 mm, matrix size = 80 x 80, 42 slices, and 168 volumes; CharProc Task: TE/TR = 35/1250 ms, voxel size = 2.46 × 2.46 × 3 mm³, slice gap = 0.3 mm, matrix size = 80 x 80, 42 slices, and 408 volumes. For all functional imaging a flip angle of 80° was used. Volumes were acquired in a descending continuous order, with a multiband factor of 2 to ensure full cerebrum coverage. To achieve steady-state magnetization, five dummy scans were performed before each functional run. High-resolution anatomical MR images were obtained using a T1-weighted Magnetization-Prepared Rapid Gradient Echo (MPRAGE) pulse sequence. Parameters were as follows: TE/TR = 3.16/6.98 ms, voxel size = 1 x 0.94 x 0.94 mm³, 3D matrix size = 176 x 288 x 288, flip angle = 8°.

### Data analysis

All behavioral and MRI data were analyzed with Python v3.9.19 and custom scripts. MRI data analysis was performed using the ‘Nilearn’ package, v0.10.4 (Abraham et al., 2014). Statistical analyses were performed based on functions from the ‘scipy’ library (Virtanen et al., 2020). We also used Nilearn to create brain-related visualizations, while other figures are generated using the ‘matplotlib’ library (Hunter, 2007).

#### Factor analysis for behavioral scores

Each child in our dataset had scores from eleven behavioral measures reflecting skills related to reading tests. To evaluate distinct aspects of reading ability, we summarized these measures using factor analysis to identify maximally independent factors. This analysis was performed using the ‘factor_analyzer’ library (Biggs, 2019), which identifies underlying factors that explain the maximum variance among the behavioral measures. We extracted eigenvalues for these factors – which represent significant variance explained by each factor –, keeping only those factors with eigenvalues greater than one according to Kaiser’s criterion. A subsequent factor analysis with three components and Varimax rotation helped in obtaining clear factor loadings. The three factors, henceforth called “Reading”, “Verbal”, and “Naming”, were based on their highest loadings, indicating the primary skill that each factor represents. Summary scores for each individual were calculated by projecting their behavioral measures onto the factor loadings. We also conducted the Bartlett’s and Kaiser-Meyer-Olkin tests to confirm the suitability of our data for factor analysis. Pearson’s correlation between each pair of individual behavioral measures was also calculated for the construction of the behavioral correlation matrix.

#### MRI data preprocessing

Results included in this manuscript come from preprocessing performed using fMRIPrep 23.1.0 (Esteban et al., 2019, 2018; RRID:SCR_016216), which is based on Nipype 1.8.6 (RRID:SCR_002502).

*<u>Anatomical data preprocessing.</u>* The T1-weighted (T1w) image was corrected for intensity non-uniformity with N4BiasFieldCorrection (Tustison et al., 2010), distributed with antsRegistration (ANTs) (Avants et al., 2008; RRID:SCR_004757), and used as T1w-reference throughout the workflow. The T1w-reference was then skull-stripped with a Nipype implementation of the antsBrainExtraction.sh workflow (from ANTs), using OASIS30ANTs as target template. Brain tissue segmentation of cerebrospinal fluid, white-matter and gray-matter was performed on the brain-extracted T1w using fast (FSL, Zhang et al., 2001; RRID:SCR_002823). Brain surfaces were reconstructed using recon-all (FreeSurfer 7.3.2, Dale et al., 1999; RRID:SCR_001847), and the brain mask estimated previously was refined with a custom variation of the method to reconcile ANTs-derived and FreeSurfer-derived segmentations of the cortical gray-matter of Mindboggle (Klein et al., 2017; RRID:SCR_002438). We counted the number of voxels within this mask and used this number as a measure of subject-specific total intracranial volume. Volume-based spatial normalization to a standard pediatric space (MNIPediatricAsym:cohort-3) was performed through nonlinear registration with ANTs, using brain-extracted versions of both T1w reference and the T1w template. The following template was selected for spatial normalization and accessed with TemplateFlow (23.0.0, Ciric et al., 2022): Montreal Neurological Institute (MNI)’s unbiased standard MRI template for pediatric data from the 4.5 to 18.5y age range [RRID:SCR_008796; TemplateFlow ID: MNIPediatricAsym:cohort-3].

*<u>Functional data preprocessing.</u>*For each of the Blood Oxygen Level-Dependent (BOLD) runs available per subject (across all tasks and sessions), the following preprocessing was performed. First, a reference volume and its skull-stripped version were generated using a custom methodology of fMRIPrep. Head-motion parameters with respect to the BOLD reference (six rotation and translation parameters) are estimated before any spatiotemporal filtering using mcflirt (FSL, Jenkinson et al., 2002). BOLD runs were slice-time corrected to the middle slice using 3dTshift from AFNI (Cox and Hyde, 1997; RRID:SCR_005927). The BOLD time-series (including slice-timing correction) were resampled onto their original, native space by applying the transforms to correct for head-motion. The BOLD reference was then co-registered to the T1w reference using bbregister (FreeSurfer), which implements boundary-based rigid registration (Greve and Fischl, 2009), with six degrees of freedom. The confound time series was derived from head motion estimates and other parameters that are included by default in fMRIprep processing (Satterthwaite et al., 2013). The BOLD time-series were resampled into standard space, generating a preprocessed BOLD run in MNIPediatricAsym:cohort-3 space. A reference volume and its skull-stripped version were generated using a custom methodology of fMRIPrep. All resamplings can be performed with a single interpolation step by composing all the pertinent transformations (i.e., head-motion transform matrices and co-registrations to anatomical and output spaces). Gridded (volumetric) resamplings were performed using antsApplyTransforms (ANTs), configured with Lanczos interpolation to minimize the smoothing effects of other kernels (Lanczos, 1964). For information on head motion criteria see below. Finally, we spatially smoothed the resulting functional images with a Gaussian kernel of 6-mm³ full-width at half maximum (FWHM).

#### Estimation of individual contrast maps

We used the preprocessed functional images to estimate activation contrasts for each individual and task, creating individual contrast maps for later use in the prediction step. Condition-related activation was estimated using general linear models (GLM) with functions implemented in nilearn. The task-specific design matrices included regressors representing the following conditions: (i) “ff”, “pseudohomophone”, “pseudoword”, and “word” for the PhonLex task; (ii) “correct” and “incorrect” trials of audiovisual presentation for the Learn task; (iii) “word” and “face” for the Localizer task; and (iv) “letter”, “FFtrained”, “FFfamiliar”, and “FFnew” for the CharProc task. For the Localizer and CharProc tasks, the regressors of interest were constructed with boxcar functions with the onset and the durations of the visual presentations of each condition for each run and participant. For the PhonLex and Learn tasks, these regressors of interest were constructed with the onset and the run-specific mean reaction time as durations for each run and participant; trials with responses given earlier than 0.1 s for both tasks, or longer than 3.5 s for the PhonLex and 2.5 for the Learn task were not modeled. In addition, regressors of no-interest representing button presses (using the recorded times as onsets and zero duration) were also included in the design matrices of all tasks. Task-specific regressors of no-interest represented other events, i.e., feedback presentation in the Learn task, and rocket presentation in the Localizer and CharProc tasks. For the PhonLex and Learn tasks, we adjusted the beta estimations using parametric modulators constructed using the demeaned reaction time for each trial, as described by Mumford et al. (Mumford et al., 2024, 2015). The design matrix also included regressors of no-interest for the six head motion parameters and for the volumes flagged as motion outliers (see the flagging procedure in the next paragraph). After convolving all regressors with a hemodynamic response function designed in SPM (Statistical Parametric Mapping − www.fil.ion.ucl.ac.uk), we estimated the GLM betas for each regressor simultaneously for all available runs per participant. A high-pass filter with a cosine drift model and a cut frequency of 1/128 Hz, alongside a first-degree auto-regressive model, were used to remove low-frequency fluctuations and temporal autocorrelation in the time-series. We then computed participant– and task-specific contrast maps using these betas. The contrasts could be either simple or subtractive, i.e., simple contrasts were comprised of a single estimated beta representing a condition relative to baseline, and subtractive contrasts were comprised of a subtraction of betas, representing the difference between two conditions. All possible subtractions between conditions within the same task were computed, but the subtraction of two given conditions was not computed twice (i.e., given conditions “A” and “B”, only the subtraction “A” minus “B” was computed, but not “B” minus “A”).

In our study, volumes were flagged as outliers based on the Euclidean distance (ED), calculated from translation and rotation parameters of head motion, and the head radius defined as 65 mm. The ED-based threshold for flagging outlier volumes was set to 1.5. Volumes exceeding this threshold were used to form regressors of no-interest in the GLM estimation. Runs with over 10% of volumes flagged as outliers were excluded from the task-specific predictive model. Due to excessive head motion, three runs from the PhonLex task and fifteen from the Learn task were removed. Additional criteria for run removal included the absence of hits for either one of the conditions of interest or number of responses to an attentional indicator (i.e., rocket presentation) below a given threshold, here set at five for the Localizer task and fifteen for the CharProc task. Consequently, further two runs in the PhonLex task, six in the Localizer task, and twelve in the CharProc task were also removed. Therefore, because of available runs and run-specific exclusion criteria, the number of subjects varied depending on the task. More specifically, we analyzed data from 93 children for the PhonLex task (8 with two runs, and 85 with one run), 97 for the Learn task (13 with four runs, 18 with three runs, 63 with two runs, and 3 with one run), 80 for the Localizer task (10 with two runs, and 70 with one run), and 73 for the CharProc task (all with one run). Although the number of volumes within some runs varied slightly, all runs lasted until the end of the presentations during the functional MRI acquisition.

#### Predictive modeling

We aimed to predict each child’s phenotype (summary and individual scores) from each of the fMRI task contrasts, evaluate the prediction performances, and understand the brain correlates of the most prominent predictions. To this mean, we used first-level (individual) contrast t-value maps to predict phenotypes, including the eleven individual test scores and three summary scores. Prediction was performed for each combination of either simple or subtractive contrasts (10 PhonLex + 3 Learn + 3 Localizer + 10 CharProc = 26 contrasts) and individual test scores or summary scores (11 individual scores + 3 summary scores = 14 phenotypes), resulting in a total of 364 predictions. The number of subjects available for each prediction varied depending on MRI data availability for the experimental subjects and the applied exclusion criteria. The predictive modeling approach was based on the Brain Basis Set model, as described in previous studies (Kessler et al., 2016; Sripada et al., 2020, 2019). This model was applied separately to each pair of phenotype and task contrast. The predictive modeling was performed based on ‘scikit-learn’ (Pedregosa et al., 2012).

First, we extracted voxel values from the contrast maps across subjects using a mask common to all participants. For each subject, the three-dimensional data was converted into a one-dimensional vector. The dataset was then split into 10 folds, with nine folds used for training and one fold for testing, constituting a 10-fold cross-validation. Any remainder from the division by 10 was randomly distributed across the folds without repetition. Subjects were randomly assigned to the folds; however, siblings were always placed in the same fold to prevent family-related bias in the model, ensuring they were not separated into training and test sets.

We then constructed the feature matrices *M_train_* and *M_test_*, for the training and test sets, respectively, where rows represented subjects and columns represented voxels. PCA was performed on *M_train_*, resulting in the coefficient matrix *Coeff*, where rows corresponded to components (with the number of components equal to the number of training subjects minus one) and columns corresponded to voxels. Next, we computed the mean of *M_train_* across subjects and subtract it from both *M_train_* and *M_test_*, yielding mean-centered matrices *M’_train_* and *M’_test_*. We then calculated the inverse of *Coeff* and projected it onto *M’_train_* and *M’_test_* using the dot product. From the projections, we selected features corresponding to the first 10 principal components. The resulting matrices were denoted as *A’_train_* and *A’_test_*, respectively.

We included several covariates in our model to ensure that phenotype prediction was driven by brain features rather than individual characteristics. Specifically, we included intracranial volume, gender, age (at MRI data acquisition), age squared, handedness. Additionally, we incorporated the mean ED across scans and its squared value as covariates. For subjects with multiple runs available for estimating contrast maps for a given task, we used the average ED across runs and its squared average as covariates. These covariates were structured as the columns of *Cov_train_* and *Cov_test_*, with rows representing training and test subjects, respectively. Next, we constructed the matrices *X_train_* and *X_test_*, which include the intercept, the main projections matrices *A’_train_* and *M’_test_*, and the centered covariate matrices 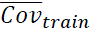 and 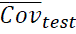, where mean covariate values of training subjects were subtracted. We then estimated a linear multiple regression, modeling the 14 phenotype values for the training subjects as a function of *X_train_* and obtained the beta coefficients. Next, we calculated the dot product of *X_test_* and the beta coefficients from the training subjects to predict the phenotype values for the testing subjects.

Following the 10-fold cross-validation, the prediction of each phenotype and contrast was repeated for all folds. Therefore, after this process we obtained the predicted values of a given phenotype for each subject and contrast. For each combination of contrast map and phenotype, we repeated the process over 500 iterations. For each iteration, we computed the Pearson’s correlation (r), the coefficient of determination (R²), and the mean squared error (MSE) between predicted and actual phenotypes across subjects. The average values of r, R², and MSE across iterations were used as measures of performance.

We compared the predictive performance across tasks and between simple or subtractive contrasts. Here, a simple contrast involves a single condition, whereas a subtractive contrast results from the subtraction of two conditions. Each iteration produced a correlation value between predicted and actual phenotype values, generating a distribution of correlation values. However, the computation of t-tests depends on the sample size, or in this case, the number of iterations. Consequently, we used Cohen’s effect sizes, which are independent of sample size, to investigate differences between tasks or contrast types. Accordingly, we calculated Cohen’s d for each combination of tasks and for simple versus subtractive contrasts within each task and a large difference was noted if d > 0.8.

#### Consensus predictive maps

For each combination of task contrast and summary score, we generated the so-called consensus predictive maps (Sripada et al., 2020), highlighting the brain regions whose heterogeneity significantly contributes to the predicted phenotypes. We first multiplied the *Coeff* matrix by the corresponding beta coefficients for each fold and phenotype, which resulted in a voxel x component matrix. Next, we summed the component values for each fold and phenotype. We then averaged these values across folds for each phenotype and iteration. Finally, we computed the voxel-wise average values across all iterations, and z-score normalized the map for each phenotype. For visual clarity, we adjusted the thresholds on these maps: z = 1.5 in Fig. 5, and z = 2 for cluster labeling in Figs. S9-S11 and Tables S1-S9 of the Supplementary Material. This higher threshold helps prevent the formation of overly large clusters, facilitating clearer labeling.

#### Associating consensus predictive maps with cognitive functions

In an exploratory analysis, to establish associations between the consensus predictive maps and the cognitive significance of the findings, we computed the spatial Pearson’s correlation between unthresholded predictive maps and 400 meta-analytic maps derived from the Neurosynth database (Yarkoni et al., 2011) using the package ‘nimare’. These meta-analytic maps were obtained with the NiMARE library, which applies the Latent Dirichlet Allocation method to analyze the abstracts or texts of articles in Neurosynth (Poldrack et al., 2012). We ranked and reported the 10 term sets from the meta-analytic maps with the highest positive correlations and 10 with the highest negative correlations. Negative correlations in simple contrasts typically reflect deactivations, whereas in subtractive contrasts, such as “condition A minus B”, they indicate that condition B has higher activation than condition A. By ranking these term sets, we aimed to identify the key neural functional aspects associated with the consensus predictive maps.

#### Assessing influence of reading-related brain regions in the predictive model

In another exploratory analysis, we investigated whether prediction performance was primarily driven by activation in individual pre-defined reading-related brain regions (univariate), as opposed to the combination of multiple regions (multivariate). For this analysis, we selected 18 regions-of-interest (ROIs), based on two sources: (1) a meta-analysis of reading-related brain regions using fMRI data (visual word form area, left and right precentral gyrus, left and right supplementary motor area, left and right inferior frontal cortex; Martin et al., 2015) and (2) meta-analytic maps of Neurosynth, generated using specific brain region terms as queries (left posterior insula, right lateral occipital cortex, left and right supramarginal gyrus, medial prefrontal cortex, posterior cingulate cortex, left and right angular gyrus, left and right superior parietal lobule, left superior temporal gyrus; Yarkoni et al., 2011). From Martin et al.’s study (2015), we chose the regions with the highest meta-analytic effect sizes, excluding the cerebellum due to partial coverage in our dataset. From Neurosynth meta-analytic maps, we first spatially smoothed the maps using a Gaussian kernel of 6-mm³ FWHM and then identified the peaks of intensity for the regions specified. We then created 5-mm-radius spherical ROIs around the MNI coordinates of these regions’ peaks using the MarsBaR toolbox (marsbar.sourceforge.net), implemented in MATLAB (www.mathworks.com). We then extracted the average consensus value across voxels within each ROI – referred here as ROI values – for all consensus predictive maps across tasks and contrasts. When an ROI value was negative, we converted it to its absolute value, acknowledging that both increases and decreases in voxel activity could influence predictions. Then, we computed the association between performance and ROI-related consensus values for each of the three predicted summary scores using linear mixed models using R (library ‘lme4’). The consensus values were nested in tasks. We adjusted the significance level using the Bonferroni method to account for multiple comparisons, i.e., 0.05/3 = 0.017. Any adjusted p-value below this level would indicate that the ROI consensus value is significantly associated with the predictive performance of the corresponding summary score.

## 3. Results

### Building the summary scores

Factor analysis using eleven subject-specific behavioral measures resulted in three factors (based on Kaiser’s criterion, as discussed in the Section “Data analysis – Factor analysis of behavioral scores”), which we labeled as “Reading”, “Verbal”, and “Naming” according to the highest factor loadings of each behavioral measure (Fig. 2A). Based on the factor loadings, the Reading summary score primarily reflects the child’s reading fluency and comprehension, as well as spelling ability; the Verbal summary score primarily captures vocabulary knowledge and verbal intelligence; and the Naming summary score predominantly represents the ability to rapidly name objects. Bartlett’s test of Sphericity yielded a p-value of < .0001, and the Kaiser-Meyer-Olkin test resulted in an estimate of .85, confirming that the dataset is suitable and with a strong level of common variance for factor analysis. These three summary scores explain 76.5% of the cumulative proportion of variance present in all the eleven behavioral measures. We also display the correlation matrix of individual behavioral measures based on Pearson’s correlations (Fig. 2B) to show how the variables are associated with each other.

**Figure 2.**
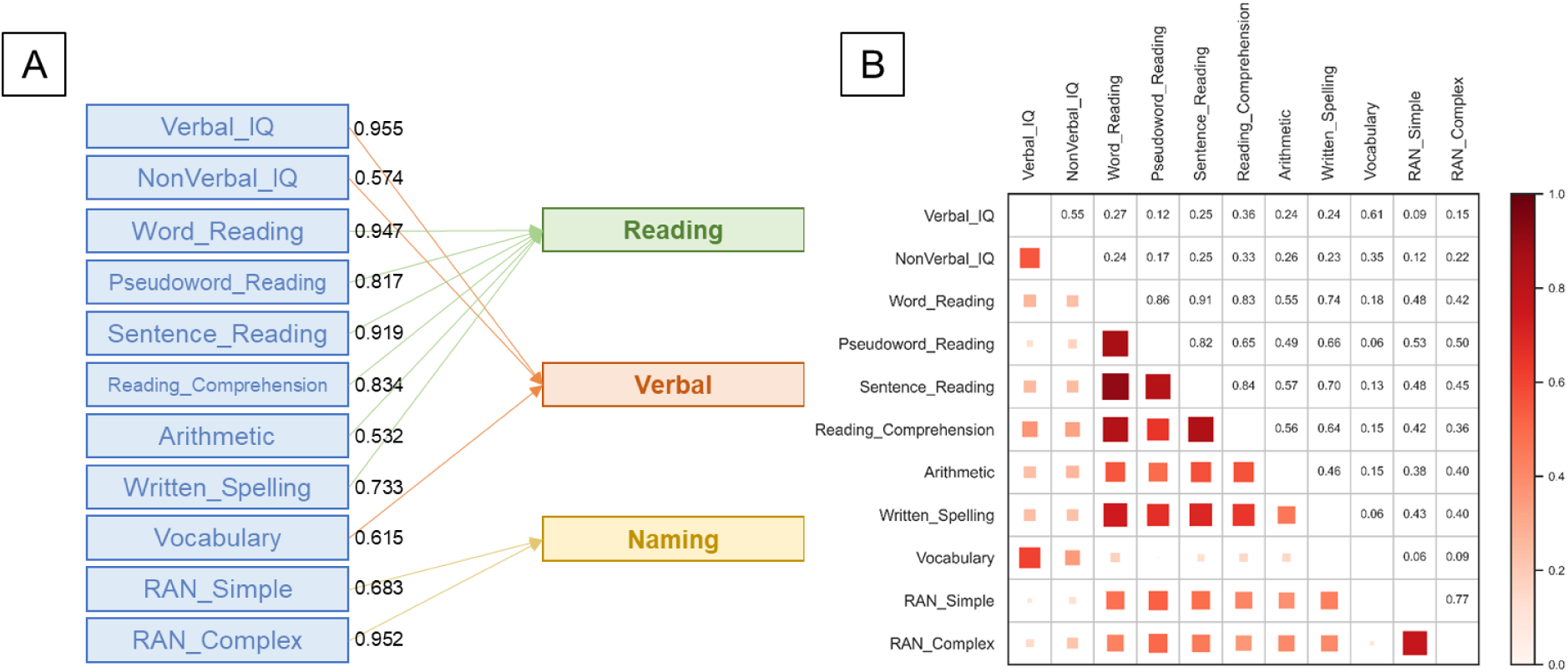
(A) Factor loadings of each normalized behavioral measure for the construction of the factors Reading, Verbal, and Naming, which represent summarizing aspects of reading and cognitive skills, given our available data. Only factor loadings above 0.4 are shown in this plot. (B) Correlation matrix based on Pearson’s correlation across behavioral measures. The strength of the correlation is represented by the squares’ area, as well as the shade of red. RAN = Rapid Automatized Naming, IQ = intelligence quotient.

### Prediction of reading skills from task-based fMRI data

Pearson correlation values between predicted and actual phenotypes, averaged across permutations, are shown for all combinations of task contrasts and phenotypes, including those derived from individual tests and summary scores (Fig. 3). Additionally, R² for each prediction and the MSE between predicted and actual phenotypes, both averaged across iterations, are provided in the Supplementary Material (Figs. S1 and S2). The prediction values for each summary score are also presented as error bar plots, together with scatterplots of predicted vs actual values of the top significant predictions (Fig. 4). Overall, the results indicate that all contrasts from the PhonLex task predict all summary scores; contrasts from the Learn task predict the Reading and Verbal scores, and the Naming score specifically with the contrast “incorrect”; the contrasts “word–face” from the Localizer task predict the Reading score; and almost all simple contrasts from the CharProc task predict the Reading and Naming scores. We also report error bar plots for individual behavioral measures (Figs. S3–S8).

**Figure 3.**
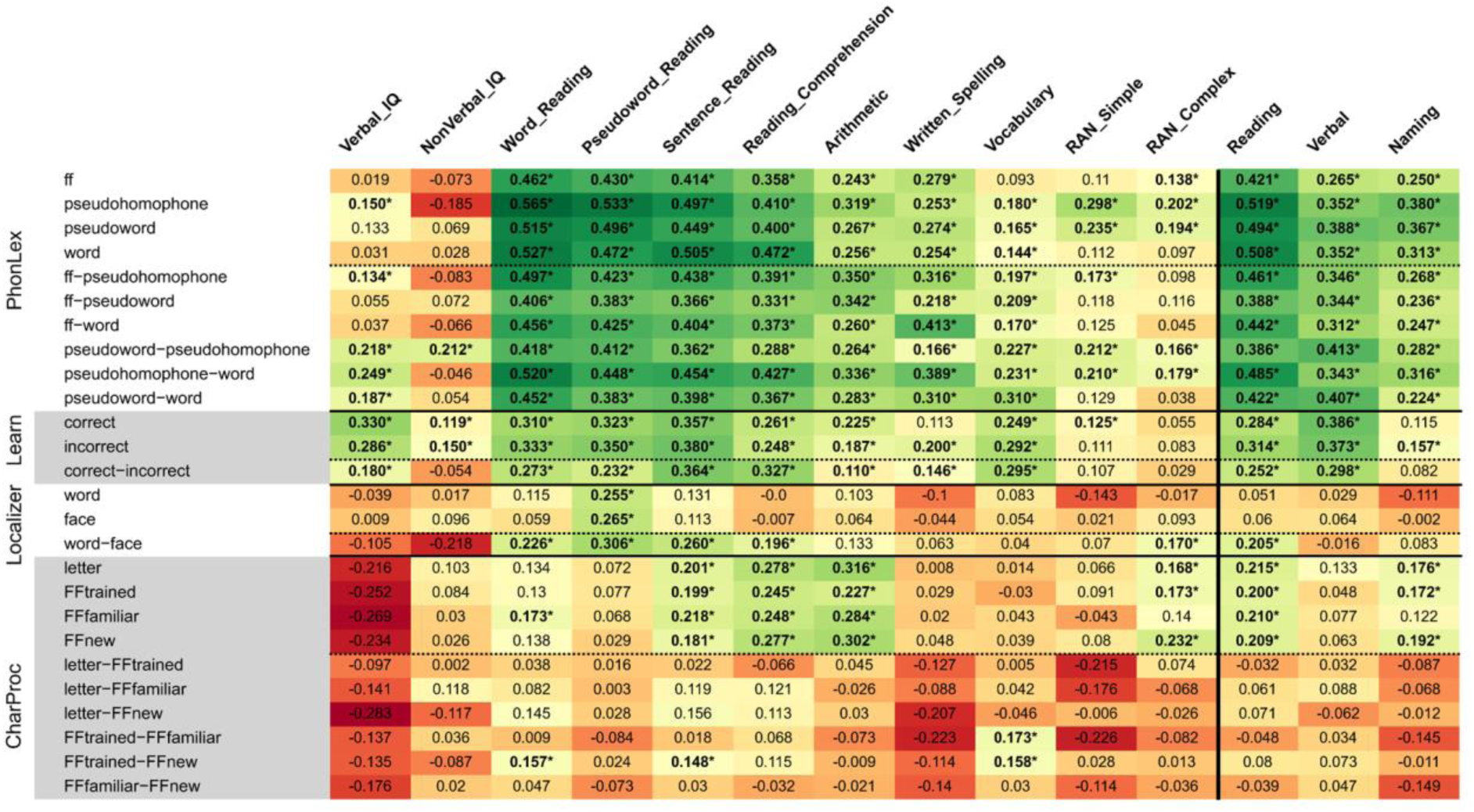
Pearson correlation values between predicted and actual phenotypes for each contrast and task, averaged across iterations. These values of correlation are displayed for individual tests on the left and for the summary scores (Reading, Verbal, and Naming) on the right. Green and red colors indicate stronger positive and negative correlation values, respectively. Asterisks denote correlations that are significantly greater than zero, based on one-sample t-tests corrected for multiple comparisons at the phenotype level of each phenotype. RAN = Rapid Automatized Naming, IQ = intelligence quotient. The solid lines separate results from different tasks, as well as individual behavioral scores and summary scores. The dashed lines separate results from simple and subtractive contrasts.

**Figure 4.**
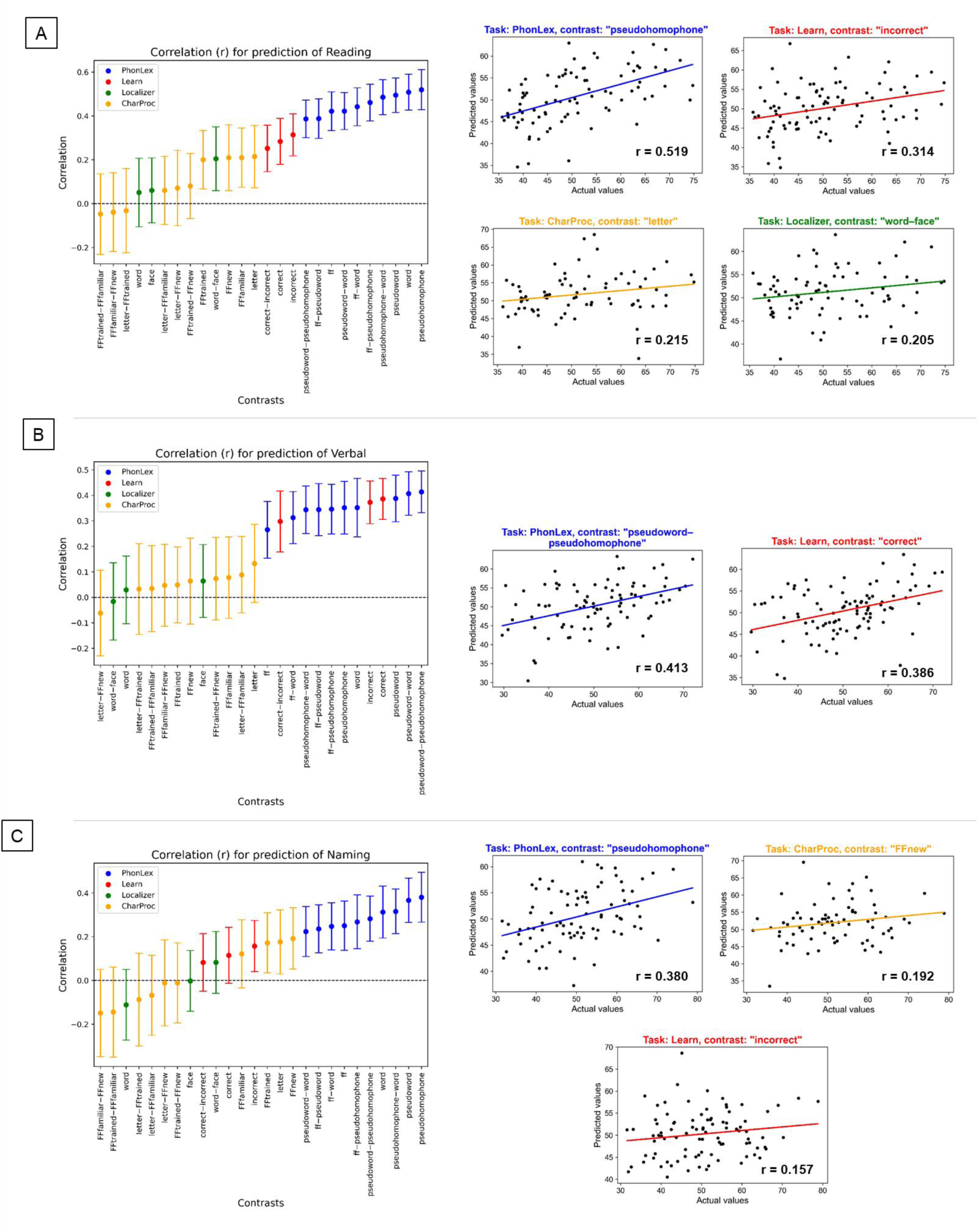
Error bar plots on the left side depict the Pearson correlation between predicted and actual summary scores for (A) Reading, (B) Verbal, and (C) Naming. Dots represent the average correlation across iterations, while the error bars show the adjusted confidence interval for each distribution. Blue, red, green, and yellow error bars correspond to the correlations for the contrasts of the tasks PhonLex, Learn, Localizer, and CharProc, respectively. On the right, we report scatterplots of predicted vs actual values of summary scores. Scatterplots are shown only for the top predictive contrasts of each task and when they are statistically significant. Because of multiple iterations, the plotted scatterplots correspond to the iteration whose correlation is the closest to the mean correlation between predicted and actual values (aggregated across folds) for each combination of task, contrast, and phenotype.

To ensure the analysis was independent of the number of iterations, we compared prediction performances across tasks and contrast types (simple and subtractive) using Cohen’s d effect sizes. We considered differences to be large if d > 0.8. Cohen’s d values for all comparisons are presented in Table 2. For the prediction of the summary scores Reading and Naming, we observed that the performance across tasks followed a consistent pattern: PhonLex > Learn > Localizer = CharProc. For the summary score Verbal, the performance followed a similar pattern, except that no difference was observed between PhonLex and Learn (i.e., PhonLex = Learn > Localizer = CharProc). When comparing contrast types, the performance prediction of the summary scores Reading and Naming was higher for simple compared to subtractive contrasts of the tasks PhonLex, Learn, and CharProc, but the inverse was observed for the Localizer task. For the summary score Verbal, the performance was higher for simple compared to subtractive contrasts of the tasks Learn and CharProc.

**Table 2.** Cohen’s d effect size values for the comparison of prediction performance across tasks and contrast types (simple or subtractive) for the summary scores Reading, Verbal, and Naming. Bold values correspond to large differences (d > 0.8).

|  | Reading | Verbal | Naming |
| --- | --- | --- | --- |
| PhonLex – Learn | **3.55** | 0.00 | **2.99** |
| PhonLex – Localizer | **4.89** | **6.01** | **3.78** |
| PhonLex – CharProc | **3.99** | **4.81** | **2.48** |
| Learn – Localizer | **2.65** | **6.15** | **1.72** |
| Learn – CharProc | **2.18** | **4.78** | **0.94** |
| Localizer – CharProc | 0.12 | -0.44 | -0.25 |
| PhonLex: simple – subtractive contrasts | **1.19** | -0.42 | **1.18** |
| Learn: simple – subtractive contrasts | **1.35** | **2.45** | **1.24** |
| Localizer: simple – subtractive contrasts | **-3.12** | **1.31** | **-2.29** |
| CharProc: simple – subtractive contrasts | **3.02** | 0.68 | **3.46** |

### Consensus predictive maps

We present the surface consensus predictive maps for contrasts of each task concerning the summary scores Reading, Verbal, and Naming (Fig. 5). We only display task contrasts that have the highest correlation between actual and predicted summary scores, provided that there was a significantly positive correlation. These maps are also available in axial views with annotated clusters, provided in the supplementary material (Figs. S9–S11). Specifically, for the PhonLex task, regions such as the fusiform gyrus, insular cortex, inferior frontal gyrus (IFG), inferior temporal gyrus (ITG), lateral occipital cortex (LOC), superior parietal lobule (SPL), supramarginal gyrus (SMG), cuneus, as well as sensorimotor regions (precentral (preCG) and postcentral gyri (postCG)) and key areas of the DMN (precuneus, middle frontal gyrus, and angular gyrus) showed high predictive values for the summary scores (Figs. S9A, S10A, and S11A). For the Learn task, high predictive values were observed for the summary scores in the lingual gyrus, insular cortex, SMG, preCG, postCG, paracingulate gyrus, IFG, ITG, and SPL (Figs. S9B, S10B, S11C). In the Localizer task, the fusiform gyrus, ITG, occipital lobe, superior frontal gyrus, IFG, preCG were particularly predictive for the Reading summary score (Fig. S9D). In the CharProc task, the fusiform gyrus, preCG, cuneus, frontal medial cortex, middle temporal gyrus, LOC, and cuneus were predictive for the Reading and Naming summary scores (Figs. S9C, S11B). A comprehensive list of all clusters identified in these predictive maps is detailed in Tables S1-S9. The main associations with meta-analytic terms of the consensus maps, for positive and negative values, are shown in Tables S10–S12.

**Figure 5.**
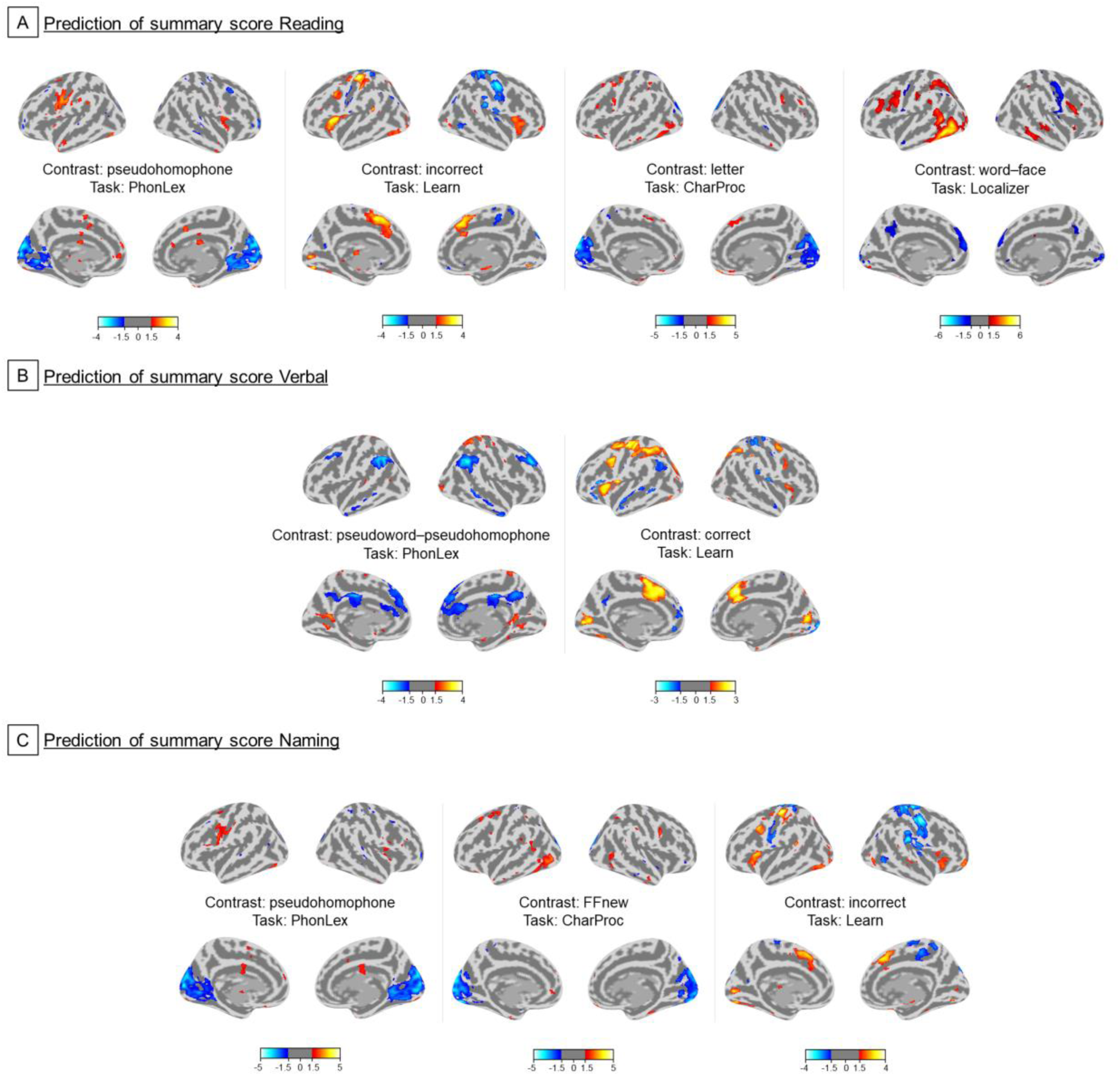
Surface predictive maps of the summary scores (A) Reading, (B) Verbal, and (C) Naming. We present only the predictive maps for the contrast of each task that showed the highest correlation between predicted and actual summary scores, provided that the correlation was significantly positive. The threshold of the displayed consensus maps was set at z = 1.5.

### Association between correlation of prediction and ROI-specific consensus values

In an exploratory analysis, we show that activity (absolute consensus values) in some reading-related ROIs, namely the left inferior frontal gyrus, medial prefrontal cortex, and right angular gyrus, individually contributes to the successful predictive performance (i.e., the average Pearson correlation between predicted and actual phenotype values across iterations) of the Reading summary score (Table 3). Conversely, some regions – the right supramarginal gyrus and the left superior temporal gyrus – negatively influence predictive performance of this summary score. No other significant individual association was observed for the other summary scores.

**Table 3.** Associations between prediction performance (measured by the average Pearson correlation between predicted and actual phenotype values across iterations) and the absolute consensus values within each region-of interest (ROI) for the three summary scores.

| ROI | Reading |  | Verbal |  | Naming |  |
| --- | --- | --- | --- | --- | --- | --- |
|  | t-value | p-value | t-value | p-value | t-value | p-value |
| VWFA | -0.85 | 0.4 | 0.19 | 0.9 | -1.15 | 0.3 |
| L postIns | 0.09 | 0.9 | 0.11 | 0.9 | 0.33 | 0.8 |
| R LOC | 0.37 | 0.7 | -0.21 | 0.8 | 2.02 | 0.11 |
| L SMG | 3.57 | 0.02 | 1.32 | 0.25 | 0.11 | 0.9 |
| R SMG | -4.15 | 0.004* | -0.85 | 0.4 | -0.83 | 0.5 |
| L preCG | 0.28 | 0.8 | -1.05 | 0.4 | 1.59 | 0.18 |
| R preCG | -1.24 | 0.29 | 0.57 | 0.6 | 0.58 | 0.6 |
| L SMA | 1.66 | 0.19 | -0.40 | 0.7 | -1.36 | 0.24 |
| R SMA | 3.02 | 0.19 | -1.11 | 0.3 | -1.54 | 0.19 |
| L IFG | 3.86 | 0.006* | -0.96 | 0.4 | -2.22 | 0.08 |
| R IFG | -0.82 | 0.46 | 0.01 | 1 | 1.06 | 0.3 |
| mPFC | 5.79 | 0.0007* | 0.35 | 0.7 | 0.15 | 0.9 |
| PCC | 3.15 | 0.13 | -1.61 | 0.18 | 0.14 | 0.9 |
| L Ang | -2.32 | 0.15 | -1.68 | 0.17 | -2.64 | 0.05 |
| R Ang | 3.53 | 0.010* | 1.46 | 0.22 | 1.41 | 0.22 |
| L SPL | -0.16 | 0.9 | -0.82 | 0.5 | -0.41 | 0.7 |
| R SPL | 2.95 | 0.06 | 1.50 | 0.21 | 0.26 | 0.8 |
| L STG | -6.95 | 0.0006* | -2.22 | 0.08 | -0.01 | 0.9 |

*Note: Asterisks show statistically significant associations according to a Bonferroni-corrected significance level (0.05/3 = 0.017). Abbreviations: L/R = left/right, VWFA = visual word form area, postIns = posterior insular cortex, LOC = lateral occipital cortex, SMG = supramarginal gyrus, preCG = precentral gyrus, SMA = supplementary motor area, IFG = inferior frontal gyrus, mPFC = medial prefrontal cortex, PCC = posterior cingulate cortex, Ang = angular gyrus, SPL = superior parietal lobule, STG = superior temporal gyrus.*

## 4. Discussion

In this study, we employed task-based fMRI to explore and predict phenotypes representing several dimensions of reading skills in children, aiming to understand the neural basis of literacy development. We constructed summary scores that primarily represented (i) reading fluency, comprehension, and spelling; (ii) vocabulary knowledge and verbal intelligence; and (iii) object naming fluency. We used condition-specific activation maps from four tasks performed during fMRI scanning as predictors: a phonological lexical decision task, an audiovisual association learning task, a visual localizer task of word and face processing, and a character processing task with characters differing in familiarization levels. The models significantly predicted all three summary scores, yielding high effect sizes, and with the highest predictive performance for contrasts from the phonological lexical decision and the audiovisual association learning tasks. Simple contrasts generally yielded higher predictive performances than those involving subtraction of conditions. Successful predictions were supported by activity in brain regions critical for language and reading abilities, including the IFG, SMG, and fusiform gyrus, as well as large-scale functional networks such as salience, dorsal attention, and default mode networks. As suggested by the results of ROI analyses, the predictions in our study may have succeeded primarily due to multivariate, rather than univariate, associations between brain features and phenotypes (Varoquaux and Poldrack, 2019; Woo et al., 2017; Yarkoni and Westfall, 2017).

### Assessment of reading scores

We administered a comprehensive battery of assessments, mostly evaluating reading skills in children, including verbal intelligence, reading fluency, reading comprehension, written spelling, vocabulary knowledge, and the speed and accuracy of naming familiar stimuli. These measures were subsequently grouped into three primary categories: Reading, Verbal, and Naming. The Reading summary score represents a broad evaluation of literacy skills, capturing a child’s efficiency in reading and understanding texts, as well as their ability to accurately interpret and comprehend written language. The Verbal summary score reflects a child’s capacity to understand language, encompassing both linguistic proficiency and verbal reasoning skills. The Naming summary score measures a child’s ability to retrieve and access lexical semantics, reflecting cognitive efficiency in visual recognition and verbal expression. It is important to note that these summary scores are not independent; they share common cognitive processes, such as lexical retrieval, visual scanning, and language representation (Norton and Wolf, 2012). Despite being conceptually distinct, arithmetic skills and reading-related measures (i.e., word and pseudoword reading, sentence reading, reading comprehension, and written spelling) (Fig. 2B) showed moderate correlations. This aligns with previous findings from behavioral, brain and also genetic studies (Baldo and Dronkers, 2007; Balhinez and Shaul, 2019; Davis et al., 2014; Pollack and Ashby, 2018; Prado et al., 2014; Singer and Strasser, 2017), and might reflect general cognitive abilities supporting both reading and arithmetic skills. In contrast, prediction of non-verbal IQ exhibited poor performance (Fig. 3), suggesting that this phenotype may serve as a more suitable control measure for demonstrating the specificity of task-based fMRI predictions to reading-related skills rather than general cognitive ability. The failure of tasks targeting reading-related regions to predict non-verbal IQ thus supports the cognitive selectivity of our approach. Furthermore, previous studies have typically predicted reading skills from unidimensional phenotypes (Kristanto et al., 2020) or from the limited reading-related measures available in the Human Connectome Project, which includes only two tests from which reading ability can be inferred (see Nakai et al., 2024 for a list of related studies using HCP data). In contrast, the present study provides a novel contribution by first conducting a comprehensive assessment of reading skills and then applying prediction modeling to multidimensional literacy measures. The resulting summary scores explained a substantial proportion of variance across the behavioral tasks (76.5%), reflecting strong explanatory power across the behavioral measures.

### Methodological factors influencing performance of literacy traits

In general, many fMRI task contrasts were effective in predicting various aspects of reading skills in our study (Figs. 3 and 4). The contrasts derived from the PhonLex task demonstrated the highest predictive values across all summary scores (Reading: r = .39–.52, Verbal: r = .26–.41, Naming: r = .22–.38) and all its contrasts yielded significant predictions. The contrasts from the Learn task were predictive for the Reading (r = .25–.31) and Verbal (r = .30–.39) summary scores, with only the “incorrect” contrast predicting the Naming summary score (r = .16). All simple contrasts from the CharProc task predicted the Reading summary score (r = .20–.22), and most also predicted the Naming summary score (r = .17–.19). Only the subtractive contrast from the Localizer task predicted a summary score (Reading, r = .20). In sum, the predictive performance of the Reading and Naming summary scores followed the pattern PhonLex > Learn > Localizer = CharProc; for the Verbal summary score, the pattern was PhonLex = Learn > Localizer = CharProc (Table 2). The superior performance of the PhonLex and Learn tasks likely reflects their stronger demands on active, reading-related cognitive processes, as participants must make explicit decisions about letter strings or characters. This is especially true for PhonLex, which requires explicit item reading and a phonological-lexical decision process. The Learn task engages individuals in audiovisual correspondence learning, evoking subsequent evidence accumulation for the decision-making process of associating symbols and sounds. Crucially, this task does not require reading skills yet still demonstrates high predictive performance for reading. This dissociation between task demands and prediction targets suggests the Learn task could serve as a valuable tool for forecasting future reading skills in pre-literate children, offering potential for early identification of individual differences in literacy trajectories (Aravena et al., 2017; Bonte and Brem, 2024; Gellert and Elbro, 2018; Horbach et al., 2018, 2015). Given that brain responses after brief symbol-speech sound training can predict early reading outcomes (Karipidis et al., 2018), future longitudinal studies could test whether fMRI-based learning measures further enhance the prediction of reading outcomes in prereaders. In contrast, the Localizer and CharProc tasks were passive, involving only implicit processing of characters or words, with responses required solely to detect an oddball stimulus. Taken together, these findings indicate that active fMRI tasks outperform passive paradigms in predicting children’s reading skills. Tasks that directly engage core reading operations, such as lexical decisions, show the strongest predictive power, yet even tasks that do not require explicit reading and can therefore be administered to illiterate or prereading children, such as audiovisual correspondence learning, also yield strong predictive power.

The prediction success observed here is consistent with reports that task-based fMRI outperforms resting-state fMRI for phenotype prediction (Greene et al., 2018; Nakai et al., 2024; Sripada et al., 2020; Tomasi and Volkow, 2020). As in treadmill testing in cardiology—where cardiac function is most revealing under load—task paradigms more informatively probe brain function by actively engaging relevant cognitive processes and networks, thereby amplifying individual differences in the systems critical for the ability of interest and suppressing irrelevant activity, which in turn boosts predictive performance (Sripada et al., 2020).

Prediction performance based on simple contrasts generally exceeded that based on subtractive contrasts (Table 2). Simple contrasts tend to yield stronger activation within the reading network, whereas subtractive contrasts can attenuate signal and diminish between-subject variability in key clusters, lowering predictive performance. The notable exception was the Localizer task, where subtraction likely isolated regions specifically involved in word and face processing and showed superior predictive performance for Reading and Naming summary scores compared to simple contrasts. In this case, the subtraction process likely isolated key regions critical for accurately predicting these phenotypes, aligning with the Localizer task’s objective of identifying brain areas specifically involved in word processing (Chauhan et al., 2024; Li et al., 2024; Vinckier et al., 2007).

Performance in tests that loaded most strongly on the Reading summary score – covering word, sentence, and pseudoword reading, reading comprehension, and spelling (Fig. 2A) – was generally well predicted by PhonLex task contrasts, although other contrasts also predicted individual test scores with lower performances (Figs. S4–S6). Learn task contrasts most strongly predicted scores contributing to the Verbal summary score, particularly verbal and non-verbal IQ and vocabulary (Figs. S3 and S7), followed by those from the PhonLex task. The RAN scores, substantially forming the Naming summary score, were well predicted by PhonLex task contrasts, with “complex RAN” also predicted by the contrasts from CharProc and Localizer tasks (Fig. S8). The strongest prediction, using the contrast “pseudohomophone” from the PhonLex, explained up to 26% of the variance of the Reading summary score and 31% of the variance of word reading fluency (Fig. S1). These values align with previous findings on the prediction of cognitive abilities based on fMRI tasks (28%, Sripada et al., 2020).

### Key brain regions influencing performance in predicting literacy traits

By leveraging individual variability in activation patterns, our brain activation maps provide important insights into neural heterogeneity that distinguishes children’s literacy skills. Our ROI analyses revealed that the left IFG contributed significantly to the prediction of the Reading summary score (Table 3), indicating that activation variability in this region was a key predictive factor. Representative IFG clusters also appeared in several consensus maps for Verbal summary scores (Fig. S10). These findings are consistent with previous work identifying the left IFG as an important hub of the language and reading network (Fedorenko et al., 2024; Price, 2012; Turker et al., 2025). Encompassing Broca’s area, the left IFG is involved in semantic processing of word reading (Fedorenko et al., 2011; Sahin et al., 2009), as well as language comprehension, production, and complex higher-order linguistic processes (Bitan et al., 2009; Friederici, 2011; Pollack et al., 2015). Moreover, this region has repeatedly emerged as an important predictor of literacy outcomes in both structural and functional imaging studies (Beyer et al., 2022; Feng et al., 2021; Tomasi and Volkow, 2020) and shows altered activation in individuals with dyslexia (Hancock et al., 2017). Notably, the PhonLex task required lexical decisions on character strings, thereby engaging cognitive processes related to detection and resolution of orthographic, phonological and semantic conflicts. These demands may have induced substantial inter-individual variability in left IFG activation (Figs. S9 and S10), which in turn contributed to the prediction of reading fluency, reading comprehension, and verbal intelligence. A previous study reported that improvements in reading skills were primarily predicted by right prefrontal brain activation and by the white matter integrity of the right superior longitudinal fasciculus, suggesting that these neural networks play a critical role in reading development among children with dyslexia (Hoeft et al., 2011). Interestingly, our findings also implied right hemispheric prefrontal regions in prediction of reading skills (see, for example, Learn task contrasts in Fig. 5). However, these associations were less robust in our study, as ROI analyses of the right inferior frontal gyrus (R IFG) and right precentral gyrus (R preCG) did not yield significant univariate effects. In addition, the supramarginal gyrus (SMG) emerged in several consensus maps (see Figs. S9B and S11C), further aligning our findings with prior literature highlighting the SMG as a predictor of literacy skills (Beyer et al., 2022; He et al., 2013). Its contribution to the prediction of reading fluency suggests that variability in SMG activity is functionally relevant, consistent with reports linking SMG underactivation to dyslexia (Linkersdörfer et al., 2012) and with the inclusion of children at risk for dyslexia in our dataset (see Table 1). Furthermore, this SMG variability may have also influenced the prediction of the Naming summary score, as rapid automatized naming scores have been reported to correlate with SMG activity (Beyer et al., 2022). However, our ROI analysis did not confirm a significant contribution of the SMG to the prediction of the Reading summary score (Table 3). In contrast, its right counterpart and the left STG showed negative contributions to prediction performance (Table 3), the interpretation of which remains unclear in the present study.

The left fusiform gyrus, which includes the VWFA, was also a recurrent region in the consensus maps of high-performance predictors (see, for instance, Fig. S9A, S9C, S9D, and S11A). This finding aligns with previous studies (Bach et al., 2013; Beyer et al., 2022; Zahia et al., 2020) showing that VWFA variability enhances the predictive performance of literacy skills. The fusiform gyrus plays a key role in word reading (Chauhan et al., 2024; Dehaene and Cohen, 2011; Li et al., 2024; McCandliss et al., 2003; White et al., 2023) and underactivation of this region has been reported in children, adolescents and adults with reading impairments (Blau et al., 2010; Brem et al., 2020; Centanni et al., 2019; Pleisch et al., 2019b; Richlan et al., 2011; Romanovska et al., 2021). Interestingly, the Localizer task, which was specifically designed to identify the VWFA, yielded relatively lower predictive performance compared with the other tasks, likely due to its lower cognitive demands and correspondingly reduced engagement of broader regions within the reading network. This aligns with recent findings demonstrating that VWFA activity is modulated by task demands, with words evoking larger responses when attended compared to when ignored (Chauhan et al., 2024). Moreover, work by White and colleagues (White et al., 2023) has shown that the VWFA has inherent selectivity for words, but its functional profile is reshaped by voluntary language processing, with maximal activation occurring during explicit linguistic tasks rather than passive viewing. This observation underscores the importance of employing cognitively engaging tasks that recruit an extended reading-related network when aiming to predict individual differences in literacy skills.

Another perspective on identifying key factors for prediction is to examine other large-scale networks (Beyer et al., 2022; Feng et al., 2021; Tomasi and Volkow, 2020), beyond the canonical reading and language networks alone. This aligns with evidence that language processing is governed by distributed neural systems that vary across individuals (Feng et al., 2021; Ullman, 2004). We found that core regions of the DMN, specifically the medial prefrontal cortex and the right angular gyrus, contributed to the predictive performance of the Reading summary score (Table 3). While the DMN typically deactivates during externally oriented task performance (Andrews-Hanna et al., 2014; Dick et al., 2009; Feng et al., 2015; McDonald et al., 2017; Pamplona et al., 2020; Yarkoni et al., 2008) and activates during internally oriented, self-generated cognitive processes such as episodic retrieval, semantic processing, imagery of novel scenes, and self-reflection (Andrews-Hanna et al., 2014; Harrison et al., 2008; Spreng, 2012), recent evidence demonstrates that the DMN dynamically reconfigures during narrative comprehension, with coupling strength predicting memory of narrative segments (Simony et al., 2016). Moreover, stronger intrinsic connectivity within the DMN is linked to better text processing and text-based memory (Zhang et al., 2019). Given that our active reading task (PhonLex) required both external stimulus processing (decoding written text) and internal semantic elaboration (comprehending meaning), individual differences in DMN activity patterns during this task may reflect variability in how effectively participants balanced attention to external visual input with internal meaning construction for lexical decisions, processes fundamental to skilled reading. The DMN’s contribution to prediction may thus capture individual differences in the capacity to flexibly modulate between externally-driven decoding and internally-driven comprehension processes during reading. The switch between externally– and internally-oriented processes is regulated by the salience network (Seeley et al., 2007). Notably, the insular cortex, a key component of this network, recurrently appeared in our consensus maps of significant predictors (see Figs. S9A, S9B, S10B, S11A, and S11C). The insula has been implicated in multiple aspects of speech, language and reading (Oh et al., 2014; Price, 2012), has extensive connections with frontal language areas, and is implicated in semantic, phonological, and syntactic aspects of language processing (Łuniewska et al., 2019; Wang et al., 2020). Therefore, individual variability in the insular cortex function may reflect differences in the efficiency of switching and balancing activity between these large, partially antagonistic networks, ultimately influencing predictive performance.

### Strengths and future directions

We would like to highlight some of the strengths of our study, as well as potential directions for future research. First, we employed repeated k-fold cross-validation with iterations, which has been demonstrated to be more reliable than the leave-one-out method (Varoquaux et al., 2017). Additionally, we applied PCA, which has the advantage of leveraging information from the entire brain in a data-driven manner for predictive modeling. This approach mitigates overfitting and enhances the generalizability of findings to other studies. Moreover, linear regression also facilitates predictor interpretability, allowing us to make straightforward inferences about brain regions that predominantly contribute to predictions. However, advanced methods capturing complex, non-linear relationships between predictors and phenotypes could enhance prediction while preserving interpretability to advance neuroscientific insights into literacy. Second, while publicly available consortium datasets offer higher sample sizes, their limited language– and reading-related behavioral tests and fMRI-based tasks reduce their specificity for literacy studies. In contrast, our study, conducted at a single center, represents an advancement in literacy prediction by integrating a moderate yet well-characterized sample. We considered data from between 73 to 93 children for prediction (depending on the task), incorporating eleven behavioral tests and four fMRI-based tasks. This comprehensive characterization enhances the specificity and relevance of our findings. Third, our study focused on children with a mean age of 8.7 years (range: 6.7–10.3), making it one of the literacy prediction studies with the youngest sample (refer to Nakai et al., 2024, for a comparison). Our findings suggest that reading skills can be predicted from brain activity in childhood, specifically during the initial stages of schooling, when reading skills are undergoing extensive development. We thus provide insights into how variability in children’s brain activation is linked to literacy skills during a critical developmental window marked by rapid behavioral and neural change, suggesting that emerging neural patterns in this period may carry information predictive of individual differences in reading and reading-related abilities.

### Limitations

Our study also poses some limitations. First, it contains a relatively low number of subjects for prediction studies (close to 100 for some fMRI tasks). A relatively lower number of subjects is commonly reported in task-based fMRI compared to resting-state fMRI in studies predicting reading skills (Nakai et al., 2024) and other cognitive skills (Vieira et al., 2022). A limited number of subjects in predictive modeling studies may limit the generalizability of the results to other datasets, can affect reproducibility, and inflate false-positive results. However, given the standardized data acquisition, extensive behavioral tests and fMRI tasks assessing multiple reading aspects, and the challenges of scanning young children, the sample size remains a significant achievement despite methodological challenges.

Second, our study predicted reading skills using neuroimaging data collected close in time to behavioral testing, limiting its ability to detect future reading difficulties or infer neural causality in reading difficulties. Due to challenges in studying prediction of cognitive skills with young children, longitudinal literacy prediction studies remain scarce (Nakai et al., 2024). To our knowledge, only one study to date has applied task-based fMRI with high-dimensional voxelwise pattern decoding to predict literacy-related scores before formal education (Yu et al., 2020). However, with growing longitudinal research on children nearing kindergarten’s end, early diagnosis of reading difficulties is likely to expand (Chyl et al., 2021).

Third, the number of components to retain after applying the PCA method was chosen a priori based on data constraints. While we acknowledge that this choice is arbitrary, a fixed number was used to simplify comparisons and reduce bias in phenotype predictions across task contrasts. Although this number is lower than in a similar previous study (Sripada et al., 2020), it remains reasonable given the proportion of retained components and sample sizes in both studies. Further analyses could refine this selection by determining the optimal number of components for specific phenotype predictions.

Fourth, subject-specific summary scores were derived from a single factor analysis, performed once rather than in each cross-validation fold. This approach was chosen due to the limited number of subjects per fold and challenges in ensuring consistent summary score definitions and satisfying the Kaiser’s criterion at each step. While some information transfer to predictive modeling was inevitable, we believe it was minimal and the method prioritized methodological simplicity and data interpretability.

Finally, even though our study achieved generally good predictions of reading skills, the strongest prediction explained only 26% of literacy skill variance based on fMRI-based contrasts, indicating that a significant proportion of literacy-related variance remains unexplained. Future research could design task sets that optimize prediction performance based on brain activity, potentially by testing other fMRI-based tasks and integrating multimodal information (structural imaging, electrophysiology, and genetics), which may improve prediction performance (Hammer et al., 2015).

## Conclusions

Here, we used machine learning with whole-brain PCA component maps to predict literacy traits in children from fMRI task contrasts. Individual reading scores were derived from an assessment of eleven behavioral measures, which we summarized into three cognitive domains. Overall, the best predictions reached r = .52 and R² = .26 for reading fluency and comprehension, r = .41 and R² = .14 for verbal intelligence, and r = .38 and R² = .12 for object-naming ability. This study combines a broad reading assessment with a fair sample size from a single center. Across fMRI tasks, predictive performance was generally higher in active tasks – where individuals responded to a stimulus after a decision-making process – than in passive tasks, which involved only stimulus processing and simple target detection (with all comparisons showing large effect sizes). Additionally, we found that maps based on simple contrasts, which tend to yield more significant clusters, yielded better predictive performance than those using subtraction-based contrasts, likely because individual variability in key reading-related regions may be cancelled out in subtraction-based analyses (with effect sizes up to d = 3.5). The interpretability of our predictive model provides insights into the neural factors influencing predictive performance. Notably, activation variability was crucial for distinguishing literacy skills in the left inferior frontal gyrus – involved in language processing, grammar acquisition, and conflict resolution –, left supramarginal gyrus – linked to reading fluency, sensory integration, and reading skills –, and visual word form area – associated with word reading. Variability in large-scale networks, such as the default-mode and salience networks, also played a role in predictive performance, likely reflecting the switch between external attention for reading and internal semantic and imagery processing. Our findings tapping into heterogeneity may also relate to neural differences that support reading acquisition in elementary-school-aged children still learning to read. Altogether, our data-driven study advances the prediction of literacy traits in children based on neuroimaging by highlighting methodological factors that influence predictive performance and offering a unique way to link individual variability in brain activity to reading skills.

## Acknowledgements

We are grateful for the support of S. Coraj, G. Pleisch, Y. Jin Ressel, M. Röthlisberger, R. Füzér, E. Montevecchi, E. Hefti, and C. Schneider in recruitment, study planning and management, behavioral assessments, and MRI recordings. Finally, we thank all families and their children for participating in this study.

## Ethics Approval Statement

The project was approved by the local ethics committee of the Canton of Zurich (BASEC No. 2018-01261) and neighboring Cantons in Switzerland, and data collection was conducted with the written consent of a legal guardian and the oral assent of all participating children.

## Data and Code Availability Statement

All obtained results and scripts used for the data analysis are available on the OSF repository: https://osf.io/q29ga/overview.

## Author Contributions

GSPP: Conceptualization, Methodology, Software, Validation, Formal analysis, Data Curation, Writing – Original Draft, Writing – Review & Editing; SS: Software, Validation, Writing – Review & Editing; BHV: Methodology, Writing – Review & Editing; SVDP: Investigation; Writing – Review & Editing; NF: Investigation, Writing – Review & Editing; CGL: Investigation, Writing – Review & Editing; IIK: Investigation, Writing – Review & Editing; SB: Conceptualization, Validation, Resources, Writing – Review & Editing.

