## Supplementary material for "Predicting children’s literacy from task-based fMRI: Neural heterogeneity and task-dependent performance"

| Task | Contrast | Verbal_IQ | NonVerbal_IQ | Word_Reading | Pseudoword_Reading | Sentence_Reading | Reading_Comprehension | Arithmetic | Written_Spelling | Vocabulary | RAN_Simple | RAN_Complex | Reading | Verbal | Naming |
|---|---|---|---|---|---|---|---|---|---|---|---|---|---|---|---|
| PhonLex | ff | -0.211 | -0.253 | 0.181 | 0.145 | 0.127 | 0.07 | -0.058 | 0.01 | -0.168 | -0.119 | -0.117 | 0.136 | -0.018 | -0.022 |
| | pseudohomophone | -0.08 | -0.305 | 0.313 | 0.276 | 0.233 | 0.144 | 0.044 | -0.015 | -0.112 | 0.055 | -0.022 | 0.26 | 0.082 | 0.125 |
| | pseudoword | -0.093 | -0.139 | 0.247 | 0.229 | 0.166 | 0.123 | 0.002 | 0.013 | -0.092 | -0.003 | -0.04 | 0.227 | 0.121 | 0.104 |
| | word | -0.165 | -0.137 | 0.269 | 0.208 | 0.244 | 0.207 | -0.016 | -0.0 | -0.111 | -0.103 | -0.101 | 0.251 | 0.097 | 0.069 |
| | ff−pseudohomophone | -0.109 | -0.28 | 0.221 | 0.134 | 0.155 | 0.119 | 0.076 | 0.039 | -0.085 | -0.056 | -0.107 | 0.184 | 0.066 | 0.009 |
| | ff−pseudoword | -0.148 | -0.152 | 0.107 | 0.077 | 0.054 | 0.042 | 0.06 | -0.067 | -0.051 | -0.11 | -0.105 | 0.088 | 0.073 | -0.04 |
| | ff−word | -0.18 | -0.265 | 0.165 | 0.132 | 0.097 | 0.077 | -0.046 | 0.135 | -0.098 | -0.12 | -0.19 | 0.148 | 0.027 | -0.039 |
| | pseudoword−pseudohomophone | -0.034 | -0.032 | 0.112 | 0.111 | 0.051 | -0.011 | -0.03 | -0.109 | -0.04 | -0.027 | -0.064 | 0.082 | 0.136 | 0.005 |
| | pseudohomophone−word | -0.013 | -0.217 | 0.255 | 0.171 | 0.179 | 0.156 | 0.062 | 0.122 | -0.019 | -0.032 | -0.062 | 0.214 | 0.067 | 0.045 |
| | pseudoword−word | -0.044 | -0.16 | 0.149 | 0.076 | 0.086 | 0.07 | -0.006 | 0.02 | 0.048 | -0.108 | -0.169 | 0.116 | 0.132 | -0.066 |
| Learn | correct | 0.072 | -0.139 | 0.007 | 0.036 | 0.071 | 0.0 | -0.041 | -0.212 | -0.026 | -0.145 | -0.157 | -0.012 | 0.113 | -0.145 |
| | incorrect | 0.035 | -0.114 | 0.037 | 0.062 | 0.094 | -0.008 | -0.078 | -0.092 | 0.024 | -0.156 | -0.133 | 0.025 | 0.1 | -0.105 |
| | correct−incorrect | -0.043 | -0.25 | -0.036 | -0.061 | 0.078 | 0.057 | -0.216 | -0.201 | 0.029 | -0.134 | -0.202 | -0.055 | 0.042 | -0.187 |
| Localizer | word | -0.26 | -0.305 | -0.132 | -0.01 | -0.107 | -0.229 | -0.174 | -0.333 | -0.217 | -0.362 | -0.34 | -0.197 | -0.296 | -0.369 |
| | face | -0.19 | -0.163 | -0.23 | -0.021 | -0.136 | -0.239 | -0.236 | -0.389 | -0.23 | -0.297 | -0.258 | -0.236 | -0.225 | -0.332 |
| | word−face | -0.269 | -0.486 | -0.031 | 0.047 | 0.025 | -0.037 | -0.083 | -0.275 | -0.224 | -0.21 | -0.117 | -0.044 | -0.262 | -0.164 |
| CharProc | letter | -0.326 | -0.153 | -0.202 | -0.209 | -0.11 | -0.023 | 0.005 | -0.274 | -0.24 | -0.256 | -0.123 | -0.108 | -0.121 | -0.134 |
| | FFtrained | -0.361 | -0.204 | -0.21 | -0.205 | -0.13 | -0.081 | -0.104 | -0.281 | -0.29 | -0.239 | -0.132 | -0.138 | -0.216 | -0.156 |
| | FFfamiliar | -0.325 | -0.213 | -0.147 | -0.182 | -0.091 | -0.054 | -0.026 | -0.266 | -0.222 | -0.323 | -0.127 | -0.097 | -0.147 | -0.159 |
| | FFnew | -0.328 | -0.231 | -0.162 | -0.231 | -0.105 | -0.012 | -0.007 | -0.233 | -0.213 | -0.239 | -0.068 | -0.093 | -0.168 | -0.116 |
| | letter−FFtrained | -0.319 | -0.218 | -0.26 | -0.239 | -0.274 | -0.299 | -0.177 | -0.397 | -0.3 | -0.457 | -0.223 | -0.297 | -0.195 | -0.334 |
| | letter−FFfamiliar | -0.34 | -0.17 | -0.241 | -0.255 | -0.183 | -0.151 | -0.305 | -0.307 | -0.262 | -0.356 | -0.339 | -0.236 | -0.219 | -0.332 |
| | letter−FFnew | -0.37 | -0.339 | -0.167 | -0.224 | -0.157 | -0.154 | -0.224 | -0.388 | -0.305 | -0.239 | -0.261 | -0.217 | -0.269 | -0.267 |
| | FFtrained−FFfamiliar | -0.329 | -0.22 | -0.267 | -0.289 | -0.27 | -0.2 | -0.309 | -0.384 | -0.128 | -0.422 | -0.347 | -0.3 | -0.225 | -0.37 |
| | FFtrained−FFnew | -0.362 | -0.284 | -0.166 | -0.242 | -0.18 | -0.163 | -0.266 | -0.376 | -0.125 | -0.208 | -0.267 | -0.22 | -0.177 | -0.276 |
| | FFfamiliar−FFnew | -0.33 | -0.274 | -0.244 | -0.307 | -0.267 | -0.275 | -0.285 | -0.362 | -0.268 | -0.352 | -0.309 | -0.312 | -0.212 | -0.394 |

*Figure S1. Coefficients of determination of each prediction for each contrast and task, averaged across iterations. These values are displayed for individual tests on the left and for the summary scores (Reading, Verbal, and Naming) on the right. Green and red colors indicate stronger positive and negative coefficients of determination, respectively. RAN = Rapid Automatized Naming, IQ = intelligence quotient. The solid lines separate results from different tasks, as well as individual behavioral scores and summary scores. The dashed lines separate results from simple and composite contrasts.*

| | | Verbal_IQ | NonVerbal_IQ | Word_Reading | Pseudoword_Reading | Sentence_Reading | Reading_Comprehension | Arithmetic | Written_Spelling | Vocabulary | RAN_Simple | RAN_Complex | Reading | Verbal | Naming |
|---|---|---|---|---|---|---|---|---|---|---|---|---|---|---|---|
| PhonLex | ff | 111.27 | 124.69 | 89.84 | 90.85 | 96.2 | 98.1 | 108.11 | 107.27 | 115.48 | 115.62 | 114.44 | 91.11 | 104.97 | 107.19 |
| | pseudohomophone | 99.24 | 129.86 | 75.35 | 76.91 | 84.47 | 90.3 | 97.66 | 109.89 | 109.96 | 97.61 | 104.66 | 78.02 | 94.7 | 91.76 |
| | pseudoword | 100.37 | 113.32 | 82.58 | 81.97 | 91.9 | 92.55 | 101.93 | 106.95 | 107.96 | 103.59 | 106.55 | 81.56 | 90.7 | 93.97 |
| | word | 107.02 | 113.16 | 80.17 | 84.16 | 83.28 | 83.65 | 103.76 | 108.35 | 109.84 | 113.95 | 112.75 | 79.03 | 93.11 | 97.66 |
| | ff−pseudohomophone | 101.89 | 127.36 | 85.37 | 91.99 | 93.13 | 92.94 | 94.41 | 104.14 | 107.25 | 109.12 | 113.35 | 86.08 | 96.32 | 103.94 |
| | ff−pseudoword | 105.43 | 114.58 | 97.9 | 98.06 | 104.26 | 101.09 | 96.0 | 115.63 | 103.89 | 114.63 | 113.19 | 96.22 | 95.56 | 109.01 |
| | ff−word | 108.36 | 125.9 | 91.51 | 92.26 | 99.47 | 97.32 | 106.82 | 93.73 | 108.56 | 115.7 | 121.89 | 89.89 | 100.37 | 108.9 |
| | pseudoword−pseudohomophone | 94.96 | 102.72 | 97.32 | 94.5 | 104.58 | 106.65 | 105.26 | 120.16 | 102.82 | 106.1 | 108.99 | 96.81 | 89.08 | 104.33 |
| | pseudohomophone−word | 93.06 | 121.07 | 81.73 | 88.13 | 90.48 | 89.03 | 95.84 | 95.07 | 100.71 | 106.6 | 108.82 | 82.94 | 96.24 | 100.18 |
| | pseudoword−word | 95.93 | 115.43 | 93.34 | 98.21 | 100.73 | 98.11 | 102.74 | 106.14 | 94.07 | 114.4 | 119.77 | 93.21 | 89.49 | 111.74 |
| Learn | correct | 84.06 | 109.41 | 102.89 | 99.42 | 99.23 | 101.18 | 102.17 | 127.18 | 100.06 | 111.35 | 112.87 | 102.29 | 85.68 | 116.08 |
| | incorrect | 87.4 | 106.94 | 99.83 | 96.73 | 96.72 | 102.02 | 105.8 | 114.61 | 95.18 | 112.41 | 110.54 | 98.51 | 86.97 | 112.02 |
| | correct−incorrect | 94.44 | 120.05 | 107.36 | 109.42 | 98.47 | 95.44 | 119.38 | 126.12 | 94.64 | 110.28 | 117.29 | 106.59 | 92.56 | 120.37 |
| Localizer | word | 99.98 | 124.97 | 121.68 | 108.97 | 120.16 | 128.33 | 115.83 | 145.3 | 110.8 | 130.76 | 124.32 | 122.32 | 112.3 | 137.06 |
| | face | 94.45 | 111.35 | 132.24 | 110.18 | 123.3 | 129.4 | 121.99 | 151.41 | 111.98 | 124.52 | 116.72 | 126.3 | 106.19 | 133.35 |
| | word−face | 100.68 | 142.3 | 110.82 | 102.81 | 105.83 | 108.32 | 106.88 | 138.91 | 111.45 | 116.18 | 103.62 | 106.64 | 109.32 | 116.56 |
| CharProc | letter | 117.42 | 97.76 | 133.81 | 130.34 | 122.56 | 110.51 | 99.6 | 140.22 | 125.55 | 118.15 | 106.43 | 114.57 | 100.54 | 112.25 |
| | FFtrained | 120.52 | 102.01 | 134.59 | 129.92 | 124.86 | 116.8 | 110.58 | 141.03 | 130.6 | 116.49 | 107.31 | 117.68 | 108.99 | 114.48 |
| | FFfamiliar | 117.4 | 102.8 | 127.61 | 127.51 | 120.51 | 113.93 | 102.72 | 139.41 | 123.73 | 124.4 | 106.86 | 113.45 | 102.82 | 114.73 |
| | FFnew | 117.68 | 104.35 | 129.35 | 132.72 | 122.03 | 109.4 | 100.84 | 135.78 | 122.86 | 116.52 | 101.29 | 113.04 | 104.68 | 110.47 |
| | letter−FFtrained | 116.8 | 103.24 | 140.23 | 133.61 | 140.73 | 140.42 | 117.84 | 153.81 | 131.6 | 136.99 | 115.98 | 134.14 | 107.14 | 132.08 |
| | letter−FFfamiliar | 118.73 | 99.15 | 138.09 | 135.38 | 130.62 | 124.4 | 130.72 | 143.94 | 127.77 | 127.55 | 126.9 | 127.82 | 109.27 | 131.88 |
| | letter−FFnew | 121.32 | 113.46 | 129.84 | 131.99 | 127.81 | 124.75 | 122.62 | 152.81 | 132.16 | 116.54 | 119.59 | 125.84 | 113.78 | 125.39 |
| | FFtrained−FFfamiliar | 117.76 | 103.38 | 141.0 | 139.01 | 140.25 | 129.67 | 131.12 | 152.38 | 114.21 | 133.69 | 127.71 | 134.37 | 109.79 | 135.59 |
| | FFtrained−FFnew | 120.69 | 108.81 | 129.8 | 133.93 | 130.3 | 125.67 | 126.8 | 151.48 | 113.91 | 113.63 | 120.09 | 126.15 | 105.53 | 126.28 |
| | FFfamiliar−FFnew | 117.81 | 108.02 | 138.46 | 140.93 | 139.89 | 137.73 | 128.68 | 149.99 | 128.44 | 127.12 | 124.11 | 135.68 | 108.66 | 137.95 |

*Figure S2. Mean squared error values between predicted and actual phenotypes for each contrast and task, averaged across iterations. These values are displayed for individual tests on the left and for the summary scores (Reading, Verbal, and Naming) on the right. Red and green colors indicate stronger positive and negative mean squared error values, respectively. RAN = Rapid Automatized Naming, IQ = intelligence quotient. The solid lines separate results from different tasks, as well as individual behavioral scores and summary scores. The dashed lines separate results from simple and composite contrasts.*

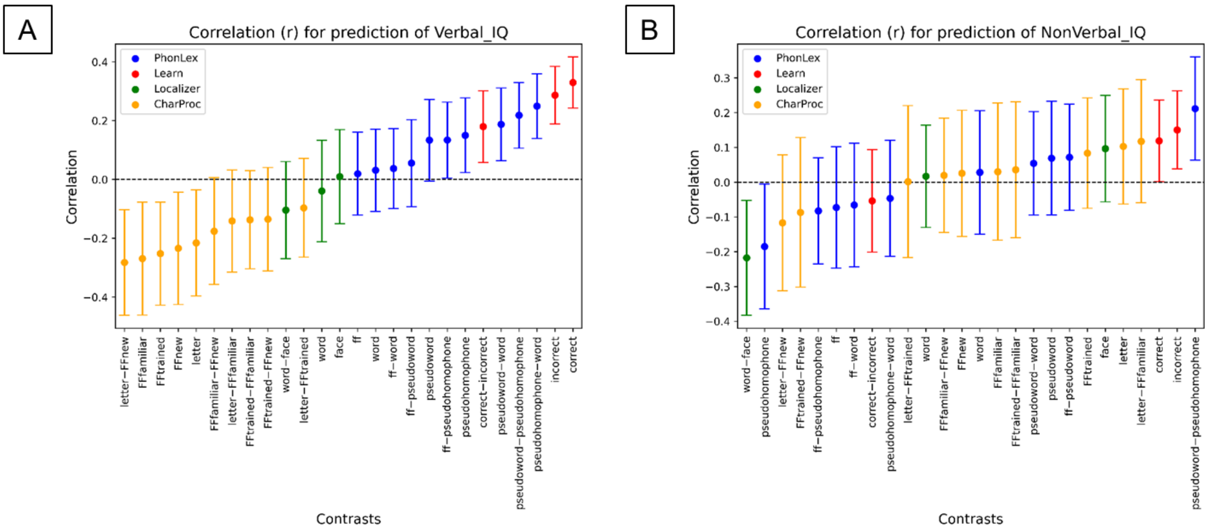


*Figure S3. Error bar plots depicting the Pearson correlation between predicted and actual scores of (A) verbal and (B) non-verbal IQ. Dots represent the average correlation across iterations, while the error bars indicate the adjusted confidence interval for each distribution. Blue, red, green and yellow error bars correspond to the correlations for the constrasts of the tasks PhonLex, Learn, Localizer, and CharProc, respectively.*

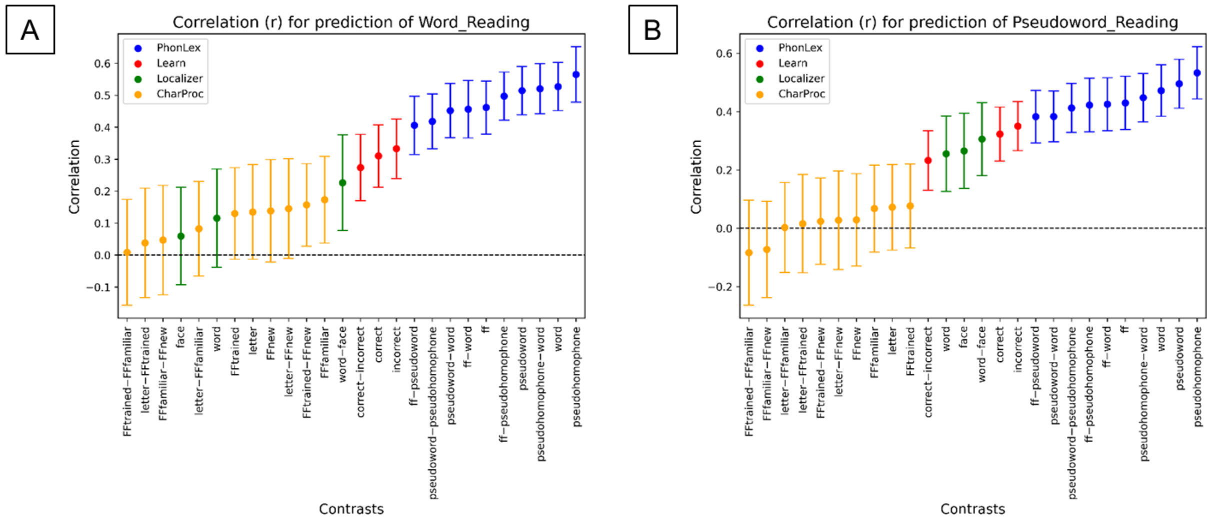


*Figure S4. Error bar plots depicting the Pearson correlation between predicted and actual scores of SLRT (Salzburger Lese- und Rechtschreibtest) (A) for word reading fluency (Word_reading) and (B) pseudoword reading fluency (Pseudoword_reading). Dots represent the average correlation across iterations, while the error bars indicate the adjusted confidence interval for each distribution. Blue, red, green and yellow error bars correspond to the correlations for the constrasts of the tasks PhonLex, Learn, Localizer, and CharProc, respectively.*

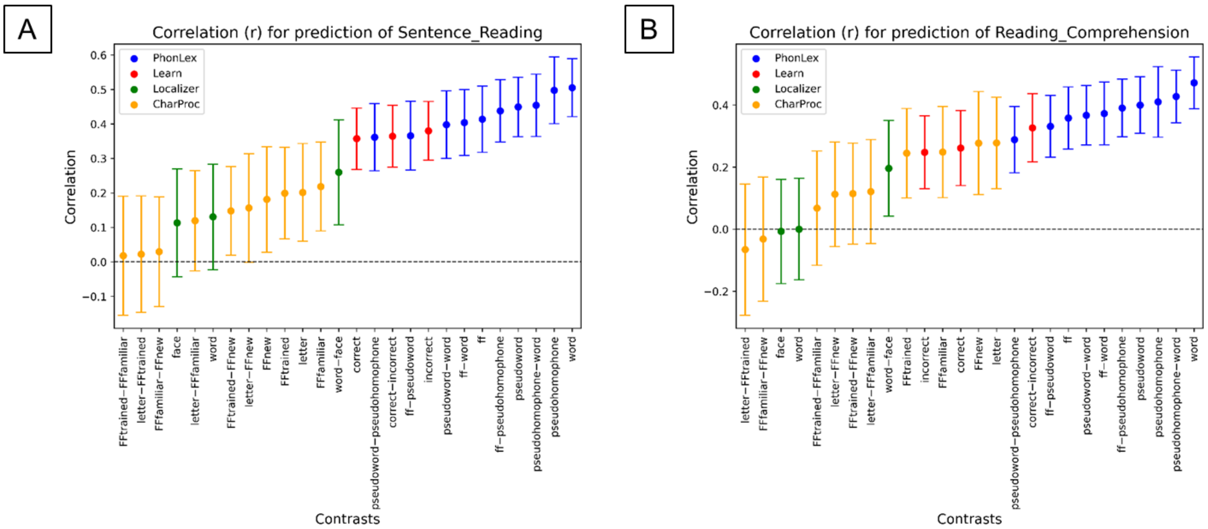


*Figure S5. Error bar plots depicting the Pearson correlation between predicted and actual scores of (A) Sentence_Reading (Salzburger Lese-Screening (SLS)) and (B) Reading_Comprehension (Ein Leseverständnistest für Erst- bis Siebtklässler (ELFE)). Dots represent the average correlation across iterations, while the error bars indicate the adjusted confidence interval for each distribution. Blue, red, green and yellow error bars correspond to the correlations for the constrasts of the tasks PhonLex, Learn, Localizer, and CharProc, respectively.*

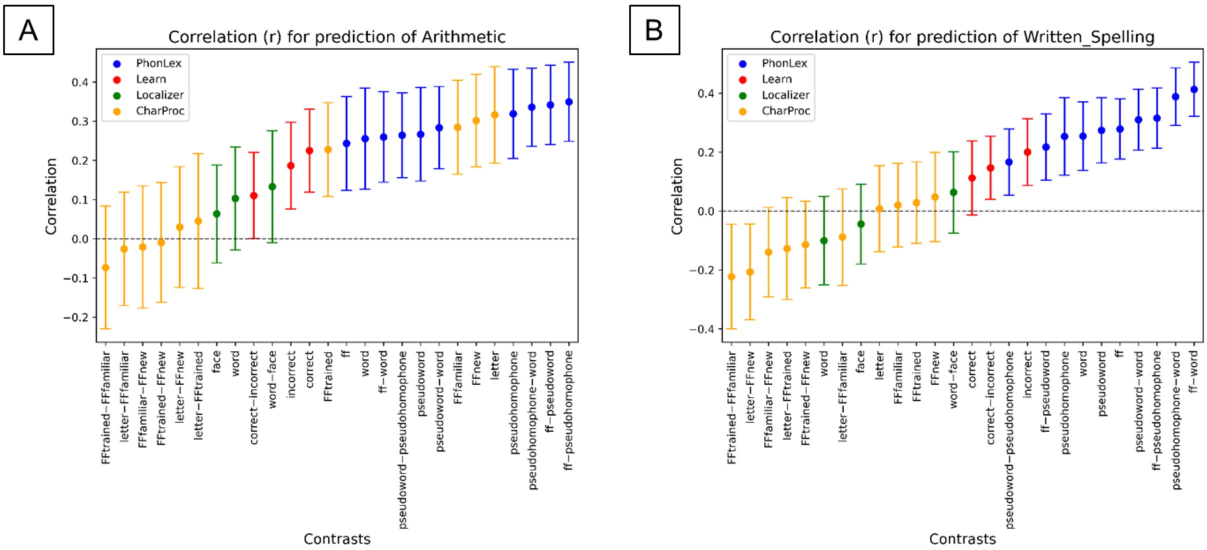


*Figure S6. Error bar plots depicting the Pearson correlation between predicted and actual scores of (A) Arithmetic (Heidelberger Rechentest (HRT)) and (B) Written_Spelling (Schreib.on Test). Dots represent the average correlation across iterations, while the error bars indicate the adjusted confidence interval for each distribution. Blue, red, green and yellow error bars correspond to the correlations for the constrasts of the tasks PhonLex, Learn, Localizer, and CharProc, respectively.*

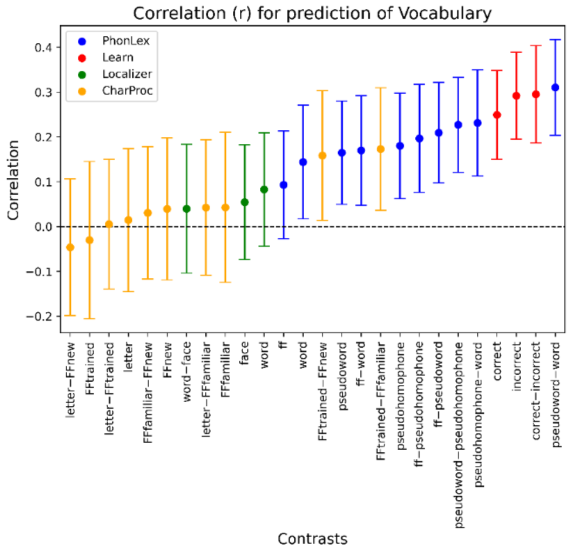


*Figure S7. Error bar plots depicting the Pearson correlation between predicted and actual scores of vocabulary knowledge assessed with the Peabody Picture Vocabulary Test. Dots represent the average correlation across iterations, while the error bars indicate the adjusted confidence interval for each distribution. Blue, red, green and yellow error bars correspond to the correlations for the constrasts of the tasks PhonLex, Learn, Localizer, and CharProc, respectively.*

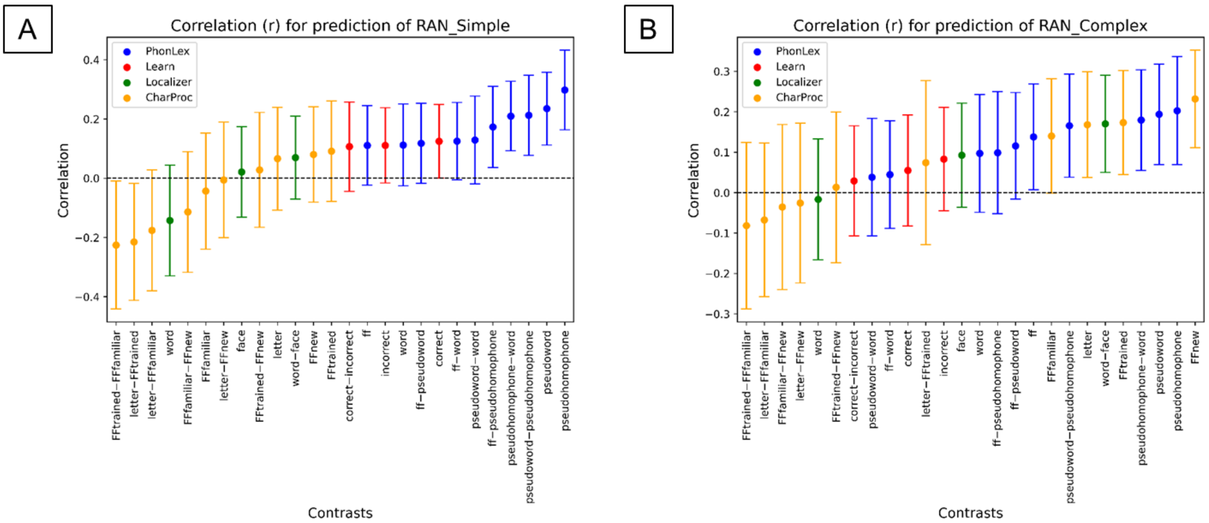


*Figure S8. Error bar plots depicting the Pearson correlation between predicted and actual scores of RAN (Rapid Automatized Naming) for (A) simple and (B) complex words. Dots represent the average correlation across iterations, while the error bars indicate the adjusted confidence interval for each distribution. Blue, red, green and yellow error bars correspond to the correlations for the constrasts of the tasks PhonLex, Learn, Localizer, and CharProc, respectively.*

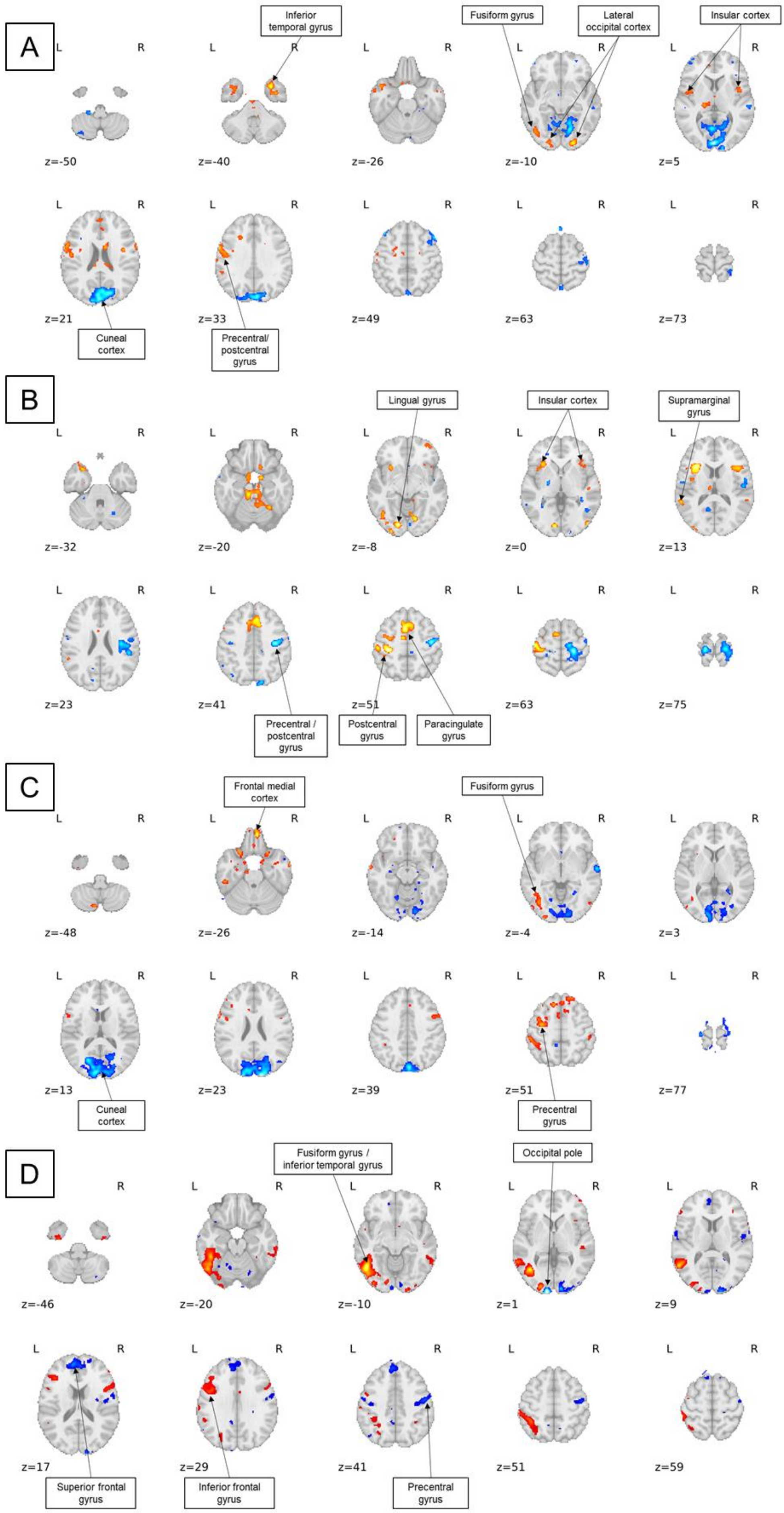
A
Inferior temporal gyrus
Fusiform gyrus
Lateral occipital cortex
Insular cortex
z=-50
z=-40
z=-26
z=-10
z=5
z=21
z=33
z=49
z=63
z=73
Cuneal cortex
Precentral/ postcentral gyrus
B
Lingual gyrus
Insular cortex
Supramarginal gyrus
z=-32
z=-20
z=-8
z=0
z=13
z=23
z=41
z=51
z=63
z=75
Precentral / postcentral gyrus
Postcentral gyrus
Paracingulate gyrus
C
Frontal medial cortex
Fusiform gyrus
z=-48
z=-26
z=-14
z=-4
z=3
z=13
z=23
z=39
z=51
z=77
Cuneal cortex
Precentral gyrus
D
Fusiform gyrus / inferior temporal gyrus
Occipital pole
z=-46
z=-20
z=-10
z=1
z=9
z=17
z=29
z=41
z=51
z=59
Superior frontal gyrus
Inferior frontal gyrus
Precentral gyrus

*Figure S9. Axial slices of predictive maps of the summary measure Reading for the contrasts (A) pseudohomophone (from PhonLex), (B) incorrect (from Learn), (C) letter (from CharProc), and (D) word–face (from Localizer). The threshold of the displayed consensus maps was set at z = 2.*

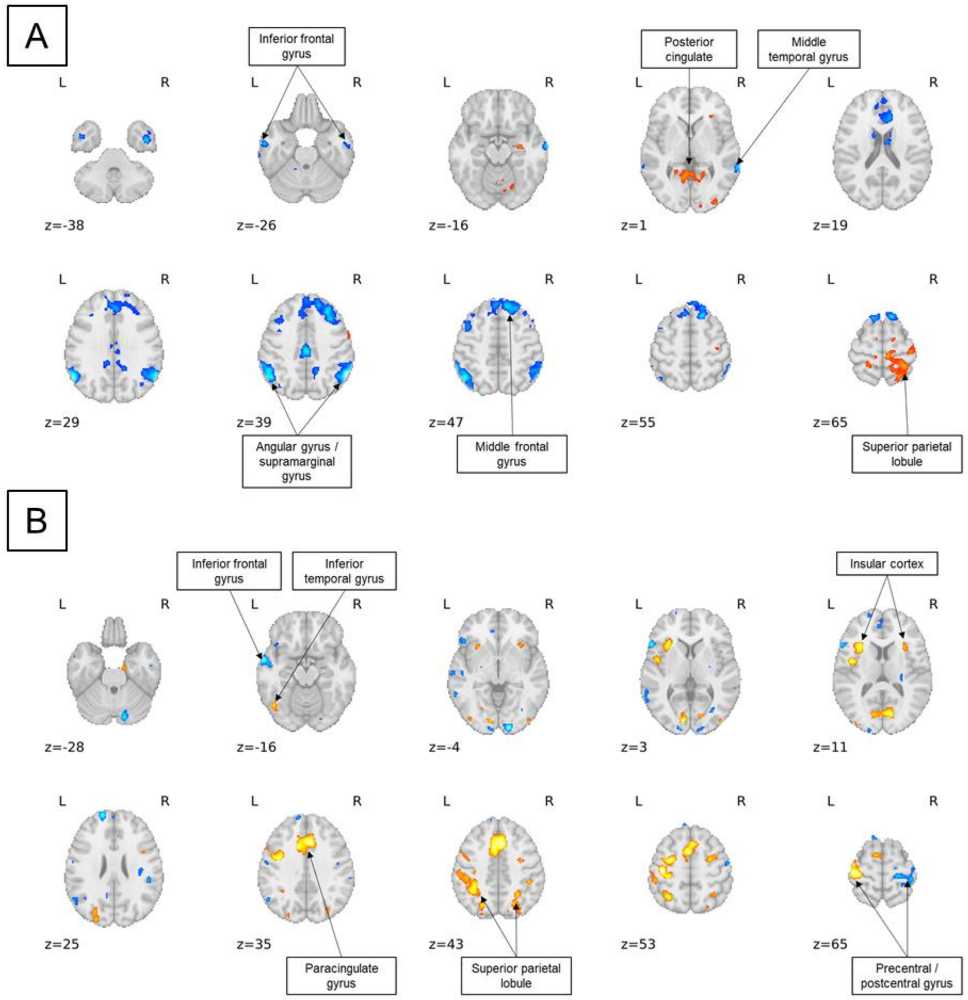


*Figure S10. Axial slices of predictive maps of the summary measure Verbal for the contrasts (A) pseudoword–pseudohomophone (from PhonLex), and (B) correct (from Learn). The threshold of the displayed consensus maps was set at z = 2.*

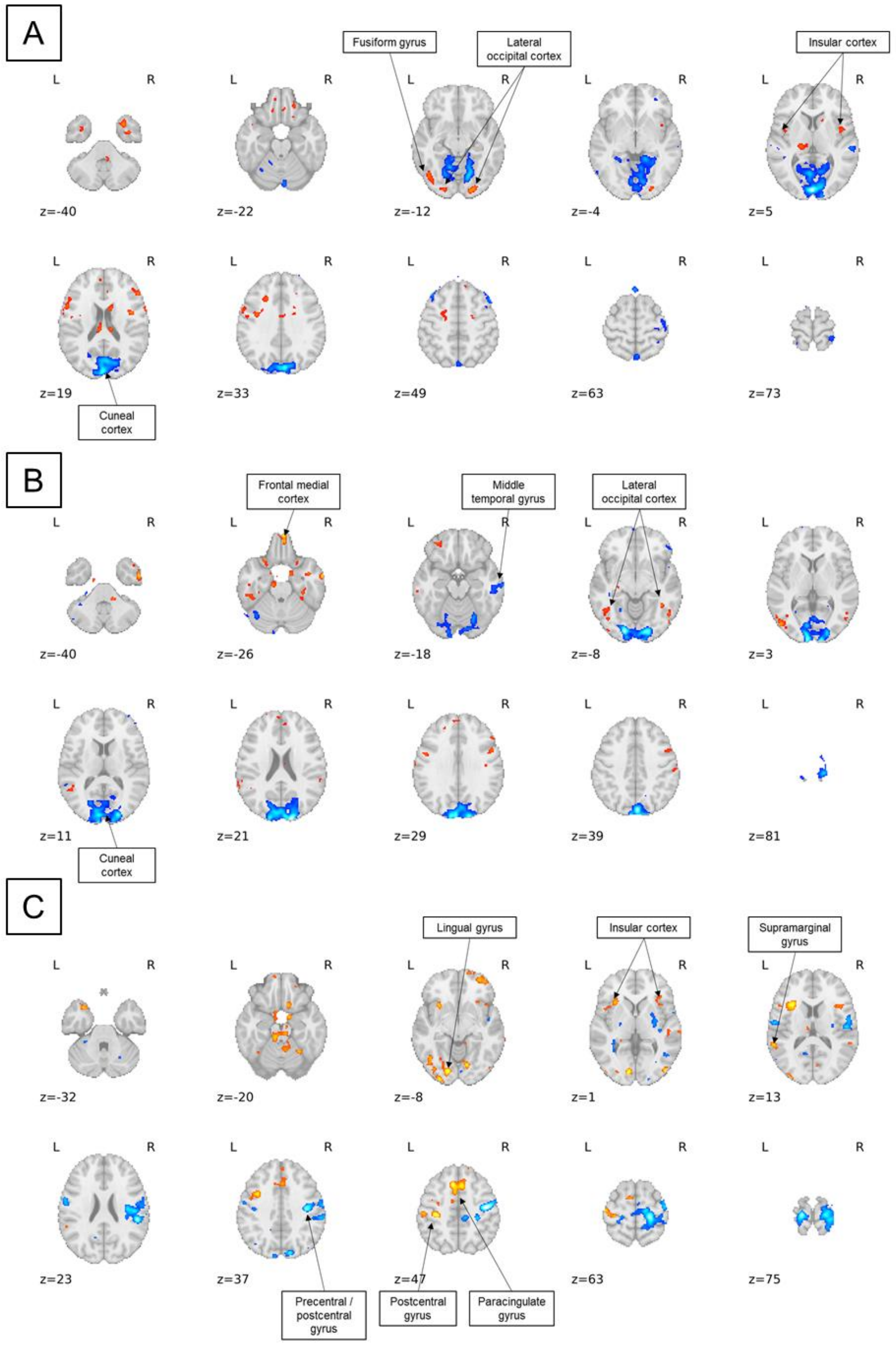


*Figure S11. Axial slices of predictive maps of the summary measure Naming for the contrasts (A) pseudohomophone (from PhonLex), (B) FFnew (from CharProc), and (C) incorrect (from Learn). The threshold of the displayed consensus maps was set at z = 2.*

*Table S1. Clusters obtained at a threshold > 2 for the pseudohomophone contrast in the PhonLex task and the prediction of the Reading summary score.*

| Sign | MNI coordinates (mm) X | Y | Z | Peak Stat | Cluster Size ($mm^3$) | Label |
|---|---|---|---|---|---|---|
| **Positive** | 11.5 | 1.5 | 25.5 | 3.608 | 1248 | Unknown |
| | 1.5 | -4.5 | 25.5 | 2.449 | | Unknown |
| | -8.5 | -12.5 | 25.5 | 2.138 | | Unknown |
| | 29.5 | 9.5 | -40.5 | 3.567 | 2464 | Temporal Pole |
| | 31.5 | -2.5 | -42.5 | 2.992 | | Temporal Fusiform Cortex, anterior division |
| | 39.5 | -2.5 | -40.5 | 2.878 | | Inferior Temporal Gyrus, anterior division |
| | 47.5 | -2.5 | -36.5 | 2.529 | | Inferior Temporal Gyrus, anterior division |
| | 23.5 | -92.5 | -8.5 | 3.449 | 1392 | Occipital Pole |
| | -54.5 | -2.5 | 27.5 | 3.35 | 6544 | Precentral Gyrus |
| | -54.5 | -12.5 | 25.5 | 2.964 | | Postcentral Gyrus |
| | -58.5 | -2.5 | 35.5 | 2.84 | | Precentral Gyrus |
| | -60.5 | -2.5 | 21.5 | 2.579 | | Precentral Gyrus |
| | -44.5 | -68.5 | -12.5 | 3.01 | 864 | Lateral Occipital Cortex, inferior division |
| | 31.5 | -22.5 | 27.5 | 2.984 | 928 | Unknown |
| | 49.5 | -0.5 | 9.5 | 2.902 | 2288 | Central Opercular Cortex |
| | 41.5 | -0.5 | 19.5 | 2.834 | | Unknown |
| | 45.5 | 5.5 | 1.5 | 2.569 | | Central Opercular Cortex |
| | 39.5 | 9.5 | 11.5 | 2.119 | | Frontal Opercular Cortex |
| | -20.5 | -90.5 | -12.5 | 2.885 | 888 | Occipital Fusiform Gyrus |
| | -16.5 | -98.5 | -8.5 | 2.497 | | Occipital Pole |
| | -32.5 | 3.5 | -34.5 | 2.798 | 1072 | Unknown |
| | -40.5 | 9.5 | -40.5 | 2.494 | | Temporal Pole |
| | 21.5 | -26.5 | 43.5 | 2.757 | 352 | Unknown |
| | -46.5 | 11.5 | -26.5 | 2.756 | 624 | Temporal Pole |
| | 65.5 | 3.5 | 19.5 | 2.747 | 240 | Precentral Gyrus |
| | 9.5 | -42.5 | -36.5 | 2.745 | 296 | Unknown |
| | 11.5 | 9.5 | 39.5 | 2.744 | 208 | Cingulate Gyrus, anterior division |
| | -18.5 | -24.5 | 7.5 | 2.724 | 896 | Unknown |
| | -8.5 | -22.5 | 7.5 | 2.355 | | Unknown |
| | -58.5 | -42.5 | 31.5 | 2.686 | 240 | Supramarginal Gyrus, posterior division |
| | -6.5 | -16.5 | -46.5 | 2.667 | 488 | Unknown |
| | 1.5 | -16.5 | -36.5 | 2.326 | | Unknown |
| | -18.5 | -12.5 | -32.5 | 2.656 | 360 | Parahippocampal Gyrus, anterior division |
| | 15.5 | -26.5 | 23.5 | 2.558 | 520 | Unknown |
| | -58.5 | 3.5 | -28.5 | 2.548 | 352 | Temporal Pole |
| | -62.5 | -2.5 | -24.5 | 2.202 | | Middle Temporal Gyrus, anterior division |
| | 5.5 | 31.5 | 17.5 | 2.539 | 288 | Cingulate Gyrus, anterior division |
| | -0.5 | -54.5 | -34.5 | 2.523 | 160 | Unknown |
| | -2.5 | -8.5 | 3.5 | 2.516 | 192 | Unknown |
| | -42.5 | -0.5 | 5.5 | 2.495 | 560 | Central Opercular Cortex |
| | -50.5 | -2.5 | 3.5 | 2.379 | | Central Opercular Cortex |
| | -10.5 | -30.5 | 21.5 | 2.478 | 184 | Unknown |
| | -12.5 | 5.5 | 43.5 | 2.455 | 248 | Unknown |
| | -2.5 | 1.5 | 57.5 | 2.452 | 304 | Juxtapositional Lobule Cortex (formerly Supplementary Motor Cortex) |
| | -46.5 | -0.5 | 39.5 | 2.445 | 456 | Precentral Gyrus |
| | -38.5 | -2.5 | 35.5 | 2.428 | | Precentral Gyrus |
| | -60.5 | -28.5 | 27.5 | 2.405 | 160 | Supramarginal Gyrus, anterior division |
| | -8.5 | -8.5 | 51.5 | 2.403 | 264 | Juxtapositional Lobule Cortex (formerly Supplementary Motor Cortex) |
| | 49.5 | -24.5 | 29.5 | 2.402 | 216 | Unknown |
| | -24.5 | 19.5 | 33.5 | 2.391 | 312 | Unknown |
| | -0.5 | 51.5 | 21.5 | 2.31 | 288 | Paracingulate Gyrus |
| **Negative** | -2.5 | -92.5 | 5.5 | 4.5 | 47168 | Occipital Pole |
| | 5.5 | -84.5 | 33.5 | 3.901 | | Cuneal Cortex |
| | -2.5 | -88.5 | 21.5 | 3.851 | | Cuneal Cortex |
| | 15.5 | -78.5 | 21.5 | 3.757 | | Cuneal Cortex |
| | 31.5 | 47.5 | -0.5 | 3.738 | 1832 | Unknown |
| | 33.5 | 35.5 | 1.5 | 2.925 | | Unknown |
| | 45.5 | 31.5 | 43.5 | 3.326 | 2480 | Unknown |
| | 45.5 | 21.5 | 53.5 | 3.259 | | Unknown |
| | 39.5 | 13.5 | 45.5 | 2.875 | | Middle Frontal Gyrus |
| | 37.5 | 9.5 | 57.5 | 2.592 | | Middle Frontal Gyrus |
| | -34.5 | 59.5 | -0.5 | 3.19 | 896 | Frontal Pole |
| | -42.5 | 51.5 | 7.5 | 2.404 | | Frontal Pole |
| | -40.5 | 29.5 | 47.5 | 3.135 | 520 | Unknown |
| | -40.5 | 23.5 | 53.5 | 3.013 | | Unknown |
| | -0.5 | 35.5 | 65.5 | 3.012 | 192 | Unknown |
| | -20.5 | -38.5 | -50.5 | 2.837 | 256 | Unknown |
| | -2.5 | 57.5 | 45.5 | 2.8 | 224 | Unknown |
| | -32.5 | -72.5 | -48.5 | 2.632 | 544 | Unknown |
| | -8.5 | -44.5 | -14.5 | 2.624 | 304 | Unknown |
| | 59.5 | -26.5 | 5.5 | 2.621 | 536 | Unknown |
| | 65.5 | -30.5 | 9.5 | 2.507 | | Superior Temporal Gyrus, posterior division |
| | 41.5 | -16.5 | 61.5 | 2.618 | 616 | Precentral Gyrus |
| | 45.5 | -28.5 | 65.5 | 2.382 | | Postcentral Gyrus |
| | 33.5 | -46.5 | 69.5 | 2.61 | 584 | Superior Parietal Lobule |
| | 27.5 | -40.5 | 73.5 | 2.397 | | Unknown |
| | 3.5 | -68.5 | 63.5 | 2.55 | 304 | Unknown |
| | -34.5 | 11.5 | 57.5 | 2.471 | 192 | Middle Frontal Gyrus |
| | 35.5 | -26.5 | 65.5 | 2.446 | 456 | Precentral Gyrus |
| | 57.5 | -28.5 | -10.5 | 2.391 | 176 | Middle Temporal Gyrus, posterior division |
| | 61.5 | -32.5 | -4.5 | 2.127 | | Middle Temporal Gyrus, posterior division |

*Table S2. Clusters obtained at a threshold > 2 for the "incorrect" contrast in the Learn task and the prediction of the Reading summary score.*

| Sign | MNI coordinates (mm) | | | Peak Stat | Cluster Size (mm³) | Label |
|---|---|---|---|---|---|---|
| | X | Y | Z | | | |
| Positive | -32.5 | -28.5 | 51.5 | 4.293 | 8368 | Postcentral Gyrus |
| | -46.5 | -18.5 | 63.5 | 3.557 | | Unknown |
| | -52.5 | -22.5 | 49.5 | 3.444 | | Postcentral Gyrus |
| | -38.5 | -2.5 | 55.5 | 3.36 | | Middle Frontal Gyrus |
| | -16.5 | -84.5 | -8.5 | 4.188 | 2464 | Unknown |
| | -12.5 | -86.5 | -0.5 | 3.64 | | Unknown |
| | -32.5 | 17.5 | 13.5 | 4.027 | 5696 | Unknown |
| | -32.5 | 21.5 | -0.5 | 3 | | Insular Cortex |
| | -46.5 | 11.5 | 3.5 | 2.583 | | Inferior Frontal Gyrus, pars opercularis |
| | -4.5 | 11.5 | 51.5 | 3.988 | 12640 | Paracingulate Gyrus |
| | -8.5 | 21.5 | 47.5 | 3.626 | | Unknown |
| | 5.5 | 25.5 | 43.5 | 3.57 | | Paracingulate Gyrus |
| | -8.5 | -6.5 | 53.5 | 3.49 | | Juxtapositional Lobule Cortex (formerly Supplementary Motor Cortex) |
| | 13.5 | -74.5 | -10.5 | 3.646 | 1528 | Lingual Gyrus |
| | -58.5 | -44.5 | 15.5 | 3.462 | 1032 | Supramarginal Gyrus, posterior division |
| | -64.5 | -38.5 | 9.5 | 2.332 | | Superior Temporal Gyrus, posterior division |
| | 45.5 | -84.5 | 3.5 | 3.349 | 768 | Lateral Occipital Cortex, inferior division |
| | 51.5 | -78.5 | 5.5 | 2.927 | | Lateral Occipital Cortex, inferior division |
| | -8.5 | -30.5 | -20.5 | 3.21 | 6120 | Unknown |
| | 27.5 | -54.5 | -20.5 | 3.057 | | Unknown |
| | 7.5 | -38.5 | -16.5 | 2.872 | | Unknown |
| | -14.5 | -24.5 | -18.5 | 2.826 | | Unknown |
| | -28.5 | -96.5 | -10.5 | 3.168 | 528 | Occipital Pole |
| | 45.5 | 13.5 | 9.5 | 3.127 | 3592 | Unknown |
| | 33.5 | 27.5 | 3.5 | 2.661 | | Unknown |
| | 29.5 | 23.5 | -6.5 | 2.331 | | Frontal Orbital Cortex |
| | 41.5 | 19.5 | -6.5 | 2.283 | | Insular Cortex |
| | 13.5 | -0.5 | -20.5 | 3.065 | 488 | Unknown |
| | -10.5 | -2.5 | -24.5 | 3.04 | 1648 | Unknown |
| | 5.5 | -10.5 | -14.5 | 2.423 | | Unknown |
| | -6.5 | -12.5 | -14.5 | 2.396 | | Unknown |
| | -32.5 | 13.5 | -32.5 | 2.969 | 624 | Temporal Pole |
| | -38.5 | 25.5 | -32.5 | 2.125 | | Temporal Pole |
| | -32.5 | -52.5 | -16.5 | 2.967 | 1080 | Temporal Occipital Fusiform Cortex |
| | -26.5 | -62.5 | -12.5 | 2.771 | | Temporal Occipital Fusiform Cortex |
| | -44.5 | -66.5 | -10.5 | 2.842 | 1872 | Lateral Occipital Cortex, inferior division |
| | -40.5 | -82.5 | -14.5 | 2.704 | | Lateral Occipital Cortex, inferior division |
| | -38.5 | -90.5 | 11.5 | 2.784 | 336 | Occipital Pole |
| | -20.5 | -96.5 | 19.5 | 2.759 | 208 | Occipital Pole |
| | -36.5 | 5.5 | 35.5 | 2.747 | 1056 | Middle Frontal Gyrus |
| | -46.5 | 1.5 | 33.5 | 2.526 | | Precentral Gyrus |
| | -6.5 | -18.5 | 11.5 | 2.598 | 472 | Unknown |
| | 7.5 | -4.5 | 27.5 | 2.575 | 232 | Unknown |
| | 11.5 | 17.5 | -20.5 | 2.568 | 368 | Subcallosal Cortex |
| | 41.5 | 53.5 | -10.5 | 2.563 | 664 | Frontal Pole |
| | 33.5 | 57.5 | -16.5 | 2.464 | | Frontal Pole |
| | 31.5 | 55.5 | -6.5 | 2.383 | | Frontal Pole |
| | -40.5 | -4.5 | 15.5 | 2.526 | 264 | Central Opercular Cortex |
| | -32.5 | -26.5 | -24.5 | 2.409 | 208 | Temporal Fusiform Cortex, posterior division |
| | -28.5 | -62.5 | 57.5 | 2.379 | 264 | Lateral Occipital Cortex, superior division |
| Negative | 41.5 | -14.5 | 47.5 | 3.959 | 13784 | Precentral Gyrus |
| | 37.5 | -16.5 | 23.5 | 3.16 | | Unknown |
| | 49.5 | -30.5 | 25.5 | 3.151 | | Parietal Opercular Cortex |
| | 45.5 | -20.5 | 21.5 | 2.961 | | Parietal Opercular Cortex |
| | 21.5 | -28.5 | 63.5 | 3.814 | 11504 | Precentral Gyrus |
| | 15.5 | -18.5 | 65.5 | 3.497 | | Precentral Gyrus |
| | 19.5 | -24.5 | 77.5 | 3.377 | | Precentral Gyrus |
| | 25.5 | -32.5 | 75.5 | 2.847 | | Postcentral Gyrus |
| | -16.5 | -26.5 | 75.5 | 3.642 | 2120 | Precentral Gyrus |
| | 13.5 | -84.5 | 39.5 | 3.462 | 1144 | Lateral Occipital Cortex, superior division |
| | -26.5 | -36.5 | 59.5 | 2.842 | 592 | Postcentral Gyrus |
| | -22.5 | -10.5 | -0.5 | 2.806 | 208 | Unknown |
| | 29.5 | -30.5 | 45.5 | 2.792 | 264 | Unknown |
| | -8.5 | -56.5 | 9.5 | 2.789 | 512 | Precuneous Cortex |
| | 41.5 | -78.5 | 33.5 | 2.714 | 1048 | Lateral Occipital Cortex, superior division |
| | 41.5 | -56.5 | 33.5 | 2.47 | | Unknown |
| | 39.5 | -70.5 | 35.5 | 2.3 | | Lateral Occipital Cortex, superior division |
| | -38.5 | -70.5 | 37.5 | 2.7 | 272 | Lateral Occipital Cortex, superior division |
| | 47.5 | -58.5 | -2.5 | 2.664 | 608 | Unknown |
| | 59.5 | -56.5 | -4.5 | 2.47 | | Middle Temporal Gyrus, temporooccipital part |
| | 31.5 | -72.5 | 5.5 | 2.648 | 224 | Unknown |
| | -8.5 | -88.5 | 37.5 | 2.606 | 880 | Occipital Pole |
| | -12.5 | -84.5 | 29.5 | 2.576 | | Unknown |
| | -24.5 | 41.5 | -2.5 | 2.583 | 160 | Unknown |
| | 27.5 | -14.5 | 3.5 | 2.576 | 496 | Unknown |
| | 37.5 | -40.5 | -0.5 | 2.523 | 232 | Unknown |
| | -34.5 | -30.5 | 31.5 | 2.481 | 264 | Unknown |
| | -58.5 | -10.5 | 11.5 | 2.481 | 312 | Central Opercular Cortex |
| | -18.5 | -46.5 | 35.5 | 2.474 | 232 | Unknown |
| | -56.5 | -6.5 | 35.5 | 2.473 | 544 | Precentral Gyrus |
| | -48.5 | -10.5 | 33.5 | 2.274 | | Precentral Gyrus |
| | -58.5 | -8.5 | 25.5 | 2.214 | | Postcentral Gyrus |
| | -34.5 | -50.5 | 1.5 | 2.458 | 296 | Unknown |
| | -32.5 | -38.5 | 1.5 | 2.113 | | Unknown |
| | 23.5 | -68.5 | -40.5 | 2.45 | 376 | Unknown |

| | | | | | | |
|---|---|---|---|---|---|---|
| | 23.5 | -64.5 | -32.5 | 2.359 | | Unknown |
| | 9.5 | -30.5 | 47.5 | 2.436 | 416 | Precentral Gyrus |
| | -40.5 | -16.5 | 39.5 | 2.411 | 304 | Precentral Gyrus |
| | -24.5 | -66.5 | -42.5 | 2.342 | 232 | Unknown |

*Table S3. Clusters obtained at a threshold > 2 for the "letter" contrast in the CharProc task and the prediction of the Reading summary score.*

| Sign | MNI coordinates (mm) | | | Peak Stat | Cluster Size ($mm^3$) | Label |
|---|---|---|---|---|---|---|
| | X | Y | Z | | | |
| Positive | 5.5 | 49.5 | -26.5 | 4.481 | 1480 | Frontal Medial Cortex |
| | -28.5 | 25.5 | -22.5 | 3.647 | 3000 | Frontal Orbital Cortex |
| | -24.5 | 49.5 | -18.5 | 3.161 | | Frontal Pole |
| | -32.5 | 47.5 | -18.5 | 3.125 | | Frontal Pole |
| | -26.5 | 39.5 | -18.5 | 2.902 | | Frontal Pole |
| | -34.5 | -8.5 | 51.5 | 3.351 | 1736 | Precentral Gyrus |
| | -38.5 | 1.5 | 53.5 | 2.803 | | Middle Frontal Gyrus |
| | -0.5 | 27.5 | -22.5 | 3.342 | 1392 | Subcallosal Cortex |
| | 9.5 | 33.5 | -22.5 | 3.034 | | Frontal Medial Cortex |
| | 23.5 | 45.5 | -20.5 | 2.862 | | Frontal Pole |
| | -50.5 | 1.5 | 27.5 | 3.338 | 1144 | Precentral Gyrus |
| | -60.5 | 9.5 | 25.5 | 3.298 | | Unknown |
| | -14.5 | 29.5 | 53.5 | 3.23 | 936 | Superior Frontal Gyrus |
| | -66.5 | -8.5 | -14.5 | 3.196 | 472 | Unknown |
| | -54.5 | -18.5 | -20.5 | 2.407 | | Unknown |
| | -66.5 | -14.5 | -20.5 | 2.132 | | Middle Temporal Gyrus, posterior division |
| | -38.5 | -70.5 | -4.5 | 3.121 | 2048 | Lateral Occipital Cortex, inferior division |
| | -44.5 | -56.5 | -6.5 | 2.72 | | Inferior Temporal Gyrus, temporooccipital part |
| | 63.5 | -2.5 | -26.5 | 3.109 | 864 | Middle Temporal Gyrus, anterior division |
| | 57.5 | -4.5 | -32.5 | 2.725 | | Middle Temporal Gyrus, anterior division |
| | 59.5 | -14.5 | -34.5 | 2.356 | | Inferior Temporal Gyrus, posterior division |
| | 23.5 | 15.5 | -28.5 | 3.083 | 296 | Unknown |
| | -50.5 | -80.5 | -0.5 | 3.061 | 496 | Lateral Occipital Cortex, inferior division |
| | 7.5 | 19.5 | 47.5 | 3.007 | 2240 | Paracingulate Gyrus |
| | 19.5 | 39.5 | 51.5 | 2.825 | | Frontal Pole |
| | 1.5 | 31.5 | 49.5 | 2.664 | | Superior Frontal Gyrus |
| | 5.5 | 27.5 | 41.5 | 2.647 | | Paracingulate Gyrus |
| | -10.5 | -76.5 | -48.5 | 2.994 | 328 | Unknown |
| | -34.5 | 11.5 | -36.5 | 2.987 | 712 | Temporal Pole |
| | -46.5 | 13.5 | -32.5 | 2.581 | | Temporal Pole |
| | -42.5 | 7.5 | -38.5 | 2.134 | | Temporal Pole |
| | -54.5 | -32.5 | 51.5 | 2.979 | 1808 | Supramarginal Gyrus, anterior division |
| | -48.5 | -40.5 | 51.5 | 2.822 | | Supramarginal Gyrus, posterior division |
| | -50.5 | -32.5 | -26.5 | 2.969 | 328 | Inferior Temporal Gyrus, posterior division |
| | 31.5 | 19.5 | 17.5 | 2.952 | 208 | Unknown |
| | 21.5 | 25.5 | -24.5 | 2.88 | 280 | Frontal Orbital Cortex |
| | -62.5 | -28.5 | -20.5 | 2.837 | 480 | Middle Temporal Gyrus, posterior division |
| | 51.5 | 29.5 | 31.5 | 2.824 | 800 | Middle Frontal Gyrus |
| | 49.5 | 21.5 | 35.5 | 2.437 | | Middle Frontal Gyrus |
| | -28.5 | -96.5 | -4.5 | 2.82 | 232 | Occipital Pole |
| | -2.5 | 39.5 | -32.5 | 2.81 | 168 | Unknown |
| | 13.5 | 47.5 | 43.5 | 2.781 | 304 | Frontal Pole |
| | -12.5 | -12.5 | -30.5 | 2.771 | 824 | Unknown |
| | -6.5 | -4.5 | -30.5 | 2.477 | | Unknown |
| | -16.5 | -10.5 | -38.5 | 2.34 | | Unknown |
| | -10.5 | 7.5 | -22.5 | 2.741 | 288 | Frontal Orbital Cortex |
| | -16.5 | -0.5 | -26.5 | 2.369 | | Parahippocampal Gyrus, anterior division |
| | 37.5 | -26.5 | -22.5 | 2.678 | 240 | Temporal Fusiform Cortex, posterior division |
| | -20.5 | -16.5 | -6.5 | 2.667 | 272 | Unknown |
| | 47.5 | 5.5 | 37.5 | 2.615 | 1456 | Precentral Gyrus |
| | 55.5 | 7.5 | 41.5 | 2.585 | | Precentral Gyrus |
| | 51.5 | 7.5 | 29.5 | 2.457 | | Precentral Gyrus |
| | 33.5 | 15.5 | 35.5 | 2.602 | 464 | Unknown |
| | 33.5 | 23.5 | 31.5 | 2.596 | | Middle Frontal Gyrus |
| | -2.5 | 3.5 | 53.5 | 2.583 | 448 | Juxtapositional Lobule Cortex (formerly Supplementary Motor Cortex) |
| | -4.5 | 11.5 | 51.5 | 2.306 | | Paracingulate Gyrus |
| | -42.5 | -88.5 | -10.5 | 2.552 | 208 | Lateral Occipital Cortex, inferior division |
| | 15.5 | 1.5 | -26.5 | 2.513 | 216 | Parahippocampal Gyrus, anterior division |
| | 15.5 | 9.5 | -24.5 | 2.12 | | Frontal Orbital Cortex |
| | 43.5 | 1.5 | -28.5 | 2.469 | 200 | Unknown |
| | -40.5 | 25.5 | 21.5 | 2.449 | 176 | Middle Frontal Gyrus |
| | -56.5 | 7.5 | 11.5 | 2.448 | 200 | Precentral Gyrus |
| | -30.5 | -14.5 | -24.5 | 2.367 | 256 | Unknown |
| | -28.5 | -4.5 | -24.5 | 2.3 | | Unknown |
| | 51.5 | -30.5 | 51.5 | 2.335 | 232 | Postcentral Gyrus |
| | -26.5 | 11.5 | 45.5 | 2.334 | 160 | Middle Frontal Gyrus |
| | 53.5 | -72.5 | -4.5 | 2.295 | 184 | Lateral Occipital Cortex, inferior division |
| Negative | 13.5 | -98.5 | 23.5 | 5.229 | 49568 | Occipital Pole |
| | -10.5 | -94.5 | 15.5 | 4.547 | | Occipital Pole |
| | -22.5 | -70.5 | 15.5 | 4.488 | | Unknown |
| | 3.5 | -84.5 | 39.5 | 4.256 | | Cuneal Cortex |
| | 63.5 | -10.5 | -4.5 | 3.781 | 672 | Unknown |
| | -24.5 | -24.5 | 79.5 | 3.479 | 408 | Unknown |
| | -28.5 | -34.5 | 75.5 | 2.524 | | Unknown |
| | 17.5 | -26.5 | 81.5 | 3.288 | 1264 | Unknown |
| | 17.5 | -6.5 | 79.5 | 2.642 | | Unknown |
| | 17.5 | 1.5 | 77.5 | 2.578 | | Unknown |
| | 25.5 | 19.5 | 67.5 | 3.06 | 496 | Unknown |
| | 21.5 | 29.5 | 63.5 | 2.695 | | Unknown |
| | 5.5 | 41.5 | 61.5 | 3.012 | 408 | Unknown |
| | -10.5 | 35.5 | 63.5 | 2.575 | | Unknown |
| | 15.5 | 37.5 | 61.5 | 2.518 | | Unknown |
| | -2.5 | 39.5 | 63.5 | 2.357 | | Unknown |
| | -34.5 | 9.5 | 65.5 | 2.94 | 208 | Unknown |
| | -8.5 | -44.5 | 53.5 | 2.93 | 480 | Precuneous Cortex |

| | | | | | | |
|---|---|---|---|---|---|---|
| | -14.5 | -2.5 | 81.5 | 2.912 | 184 | Unknown |
| | -8.5 | -50.5 | 79.5 | 2.851 | 176 | Unknown |
| | -52.5 | -50.5 | -30.5 | 2.82 | 384 | Unknown |
| | -56.5 | -62.5 | -28.5 | 2.506 | | Unknown |
| | -44.5 | -82.5 | 19.5 | 2.801 | 232 | Lateral Occipital Cortex, superior division |
| | 23.5 | -48.5 | 3.5 | 2.65 | 496 | Unknown |
| | 23.5 | -54.5 | -2.5 | 2.606 | | Lingual Gyrus |
| | -16.5 | 11.5 | 75.5 | 2.615 | 272 | Unknown |
| | -24.5 | 9.5 | 71.5 | 2.603 | | Unknown |
| | 15.5 | -50.5 | 45.5 | 2.556 | 176 | Unknown |
| | -18.5 | -62.5 | -14.5 | 2.53 | 296 | Lingual Gyrus |
| | -18.5 | -46.5 | 65.5 | 2.482 | 528 | Postcentral Gyrus |
| | -40.5 | -34.5 | -12.5 | 2.46 | 200 | Unknown |
| | -20.5 | -30.5 | 63.5 | 2.425 | 232 | Postcentral Gyrus |

*Table S4. Clusters obtained at a threshold > 2 for the word–face contrast in the Localizer task and the prediction of the Reading summary score.*

| Sign | MNI coordinates (mm) X | Y | Z | Peak Stat | Cluster Size ($mm^3$) | Label |
|---|---|---|---|---|---|---|
| Negative | -12.5 | -102.5 | 1.5 | 6.786 | 2304 | Occipital Pole |
| | 13.5 | -98.5 | 3.5 | 4.799 | 4632 | Occipital Pole |
| | 9.5 | -88.5 | -4.5 | 4.554 | | Lingual Gyrus |
| | 13.5 | -100.5 | 13.5 | 3.717 | | Occipital Pole |
| | 25.5 | -100.5 | 3.5 | 3.415 | | Occipital Pole |
| | -8.5 | 53.5 | 17.5 | 3.809 | 10504 | Unknown |
| | -4.5 | 45.5 | 43.5 | 3.317 | | Superior Frontal Gyrus |
| | 3.5 | 49.5 | 17.5 | 3.305 | | Paracingulate Gyrus |
| | 9.5 | 49.5 | 23.5 | 3.209 | | Unknown |
| | 47.5 | -10.5 | 45.5 | 3.333 | 6864 | Precentral Gyrus |
| | 37.5 | -12.5 | 47.5 | 2.809 | | Precentral Gyrus |
| | 53.5 | -8.5 | 11.5 | 2.796 | | Central Opercular Cortex |
| | 59.5 | -0.5 | 31.5 | 2.649 | | Precentral Gyrus |
| | -8.5 | -48.5 | 33.5 | 2.889 | 1776 | Cingulate Gyrus, posterior division |
| | 9.5 | -52.5 | 37.5 | 2.339 | | Precuneous Cortex |
| | 13.5 | 31.5 | 65.5 | 2.735 | 344 | Unknown |
| | 39.5 | 23.5 | -18.5 | 2.688 | 160 | Frontal Orbital Cortex |
| | -56.5 | -6.5 | 39.5 | 2.618 | 608 | Precentral Gyrus |
| | -54.5 | 3.5 | -26.5 | 2.61 | 512 | Temporal Pole |
| | -44.5 | -72.5 | 45.5 | 2.598 | 264 | Lateral Occipital Cortex, superior division |
| | -40.5 | -16.5 | 43.5 | 2.586 | 432 | Precentral Gyrus |
| | -62.5 | -2.5 | 9.5 | 2.475 | 304 | Precentral Gyrus |
| | -60.5 | -12.5 | 9.5 | 2.084 | | Central Opercular Cortex |
| | 17.5 | 57.5 | 17.5 | 2.461 | 176 | Frontal Pole |
| | -12.5 | -64.5 | -16.5 | 2.456 | 568 | Unknown |
| | -16.5 | -54.5 | -18.5 | 2.413 | | Unknown |
| | -10.5 | -62.5 | -20.5 | 2.39 | | Unknown |
| | -12.5 | -48.5 | -16.5 | 2.084 | | Unknown |
| | -8.5 | 29.5 | 61.5 | 2.428 | 440 | Superior Frontal Gyrus |
| | -8.5 | 27.5 | 69.5 | 2.4 | | Unknown |
| | -16.5 | 33.5 | 61.5 | 2.227 | | Unknown |
| | 27.5 | -76.5 | -34.5 | 2.359 | 216 | Unknown |
| | -2.5 | 37.5 | -6.5 | 2.31 | 192 | Cingulate Gyrus, anterior division |
| | 5.5 | 39.5 | -6.5 | 2.144 | | Paracingulate Gyrus |
| | -24.5 | -32.5 | 65.5 | 2.266 | 168 | Postcentral Gyrus |
| Positive | -50.5 | -60.5 | -10.5 | 5.885 | 26904 | Inferior Temporal Gyrus, temporooccipital part |
| | -42.5 | -66.5 | -2.5 | 5.656 | | Lateral Occipital Cortex, inferior division |
| | -56.5 | -58.5 | 7.5 | 4.967 | | Middle Temporal Gyrus, temporooccipital part |
| | -46.5 | -42.5 | -20.5 | 4.358 | | Inferior Temporal Gyrus, posterior division |
| | -22.5 | -94.5 | 1.5 | 4.047 | 1192 | Occipital Pole |
| | -20.5 | -98.5 | 9.5 | 2.347 | | Occipital Pole |
| | -14.5 | -84.5 | -12.5 | 3.683 | 640 | Occipital Fusiform Gyrus |
| | 49.5 | 11.5 | 21.5 | 3.654 | 2328 | Inferior Frontal Gyrus, pars opercularis |
| | -40.5 | -44.5 | 51.5 | 3.472 | 8512 | Superior Parietal Lobule |
| | -50.5 | -36.5 | 59.5 | 3.185 | | Unknown |
| | -30.5 | -64.5 | 47.5 | 2.831 | | Lateral Occipital Cortex, superior division |
| | -58.5 | -22.5 | 35.5 | 2.709 | | Postcentral Gyrus |
| | 57.5 | -42.5 | -16.5 | 3.274 | 2800 | Inferior Temporal Gyrus, temporooccipital part |
| | 63.5 | -54.5 | -12.5 | 3.102 | | Inferior Temporal Gyrus, temporooccipital part |
| | 67.5 | -28.5 | -20.5 | 2.669 | | Middle Temporal Gyrus, posterior division |
| | 67.5 | -38.5 | -18.5 | 2.629 | | Unknown |
| | -42.5 | 9.5 | 29.5 | 3.208 | 7696 | Middle Frontal Gyrus |
| | -52.5 | 9.5 | 25.5 | 3.149 | | Precentral Gyrus |
| | -40.5 | 27.5 | 19.5 | 3.143 | | Inferior Frontal Gyrus, pars triangularis |
| | -58.5 | 13.5 | 33.5 | 2.921 | | Unknown |
| | -34.5 | -12.5 | -46.5 | 3.183 | 472 | Unknown |
| | -42.5 | -4.5 | -48.5 | 2.204 | | Inferior Temporal Gyrus, anterior division |
| | -34.5 | -90.5 | 9.5 | 3.05 | 408 | Occipital Pole |
| | -26.5 | -88.5 | 7.5 | 2.38 | | Unknown |
| | 19.5 | -100.5 | -12.5 | 3.036 | 264 | Occipital Pole |
| | 51.5 | -20.5 | -2.5 | 3.027 | 1400 | Superior Temporal Gyrus, posterior division |
| | 49.5 | -28.5 | 5.5 | 2.203 | | Unknown |
| | 35.5 | -12.5 | -30.5 | 2.786 | 408 | Temporal Fusiform Cortex, posterior division |
| | 31.5 | 51.5 | -2.5 | 2.566 | 224 | Frontal Pole |
| | -56.5 | -44.5 | 29.5 | 2.525 | 224 | Supramarginal Gyrus, posterior division |
| | 33.5 | -84.5 | -6.5 | 2.448 | 200 | Lateral Occipital Cortex, inferior division |
| | 35.5 | -92.5 | -8.5 | 2.326 | | Occipital Pole |
| | -40.5 | -12.5 | -30.5 | 2.433 | 216 | Temporal Fusiform Cortex, posterior division |
| | -44.5 | -16.5 | -36.5 | 2.411 | | Inferior Temporal Gyrus, posterior division |
| | 41.5 | 17.5 | -2.5 | 2.281 | 208 | Insular Cortex |

*Table S5. Clusters obtained at a threshold > 2 for the pseudoword–pseudohomophone contrast in the PhonLex task and the prediction of the Verbal summary score.*

| Sign | MNI coordinates (mm) X | Y | Z | Peak Stat | Cluster Size (mm³) | Label |
|---|---|---|---|---|---|---|
| Positive | 7.5 | -30.5 | 67.5 | 3.228 | 7256 | Unknown |
| | 15.5 | -48.5 | 71.5 | 3.062 | | Superior Parietal Lobule |
| | 31.5 | -44.5 | 65.5 | 3.01 | | Superior Parietal Lobule |
| | 15.5 | -54.5 | 71.5 | 2.943 | | Superior Parietal Lobule |
| | 15.5 | -50.5 | 5.5 | 3.02 | 6080 | Cingulate Gyrus, posterior division |
| | -8.5 | -56.5 | 5.5 | 2.976 | | Precuneous Cortex |
| | -2.5 | -56.5 | -10.5 | 2.606 | | Unknown |
| | 1.5 | -50.5 | 1.5 | 2.555 | | Unknown |
| | 27.5 | -12.5 | 75.5 | 2.801 | 384 | Unknown |
| | 29.5 | -20.5 | 75.5 | 2.418 | | Unknown |
| | -0.5 | 3.5 | -10.5 | 2.702 | 296 | Unknown |
| | 33.5 | -92.5 | -0.5 | 2.693 | 536 | Occipital Pole |
| | 31.5 | -12.5 | -16.5 | 2.628 | 232 | Unknown |
| | 41.5 | -24.5 | 63.5 | 2.625 | 912 | Postcentral Gyrus |
| | 37.5 | -18.5 | 55.5 | 2.172 | | Precentral Gyrus |
| | 27.5 | 31.5 | -2.5 | 2.625 | 240 | Unknown |
| | 19.5 | -24.5 | 81.5 | 2.606 | 272 | Unknown |
| | -22.5 | -46.5 | 75.5 | 2.591 | 640 | Unknown |
| | -20.5 | -42.5 | 67.5 | 2.475 | | Postcentral Gyrus |
| | 17.5 | -96.5 | -4.5 | 2.493 | 192 | Occipital Pole |
| | -48.5 | -30.5 | 9.5 | 2.468 | 312 | Planum Temporale |
| | -54.5 | -26.5 | 5.5 | 2.338 | | Planum Temporale |
| | -0.5 | -52.5 | -60.5 | 2.461 | 648 | Unknown |
| | 15.5 | -6.5 | 63.5 | 2.445 | 168 | Unknown |
| | -0.5 | 19.5 | -6.5 | 2.333 | 160 | Subcallosal Cortex |
| | -2.5 | -70.5 | -44.5 | 2.322 | 208 | Unknown |
| | 7.5 | -74.5 | -14.5 | 2.304 | 352 | Unknown |
| | 13.5 | -64.5 | -14.5 | 2.197 | | Unknown |
| | -14.5 | -30.5 | 71.5 | 2.255 | 176 | Precentral Gyrus |
| | 7.5 | -90.5 | -20.5 | 2.235 | 192 | Unknown |
| Negative | 57.5 | -56.5 | 39.5 | 4.649 | 10864 | Angular Gyrus |
| | 59.5 | -56.5 | 29.5 | 3.703 | | Angular Gyrus |
| | 55.5 | -46.5 | 53.5 | 3.221 | | Angular Gyrus |
| | 45.5 | -58.5 | 33.5 | 3.079 | | Unknown |
| | -58.5 | -60.5 | 29.5 | 4.055 | 9376 | Angular Gyrus |
| | -48.5 | -62.5 | 43.5 | 3.971 | | Lateral Occipital Cortex, superior division |
| | -52.5 | -58.5 | 37.5 | 3.574 | | Angular Gyrus |
| | -52.5 | -50.5 | 39.5 | 3.423 | | Supramarginal Gyrus, posterior division |
| | 29.5 | 33.5 | 37.5 | 3.687 | 30576 | Middle Frontal Gyrus |
| | 31.5 | 25.5 | 37.5 | 3.396 | | Middle Frontal Gyrus |
| | 17.5 | 27.5 | 55.5 | 3.293 | | Superior Frontal Gyrus |
| | 13.5 | 25.5 | 65.5 | 3.259 | | Unknown |
| | -0.5 | -22.5 | 39.5 | 3.479 | 3360 | Cingulate Gyrus, posterior division |
| | 69.5 | -40.5 | 1.5 | 3.453 | 2368 | Middle Temporal Gyrus, temporooccipital part |
| | 67.5 | -12.5 | -16.5 | 3.168 | | Middle Temporal Gyrus, posterior division |
| | 65.5 | -34.5 | -4.5 | 2.934 | | Middle Temporal Gyrus, posterior division |
| | 65.5 | -20.5 | -10.5 | 2.44 | | Middle Temporal Gyrus, posterior division |
| | 45.5 | -2.5 | -38.5 | 3.128 | 2176 | Inferior Temporal Gyrus, anterior division |
| | 55.5 | 5.5 | -34.5 | 2.999 | | Temporal Pole |
| | 55.5 | -8.5 | -26.5 | 2.435 | | Unknown |
| | 47.5 | 5.5 | -46.5 | 2.062 | | Unknown |
| | -46.5 | 1.5 | -42.5 | 3.078 | 528 | Temporal Pole |
| | 13.5 | -54.5 | 37.5 | 3.043 | 2928 | Precuneous Cortex |
| | 15.5 | -46.5 | 33.5 | 3.009 | | Cingulate Gyrus, posterior division |
| | 5.5 | -38.5 | 29.5 | 2.298 | | Cingulate Gyrus, posterior division |
| | -58.5 | -8.5 | -26.5 | 2.993 | 936 | Middle Temporal Gyrus, anterior division |
| | -64.5 | -16.5 | -22.5 | 2.797 | | Middle Temporal Gyrus, posterior division |
| | -66.5 | -24.5 | -26.5 | 2.287 | | Unknown |
| | -42.5 | 11.5 | 49.5 | 2.872 | 2296 | Middle Frontal Gyrus |
| | -44.5 | 19.5 | 43.5 | 2.64 | | Middle Frontal Gyrus |
| | -40.5 | 5.5 | 43.5 | 2.401 | | Middle Frontal Gyrus |
| | -36.5 | 27.5 | 45.5 | 2.363 | | Middle Frontal Gyrus |
| | 7.5 | -6.5 | 19.5 | 2.773 | 648 | Unknown |
| | 9.5 | 5.5 | 17.5 | 2.044 | | Unknown |
| | -64.5 | -30.5 | -4.5 | 2.669 | 1184 | Middle Temporal Gyrus, posterior division |
| | -64.5 | -42.5 | -4.5 | 2.332 | | Middle Temporal Gyrus, posterior division |
| | 17.5 | 57.5 | 27.5 | 2.485 | 392 | Frontal Pole |
| | -10.5 | -40.5 | 29.5 | 2.48 | 632 | Unknown |
| | -10.5 | -30.5 | 23.5 | 2.399 | 192 | Unknown |

*Table S6. Clusters obtained at a threshold > 2 for the "correct" contrast in the Learn task and the prediction of the Verbal summary score.*

| Sign | MNI coordinates (mm) | | | Peak Stat | Cluster Size ($mm^3$) | Label |
|---|---|---|---|---|---|---|
| | X | Y | Z | | | |
| Positive | -40.5 | -20.5 | 53.5 | 3.992 | 18656 | Precentral Gyrus |
| | -40.5 | -24.5 | 65.5 | 3.848 | | Postcentral Gyrus |
| | -30.5 | -6.5 | 51.5 | 3.645 | | Precentral Gyrus |
| | -32.5 | -26.5 | 61.5 | 3.572 | | Postcentral Gyrus |
| | 5.5 | 21.5 | 35.5 | 3.942 | 15728 | Paracingulate Gyrus |
| | -2.5 | 7.5 | 55.5 | 3.742 | | Juxtapositional Lobule Cortex (formerly Supplementary Motor Cortex) |
| | -6.5 | 1.5 | 59.5 | 3.645 | | Juxtapositional Lobule Cortex (formerly Supplementary Motor Cortex) |
| | -10.5 | 15.5 | 37.5 | 3.558 | | Paracingulate Gyrus |
| | -36.5 | -0.5 | 35.5 | 3.597 | 3280 | Unknown |
| | -46.5 | 1.5 | 31.5 | 3.101 | | Precentral Gyrus |
| | -54.5 | 9.5 | 39.5 | 2.663 | | Unknown |
| | -10.5 | -86.5 | 1.5 | 3.596 | 4312 | Intracalcarine Cortex |
| | 7.5 | -76.5 | 11.5 | 3.332 | | Intracalcarine Cortex |
| | -14.5 | -80.5 | 7.5 | 3.224 | | Intracalcarine Cortex |
| | -46.5 | -0.5 | 7.5 | 3.51 | 2296 | Central Opercular Cortex |
| | -42.5 | -4.5 | 15.5 | 3.418 | | Central Opercular Cortex |
| | -26.5 | -90.5 | 29.5 | 3.435 | 4032 | Occipital Pole |
| | -26.5 | -82.5 | 17.5 | 3.266 | | Lateral Occipital Cortex, superior division |
| | -22.5 | -68.5 | 45.5 | 3.062 | | Lateral Occipital Cortex, superior division |
| | -26.5 | -84.5 | 39.5 | 2.439 | | Lateral Occipital Cortex, superior division |
| | -32.5 | 19.5 | 11.5 | 3.381 | 3096 | Frontal Opercular Cortex |
| | 43.5 | -36.5 | 45.5 | 3.057 | 480 | Supramarginal Gyrus, posterior division |
| | -46.5 | -64.5 | -14.5 | 2.928 | 792 | Inferior Temporal Gyrus, temporooccipital part |
| | 27.5 | -4.5 | 51.5 | 2.853 | 1552 | Unknown |
| | 23.5 | -70.5 | 45.5 | 2.816 | 3672 | Lateral Occipital Cortex, superior division |
| | 27.5 | -54.5 | 43.5 | 2.807 | | Superior Parietal Lobule |
| | 25.5 | -74.5 | 57.5 | 2.552 | | Lateral Occipital Cortex, superior division |
| | 29.5 | -76.5 | 35.5 | 2.419 | | Lateral Occipital Cortex, superior division |
| | 29.5 | -82.5 | 17.5 | 2.726 | 320 | Lateral Occipital Cortex, superior division |
| | -26.5 | -58.5 | -10.5 | 2.647 | 712 | Temporal Occipital Fusiform Cortex |
| | -24.5 | -70.5 | -6.5 | 2.424 | | Occipital Fusiform Gyrus |
| | 31.5 | 17.5 | 11.5 | 2.598 | 440 | Frontal Opercular Cortex |
| | -30.5 | -92.5 | -10.5 | 2.546 | 464 | Occipital Pole |
| | 15.5 | -12.5 | -28.5 | 2.485 | 216 | Parahippocampal Gyrus, anterior division |
| | 41.5 | 5.5 | 27.5 | 2.478 | 288 | Precentral Gyrus |
| | 21.5 | -2.5 | -34.5 | 2.425 | 392 | Parahippocampal Gyrus, anterior division |
| | 13.5 | -74.5 | -10.5 | 2.424 | 176 | Lingual Gyrus |
| | -46.5 | -52.5 | -20.5 | 2.407 | 208 | Inferior Temporal Gyrus, temporooccipital part |
| | 41.5 | -86.5 | 3.5 | 2.377 | 304 | Lateral Occipital Cortex, inferior division |
| | 41.5 | -88.5 | -6.5 | 2.361 | | Lateral Occipital Cortex, inferior division |
| | -38.5 | -84.5 | -6.5 | 2.356 | 264 | Lateral Occipital Cortex, inferior division |
| | 31.5 | 21.5 | -4.5 | 2.342 | 248 | Insular Cortex |
| | 47.5 | -72.5 | -12.5 | 2.325 | 176 | Lateral Occipital Cortex, inferior division |
| | 47.5 | -64.5 | -14.5 | 2.04 | | Lateral Occipital Cortex, inferior division |
| Negative | 15.5 | -96.5 | -4.5 | 3.643 | 4992 | Occipital Pole |
| | 15.5 | -82.5 | -28.5 | 2.977 | | Unknown |
| | 13.5 | -80.5 | -42.5 | 2.565 | | Unknown |
| | 23.5 | -74.5 | -34.5 | 2.486 | | Unknown |
| | -56.5 | 21.5 | 5.5 | 3.49 | 2952 | Inferior Frontal Gyrus, pars triangularis |
| | -42.5 | 23.5 | -12.5 | 2.945 | | Frontal Orbital Cortex |
| | -48.5 | 27.5 | -2.5 | 2.529 | | Inferior Frontal Gyrus, pars triangularis |
| | -48.5 | 31.5 | -12.5 | 2.241 | | Frontal Orbital Cortex |
| | -60.5 | -0.5 | -16.5 | 3.423 | 2856 | Middle Temporal Gyrus, anterior division |
| | -62.5 | -14.5 | -8.5 | 2.713 | | Middle Temporal Gyrus, posterior division |
| | -52.5 | -6.5 | -16.5 | 2.537 | | Middle Temporal Gyrus, anterior division |
| | -50.5 | -16.5 | -6.5 | 2.455 | | Unknown |
| | -16.5 | 59.5 | 25.5 | 3.234 | 2688 | Frontal Pole |
| | -16.5 | 61.5 | 15.5 | 2.878 | | Frontal Pole |
| | -12.5 | 55.5 | 35.5 | 2.449 | | Frontal Pole |
| | -20.5 | 67.5 | 7.5 | 2.339 | | Frontal Pole |
| | 37.5 | -18.5 | 21.5 | 3.054 | 888 | Central Opercular Cortex |
| | 41.5 | -32.5 | 67.5 | 2.957 | 5216 | Postcentral Gyrus |
| | 31.5 | -28.5 | 69.5 | 2.821 | | Postcentral Gyrus |
| | 23.5 | -26.5 | 61.5 | 2.786 | | Precentral Gyrus |
| | 25.5 | -26.5 | 73.5 | 2.643 | | Precentral Gyrus |
| | -4.5 | -82.5 | -8.5 | 2.891 | 816 | Lingual Gyrus |
| | 5.5 | -84.5 | -8.5 | 2.816 | | Lingual Gyrus |
| | -16.5 | -98.5 | 5.5 | 2.668 | 816 | Occipital Pole |
| | -12.5 | -98.5 | -4.5 | 2.484 | | Occipital Pole |
| | 7.5 | -52.5 | -42.5 | 2.667 | 232 | Unknown |
| | 35.5 | -48.5 | -42.5 | 2.593 | 176 | Unknown |
| | 49.5 | -10.5 | 49.5 | 2.555 | 496 | Precentral Gyrus |
| | 47.5 | -10.5 | 61.5 | 2.204 | | Unknown |
| | -62.5 | -46.5 | -0.5 | 2.544 | 944 | Middle Temporal Gyrus, temporooccipital part |
| | -64.5 | -54.5 | 3.5 | 2.44 | | Middle Temporal Gyrus, temporooccipital part |
| | 49.5 | -34.5 | 25.5 | 2.525 | 320 | Parietal Opercular Cortex |
| | -42.5 | -64.5 | 31.5 | 2.5 | 1384 | Lateral Occipital Cortex, superior division |
| | -54.5 | -60.5 | 35.5 | 2.444 | | Lateral Occipital Cortex, superior division |
| | -52.5 | -68.5 | 29.5 | 2.346 | | Lateral Occipital Cortex, superior division |
| | -54.5 | -60.5 | 23.5 | 2.315 | | Angular Gyrus |
| | -10.5 | 41.5 | -8.5 | 2.464 | 224 | Unknown |
| | -4.5 | 47.5 | 13.5 | 2.413 | 696 | Paracingulate Gyrus |
| | -2.5 | 55.5 | 9.5 | 2.399 | | Paracingulate Gyrus |
| | -4.5 | 65.5 | 19.5 | 2.403 | 552 | Frontal Pole |
| | -2.5 | 57.5 | 27.5 | 2.27 | | Frontal Pole |

| | | | | | | |
|---|---|---|---|---|---|---|
| | 31.5 | -72.5 | 1.5 | 2.402 | 320 | Unknown |
| | -14.5 | 31.5 | 61.5 | 2.374 | 296 | Unknown |
| | -46.5 | 5.5 | -32.5 | 2.307 | 160 | Temporal Pole |
| | 57.5 | -0.5 | -20.5 | 2.26 | 168 | Middle Temporal Gyrus, anterior division |
| | -58.5 | -34.5 | -0.5 | 2.23 | 336 | Middle Temporal Gyrus, posterior division |
| | 39.5 | -14.5 | 39.5 | 2.177 | 168 | Precentral Gyrus |
| | 45.5 | -8.5 | 37.5 | 2.135 | | Precentral Gyrus |

*Table S7. Clusters obtained at a threshold > 2 for the pseudohomophone contrast in the PhonLex task and the prediction of the Naming summary score.*

| Sign | MNI coordinates (mm) X | Y | Z | Peak Stat | Cluster Size ($mm^3$) | Label |
|---|---|---|---|---|---|---|
| Positive | 21.5 | -92.5 | -10.5 | 3.213 | 1344 | Occipital Pole |
| | 31.5 | -30.5 | 27.5 | 3.119 | 1184 | Unknown |
| | 33.5 | -20.5 | 27.5 | 3.039 | | Unknown |
| | -18.5 | 55.5 | -16.5 | 3.084 | 280 | Frontal Pole |
| | 11.5 | 1.5 | 25.5 | 3.077 | 720 | Unknown |
| | 7.5 | 5.5 | 17.5 | 2.598 | | Unknown |
| | -54.5 | 7.5 | 27.5 | 3.025 | 3664 | Precentral Gyrus |
| | -56.5 | 11.5 | 19.5 | 2.634 | | Inferior Frontal Gyrus, pars opercularis |
| | -60.5 | 3.5 | 35.5 | 2.601 | | Unknown |
| | -54.5 | -4.5 | 27.5 | 2.541 | | Precentral Gyrus |
| | 41.5 | -0.5 | 21.5 | 2.99 | 416 | Unknown |
| | 7.5 | -42.5 | -36.5 | 2.916 | 576 | Unknown |
| | 29.5 | 9.5 | -40.5 | 2.897 | 560 | Temporal Pole |
| | -38.5 | -2.5 | 35.5 | 2.856 | 792 | Precentral Gyrus |
| | -46.5 | -0.5 | 37.5 | 2.421 | | Precentral Gyrus |
| | -28.5 | 3.5 | 35.5 | 2.247 | | Unknown |
| | -18.5 | -24.5 | 7.5 | 2.802 | 1512 | Unknown |
| | -6.5 | -24.5 | 19.5 | 2.646 | | Unknown |
| | -8.5 | -22.5 | 7.5 | 2.277 | | Unknown |
| | 9.5 | -26.5 | 21.5 | 2.79 | 816 | Unknown |
| | 19.5 | -28.5 | 25.5 | 2.698 | | Unknown |
| | -20.5 | -90.5 | -14.5 | 2.763 | 368 | Occipital Fusiform Gyrus |
| | 53.5 | 29.5 | 19.5 | 2.759 | 656 | Inferior Frontal Gyrus, pars triangularis |
| | 45.5 | 35.5 | 17.5 | 2.48 | | Frontal Pole |
| | -44.5 | -68.5 | -12.5 | 2.677 | 824 | Lateral Occipital Cortex, inferior division |
| | -40.5 | -76.5 | -10.5 | 2.536 | | Lateral Occipital Cortex, inferior division |
| | 39.5 | -2.5 | -40.5 | 2.657 | 496 | Inferior Temporal Gyrus, anterior division |
| | 31.5 | -2.5 | -42.5 | 2.621 | | Temporal Fusiform Cortex, anterior division |
| | 49.5 | -2.5 | -36.5 | 2.493 | | Inferior Temporal Gyrus, anterior division |
| | -22.5 | 21.5 | 31.5 | 2.649 | 584 | Unknown |
| | -8.5 | 51.5 | -26.5 | 2.597 | 192 | Frontal Pole |
| | -0.5 | -54.5 | -34.5 | 2.594 | 184 | Unknown |
| | 51.5 | -0.5 | 9.5 | 2.593 | 1008 | Central Opercular Cortex |
| | 43.5 | 1.5 | 7.5 | 2.463 | | Central Opercular Cortex |
| | 39.5 | 9.5 | -4.5 | 2.158 | | Insular Cortex |
| | -6.5 | 19.5 | -28.5 | 2.58 | 336 | Unknown |
| | 21.5 | -26.5 | 43.5 | 2.55 | 208 | Unknown |
| | -12.5 | 31.5 | -20.5 | 2.526 | 176 | Frontal Orbital Cortex |
| | -4.5 | -16.5 | -46.5 | 2.499 | 192 | Unknown |
| | -56.5 | -18.5 | 45.5 | 2.486 | 208 | Postcentral Gyrus |
| | -14.5 | -8.5 | -36.5 | 2.437 | 160 | Unknown |
| | -4.5 | 1.5 | 57.5 | 2.432 | 208 | Juxtapositional Lobule Cortex (formerly Supplementary Motor Cortex) |
| | -2.5 | -10.5 | 1.5 | 2.42 | 176 | Unknown |
| | 11.5 | 37.5 | 53.5 | 2.389 | 208 | Frontal Pole |
| | 1.5 | -4.5 | 29.5 | 2.359 | 376 | Cingulate Gyrus, anterior division |
| | -34.5 | 1.5 | -38.5 | 2.304 | 304 | Temporal Pole |
| | -20.5 | 1.5 | 47.5 | 2.297 | 536 | Unknown |
| | -12.5 | 5.5 | 43.5 | 2.292 | | Unknown |
| | -18.5 | -8.5 | 47.5 | 2.26 | | Unknown |
| | -24.5 | -0.5 | 49.5 | 2.175 | | Middle Frontal Gyrus |
| Negative | -0.5 | -92.5 | 5.5 | 5.078 | 61144 | Occipital Pole |
| | 5.5 | -84.5 | 33.5 | 4.308 | | Cuneal Cortex |
| | -2.5 | -90.5 | 19.5 | 4.081 | | Occipital Pole |
| | 13.5 | -82.5 | -0.5 | 3.931 | | Intracalcarine Cortex |
| | -0.5 | 35.5 | 65.5 | 3.277 | 264 | Unknown |
| | 31.5 | 47.5 | -0.5 | 3.267 | 776 | Unknown |
| | 33.5 | 35.5 | 1.5 | 2.362 | | Unknown |
| | 49.5 | 19.5 | 49.5 | 3.088 | 760 | Unknown |
| | 45.5 | 29.5 | 45.5 | 2.88 | | Unknown |
| | -40.5 | 27.5 | 49.5 | 3.057 | 560 | Unknown |
| | -0.5 | 57.5 | 45.5 | 3.012 | 440 | Unknown |
| | 9.5 | 59.5 | 41.5 | 2.749 | | Unknown |
| | 59.5 | -26.5 | 5.5 | 2.902 | 704 | Unknown |
| | 65.5 | -32.5 | 9.5 | 2.568 | | Superior Temporal Gyrus, posterior division |
| | 35.5 | -46.5 | 67.5 | 2.665 | 976 | Superior Parietal Lobule |
| | 29.5 | -38.5 | 73.5 | 2.614 | | Unknown |
| | 29.5 | -44.5 | 57.5 | 2.499 | | Superior Parietal Lobule |
| | 3.5 | -66.5 | 63.5 | 2.648 | 448 | Unknown |
| | -14.5 | 7.5 | 77.5 | 2.599 | 160 | Unknown |
| | -34.5 | 59.5 | -0.5 | 2.579 | 208 | Frontal Pole |
| | 3.5 | -64.5 | -60.5 | 2.555 | 472 | Unknown |
| | 3.5 | -74.5 | -48.5 | 2.476 | | Unknown |
| | 41.5 | -16.5 | 61.5 | 2.529 | 896 | Precentral Gyrus |
| | 45.5 | -28.5 | 65.5 | 2.46 | | Postcentral Gyrus |
| | 35.5 | -26.5 | 67.5 | 2.401 | | Precentral Gyrus |
| | 45.5 | -22.5 | 63.5 | 2.316 | | Postcentral Gyrus |
| | 37.5 | 7.5 | 55.5 | 2.485 | 280 | Middle Frontal Gyrus |

*Table S8. Clusters obtained at a threshold > 2 for the FFnew contrast in the CharProc task and the prediction of the Naming summary score.*

| Sign | MNI coordinates (mm) X | Y | Z | Peak Stat | Cluster Size ($mm^3$) | Label |
|---|---|---|---|---|---|---|
| Positive | -50.5 | -80.5 | -0.5 | 4.009 | 4256 | Lateral Occipital Cortex, inferior division |
| | -40.5 | -70.5 | -4.5 | 3.244 | | Lateral Occipital Cortex, inferior division |
| | -44.5 | -56.5 | -6.5 | 2.518 | | Inferior Temporal Gyrus, temporooccipital part |
| | -54.5 | -66.5 | 1.5 | 2.411 | | Lateral Occipital Cortex, inferior division |
| | 3.5 | 59.5 | -26.5 | 3.765 | 704 | Frontal Pole |
| | 5.5 | 49.5 | -26.5 | 3.394 | | Frontal Medial Cortex |
| | 63.5 | -4.5 | -26.5 | 3.575 | 1720 | Middle Temporal Gyrus, anterior division |
| | 53.5 | -2.5 | -40.5 | 3.473 | | Inferior Temporal Gyrus, anterior division |
| | 57.5 | -18.5 | -36.5 | 2.549 | | Inferior Temporal Gyrus, posterior division |
| | 47.5 | 11.5 | -38.5 | 2.447 | | Temporal Pole |
| | 53.5 | -72.5 | -2.5 | 3.145 | 1448 | Lateral Occipital Cortex, inferior division |
| | 51.5 | -62.5 | -10.5 | 2.764 | | Lateral Occipital Cortex, inferior division |
| | 53.5 | -60.5 | 1.5 | 2.475 | | Lateral Occipital Cortex, inferior division |
| | 49.5 | -64.5 | 7.5 | 2.253 | | Lateral Occipital Cortex, inferior division |
| | -38.5 | -8.5 | -46.5 | 3.046 | 256 | Unknown |
| | 43.5 | -0.5 | -28.5 | 2.989 | 312 | Unknown |
| | -12.5 | -16.5 | -26.5 | 2.82 | 440 | Unknown |
| | 13.5 | -38.5 | -40.5 | 2.808 | 280 | Unknown |
| | 21.5 | -42.5 | -44.5 | 2.379 | | Unknown |
| | -44.5 | 13.5 | -32.5 | 2.803 | 392 | Temporal Pole |
| | 39.5 | -48.5 | -8.5 | 2.8 | 200 | Unknown |
| | 15.5 | -0.5 | -26.5 | 2.792 | 224 | Parahippocampal Gyrus, anterior division |
| | 47.5 | 3.5 | 33.5 | 2.792 | 1136 | Precentral Gyrus |
| | 51.5 | 9.5 | 41.5 | 2.416 | | Middle Frontal Gyrus |
| | -36.5 | -8.5 | 49.5 | 2.781 | 1336 | Precentral Gyrus |
| | -24.5 | -4.5 | 51.5 | 2.647 | | Middle Frontal Gyrus |
| | -34.5 | -0.5 | 53.5 | 2.243 | | Middle Frontal Gyrus |
| | -28.5 | 25.5 | -24.5 | 2.739 | 584 | Frontal Orbital Cortex |
| | -34.5 | 13.5 | -36.5 | 2.731 | 192 | Temporal Pole |
| | 31.5 | -36.5 | -24.5 | 2.722 | 792 | Temporal Fusiform Cortex, posterior division |
| | 33.5 | -28.5 | -28.5 | 2.643 | | Temporal Fusiform Cortex, posterior division |
| | 37.5 | -22.5 | -24.5 | 2.532 | | Temporal Fusiform Cortex, posterior division |
| | -0.5 | 27.5 | -22.5 | 2.711 | 176 | Subcallosal Cortex |
| | -8.5 | -76.5 | -48.5 | 2.704 | 280 | Unknown |
| | -58.5 | -8.5 | -28.5 | 2.689 | 192 | Middle Temporal Gyrus, anterior division |
| | -42.5 | -40.5 | -28.5 | 2.683 | 424 | Temporal Fusiform Cortex, posterior division |
| | -48.5 | -32.5 | -24.5 | 2.496 | | Inferior Temporal Gyrus, posterior division |
| | 29.5 | -50.5 | 51.5 | 2.65 | 304 | Superior Parietal Lobule |
| | -50.5 | 1.5 | 29.5 | 2.643 | 304 | Precentral Gyrus |
| | -26.5 | 13.5 | 47.5 | 2.619 | 568 | Middle Frontal Gyrus |
| | -48.5 | -52.5 | 9.5 | 2.579 | 392 | Middle Temporal Gyrus, temporooccipital part |
| | 53.5 | -26.5 | 49.5 | 2.497 | 544 | Postcentral Gyrus |
| | 57.5 | -24.5 | 39.5 | 2.257 | | Supramarginal Gyrus, anterior division |
| | -14.5 | 29.5 | 53.5 | 2.484 | 192 | Superior Frontal Gyrus |
| | -6.5 | 31.5 | 55.5 | 2.197 | | Superior Frontal Gyrus |
| | 65.5 | -12.5 | 35.5 | 2.439 | 160 | Postcentral Gyrus |
| | -22.5 | -16.5 | -34.5 | 2.423 | 224 | Parahippocampal Gyrus, anterior division |
| | -32.5 | -10.5 | -36.5 | 2.153 | | Temporal Fusiform Cortex, posterior division |
| | -6.5 | 45.5 | -2.5 | 2.339 | 160 | Paracingulate Gyrus |
| Negative | 17.5 | -96.5 | 21.5 | 5.073 | 58848 | Occipital Pole |
| | 3.5 | -84.5 | 39.5 | 4.799 | | Cuneal Cortex |
| | -16.5 | -88.5 | -8.5 | 4.769 | | Unknown |
| | -8.5 | -102.5 | 3.5 | 4.675 | | Occipital Pole |
| | 17.5 | -26.5 | 81.5 | 3.468 | 1744 | Unknown |
| | 17.5 | -4.5 | 79.5 | 2.877 | | Unknown |
| | -24.5 | -24.5 | 79.5 | 3.401 | 656 | Unknown |
| | -28.5 | -32.5 | 73.5 | 3.036 | | Postcentral Gyrus |
| | 49.5 | -2.5 | 57.5 | 3.189 | 392 | Unknown |
| | 51.5 | -12.5 | 59.5 | 2.29 | | Unknown |
| | 5.5 | 39.5 | 61.5 | 3.064 | 392 | Unknown |
| | 17.5 | 37.5 | 59.5 | 2.605 | | Unknown |
| | 63.5 | -10.5 | -4.5 | 2.959 | 256 | Unknown |
| | 69.5 | -16.5 | -18.5 | 2.811 | 832 | Middle Temporal Gyrus, posterior division |
| | 55.5 | -20.5 | -16.5 | 2.81 | | Unknown |
| | -24.5 | 9.5 | 71.5 | 2.788 | 384 | Unknown |
| | -34.5 | 7.5 | 65.5 | 2.615 | | Unknown |
| | -28.5 | 17.5 | 65.5 | 2.205 | | Unknown |
| | -8.5 | -44.5 | 53.5 | 2.643 | 344 | Precuneous Cortex |
| | -50.5 | -52.5 | -30.5 | 2.632 | 536 | Unknown |
| | -56.5 | -62.5 | -30.5 | 2.618 | | Unknown |
| | 3.5 | 15.5 | 67.5 | 2.62 | 240 | Unknown |
| | 7.5 | 21.5 | 71.5 | 2.254 | | Unknown |
| | 23.5 | -54.5 | -2.5 | 2.609 | 168 | Lingual Gyrus |
| | -14.5 | -2.5 | -16.5 | 2.595 | 168 | Unknown |
| | 25.5 | 21.5 | 65.5 | 2.505 | 368 | Unknown |
| | 11.5 | 31.5 | 65.5 | 2.487 | | Unknown |
| | 53.5 | 37.5 | -8.5 | 2.346 | 360 | Frontal Pole |
| | 43.5 | 41.5 | -14.5 | 2.261 | | Frontal Pole |

*Table S9. Clusters obtained at a threshold > 2 for the "incorrect" contrast in the Learn task and the prediction of the Naming summary score.*

| Sign | MNI coordinates (mm) X | Y | Z | Peak Stat | Cluster Size ($mm^3$) | Label |
|---|---|---|---|---|---|---|
| Positive | -34.5 | -26.5 | 51.5 | 4.161 | 4760 | Postcentral Gyrus |
| | -52.5 | -22.5 | 49.5 | 3.08 | | Postcentral Gyrus |
| | -46.5 | -18.5 | 61.5 | 2.952 | | Postcentral Gyrus |
| | -40.5 | -36.5 | 53.5 | 2.259 | | Postcentral Gyrus |
| | -16.5 | -84.5 | -8.5 | 3.655 | 5872 | Unknown |
| | -10.5 | -86.5 | 1.5 | 3.628 | | Intracalcarine Cortex |
| | -42.5 | -82.5 | -14.5 | 3.361 | | Lateral Occipital Cortex, inferior division |
| | -28.5 | -96.5 | -10.5 | 3.2 | | Occipital Pole |
| | -8.5 | 21.5 | 47.5 | 3.635 | 8056 | Unknown |
| | 7.5 | 19.5 | 43.5 | 3.622 | | Paracingulate Gyrus |
| | -4.5 | 13.5 | 51.5 | 3.439 | | Paracingulate Gyrus |
| | -10.5 | 1.5 | 63.5 | 2.517 | | Superior Frontal Gyrus |
| | -30.5 | 19.5 | 13.5 | 3.425 | 3744 | Frontal Opercular Cortex |
| | -30.5 | 17.5 | -6.5 | 2.932 | | Insular Cortex |
| | -46.5 | 11.5 | 3.5 | 2.351 | | Inferior Frontal Gyrus, pars opercularis |
| | -40.5 | 15.5 | -0.5 | 2.16 | | Frontal Opercular Cortex |
| | -58.5 | -44.5 | 15.5 | 3.321 | 1088 | Supramarginal Gyrus, posterior division |
| | -64.5 | -38.5 | 7.5 | 2.575 | | Superior Temporal Gyrus, posterior division |
| | 13.5 | -76.5 | -10.5 | 3.305 | 640 | Lingual Gyrus |
| | 47.5 | -82.5 | -0.5 | 3.302 | 1024 | Lateral Occipital Cortex, inferior division |
| | 51.5 | -78.5 | 5.5 | 3.022 | | Lateral Occipital Cortex, inferior division |
| | 49.5 | -72.5 | -12.5 | 2.352 | | Lateral Occipital Cortex, inferior division |
| | -38.5 | -2.5 | 55.5 | 3.266 | 1360 | Middle Frontal Gyrus |
| | -32.5 | -14.5 | 55.5 | 2.137 | | Precentral Gyrus |
| | -8.5 | -6.5 | 53.5 | 3.254 | 512 | Juxtapositional Lobule Cortex (formerly Supplementary Motor Cortex) |
| | -32.5 | -52.5 | -16.5 | 3.2 | 1088 | Temporal Occipital Fusiform Cortex |
| | -26.5 | -62.5 | -12.5 | 2.627 | | Temporal Occipital Fusiform Cortex |
| | -34.5 | -94.5 | 9.5 | 3.175 | 480 | Occipital Pole |
| | -36.5 | 5.5 | 35.5 | 3.099 | 1624 | Middle Frontal Gyrus |
| | -46.5 | 1.5 | 31.5 | 2.82 | | Precentral Gyrus |
| | 33.5 | 57.5 | -16.5 | 2.98 | 2664 | Frontal Pole |
| | 31.5 | 55.5 | -6.5 | 2.913 | | Frontal Pole |
| | 41.5 | 53.5 | -10.5 | 2.903 | | Frontal Pole |
| | 25.5 | 65.5 | -12.5 | 2.555 | | Frontal Pole |
| | -30.5 | 13.5 | -32.5 | 2.863 | 424 | Temporal Pole |
| | -8.5 | -32.5 | -20.5 | 2.855 | 3064 | Unknown |
| | 7.5 | -26.5 | -18.5 | 2.679 | | Unknown |
| | 11.5 | -42.5 | -10.5 | 2.612 | | Unknown |
| | 7.5 | -50.5 | -20.5 | 2.456 | | Unknown |
| | 27.5 | -54.5 | -20.5 | 2.839 | 360 | Unknown |
| | 13.5 | -0.5 | -20.5 | 2.825 | 312 | Unknown |
| | -20.5 | -96.5 | 19.5 | 2.798 | 216 | Occipital Pole |
| | 67.5 | -10.5 | -2.5 | 2.766 | 360 | Superior Temporal Gyrus, posterior division |
| | 69.5 | -20.5 | -4.5 | 2.57 | | Middle Temporal Gyrus, posterior division |
| | 11.5 | 17.5 | -20.5 | 2.754 | 520 | Subcallosal Cortex |
| | -12.5 | 63.5 | -16.5 | 2.725 | 288 | Frontal Pole |
| | 45.5 | 13.5 | 9.5 | 2.667 | 776 | Unknown |
| | -10.5 | -4.5 | -24.5 | 2.603 | 632 | Parahippocampal Gyrus, anterior division |
| | -6.5 | 7.5 | -20.5 | 2.326 | | Unknown |
| | -30.5 | -26.5 | -24.5 | 2.547 | 248 | Parahippocampal Gyrus, posterior division |
| | 29.5 | 23.5 | -6.5 | 2.537 | 1136 | Frontal Orbital Cortex |
| | 43.5 | 19.5 | -4.5 | 2.3 | | Frontal Orbital Cortex |
| | 37.5 | 31.5 | 1.5 | 2.263 | | Unknown |
| | 39.5 | 17.5 | 1.5 | 2.096 | | Insular Cortex |
| | 49.5 | -20.5 | -0.5 | 2.531 | 560 | Unknown |
| | 49.5 | -28.5 | 3.5 | 2.116 | | Unknown |
| | 3.5 | 45.5 | 39.5 | 2.413 | 256 | Superior Frontal Gyrus |
| Negative | 45.5 | -12.5 | 47.5 | 4.214 | 36072 | Precentral Gyrus |
| | 37.5 | -14.5 | 43.5 | 4.058 | | Precentral Gyrus |
| | 51.5 | -32.5 | 23.5 | 3.997 | | Parietal Opercular Cortex |
| | 21.5 | -28.5 | 63.5 | 3.949 | | Precentral Gyrus |
| | -16.5 | -24.5 | 75.5 | 3.836 | 3776 | Precentral Gyrus |
| | -26.5 | -36.5 | 59.5 | 2.591 | | Postcentral Gyrus |
| | 13.5 | -84.5 | 37.5 | 3.412 | 824 | Lateral Occipital Cortex, superior division |
| | -26.5 | -60.5 | -60.5 | 3.059 | 248 | Unknown |
| | 29.5 | -30.5 | 47.5 | 3.035 | 432 | Unknown |
| | -56.5 | -6.5 | 33.5 | 2.878 | 3072 | Precentral Gyrus |
| | -58.5 | -8.5 | 25.5 | 2.87 | | Postcentral Gyrus |
| | -58.5 | -10.5 | 13.5 | 2.865 | | Central Opercular Cortex |
| | -60.5 | -2.5 | 23.5 | 2.823 | | Precentral Gyrus |
| | 31.5 | -72.5 | 5.5 | 2.738 | 296 | Unknown |
| | 11.5 | -30.5 | 45.5 | 2.736 | 1088 | Precentral Gyrus |
| | 49.5 | -58.5 | -2.5 | 2.682 | 704 | Unknown |
| | -22.5 | -66.5 | -44.5 | 2.628 | 488 | Unknown |
| | -22.5 | -10.5 | -0.5 | 2.615 | 184 | Unknown |
| | -10.5 | -88.5 | 35.5 | 2.505 | 584 | Unknown |
| | 47.5 | -4.5 | -6.5 | 2.505 | 160 | Planum Polare |
| | -8.5 | -56.5 | 9.5 | 2.412 | 232 | Precuneous Cortex |
| | 23.5 | -68.5 | -40.5 | 2.374 | 176 | Unknown |
| | -32.5 | -38.5 | 9.5 | 2.37 | 496 | Unknown |
| | -34.5 | -48.5 | 1.5 | 2.287 | | Unknown |
| | 41.5 | -78.5 | 33.5 | 2.358 | 208 | Lateral Occipital Cortex, superior division |
| | -34.5 | -30.5 | 31.5 | 2.246 | 192 | Unknown |

*Table S10. Top ten associations of consensus maps, for positive and negative values, and meta-analytical terms for the prediction of the summary score Reading.*

| Task and contrast | Positive | Negative |
|---|---|---|
| Task: PhonLex, contrast: pseudohomophone | #1: taste_swallowing_gustatory<br>#2: pain_painful_chronic<br>#3: anterior_insula_cortex<br>#4: empathy_social_empathic<br>#5: thalamus_insula_putamen<br>#6: ibs_visceral_rectal<br>#7: insula_disgust_insular<br>#8: placebo_mg_blind<br>#9: stimulation_somatosensory_representation<br>#10: stimulation_somatosensory_contralateral | #1: blind_sighted_global<br>#2: visual_occipital_early<br>#3: spatial_location_locations<br>#4: navigation_virtual_gait<br>#5: gyrus_fusiform_parahippocampal<br>#6: gaze_eye_eyes<br>#7: visual_auditory_modality<br>#8: presented_visual_presentation<br>#9: motion_mt_visual<br>#10: reference_frame_relations |
| Task: Learn, contrast: incorrect | #1: likelihood_consistency_estimation<br>#2: frontal_inferior_gyrus<br>#3: conflict_response_monitoring<br>#4: task_relevant_irrelevant<br>#5: detection_novelty_oddball<br>#6: task_performing_cognitive<br>#7: reading_phonological_dyslexia<br>#8: rule_rules_abstraction<br>#9: task_switching_set<br>#10: stimulus_response_type | #1: state_resting_seed<br>#2: amplitude_spontaneous_frequency<br>#3: surface_colour_thinning<br>#4: power_gamma_hz<br>#5: stroke_recovery_acute<br>#6: reho_regional_homogeneity<br>#7: network_default_dmn<br>#8: white_matter_tensor<br>#9: lesions_lesion_patient<br>#10: pd_disease_parkinson |
| Task: CharProc, contrast: letter | #1: likelihood_consistency_estimation<br>#2: prefrontal_cortex_dorsolateral<br>#3: dlpfc_prefrontal_cortex<br>#4: cortex_prefrontal_orbitofrontal<br>#5: dacc_dorsal_cingulate<br>#6: empathy_social_empathic<br>#7: cognitive_control_cognition<br>#8: cortex_lateral_prefrontal<br>#9: uncertainty_ambiguous_ambiguity<br>#10: moral_psychopathy_harm | #1: visual_occipital_early<br>#2: gyrus_fusiform_parahippocampal<br>#3: blind_sighted_global<br>#4: gaze_eye_eyes<br>#5: navigation_virtual_gait<br>#6: surface_colour_thinning<br>#7: reference_frame_relations<br>#8: motion_mt_visual<br>#9: frequency_hz_slow<br>#10: spatial_location_locations |
| Task: Localizer, contrast: word–face | #1: reading_letter_chinese<br>#2: reading_phonological_dyslexia<br>#3: reasoning_hearing_deaf<br>#4: cortex_visual_temporal<br>#5: ventral_dorsal_visual<br>#6: detection_novelty_oddball<br>#7: number_numerical_numbers<br>#8: language_english_native<br>#9: task_performing_cognitive<br>#10: words_word_lexical | #1: network_default_dmn<br>#2: mpfc_medial_prefrontal<br>#3: state_resting_seed<br>#4: amplitude_spontaneous_frequency<br>#5: negative_positive_valence<br>#6: fc_resting_state<br>#7: pcc_cingulate_precuneus<br>#8: tinnitus_mindfulness_meditation<br>#9: individual_variability_inter<br>#10: cortex_anterior_cingulate |

*Table S11. Top ten associations of consensus maps, for positive and negative values, and meta-analytical terms for the prediction of the summary score Verbal.*

| Task and contrast | Positive | Negative |
|---|---|---|
| Task: PhonLex, contrast: pseudoword–pseudohomophone | #1: gyrus_fusiform_parahippocampal<br>#2: visual_occipital_early<br>#3: hand_hands_foot<br>#4: gaze_eye_eyes<br>#5: movements_movement_motor<br>#6: motor_cortex_sensorimotor<br>#7: motor_cortex_supplementary<br>#8: force_motor_grip<br>#9: finger_tapping_index<br>#10: hemisphere_hemispheric_lateralization | #1: prefrontal_cortex_dorsolateral<br>#2: judgments_judgment_judged<br>#3: moral_psychopathy_harm<br>#4: deception_truth_lying<br>#5: mpfc_medial_prefrontal<br>#6: network_default_dmn<br>#7: cortex_anterior_cingulate<br>#8: mental_mentalizing_states<br>#9: items_recognition_item<br>#10: dlpfc_prefrontal_cortex |
| Task: Learn, contrast: correct | #1: sma_pre_motor<br>#2: sulcus_intraparietal_ips<br>#3: stimulus_response_type<br>#4: tool_object_hand<br>#5: writing_drawing_figure<br>#6: motor_cortex_supplementary<br>#7: motor_cortex_sensorimotor<br>#8: finger_tapping_index<br>#9: presented_visual_presentation<br>#10: trial_single_task | #1: mpfc_medial_prefrontal<br>#2: mental_mentalizing_states<br>#3: negative_positive_valence<br>#4: individuals_resonance_magnetic<br>#5: tom_mind_theory<br>#6: social_cognition_interactions<br>#7: depression_mdd_depressive<br>#8: network_default_dmn<br>#9: bipolar_bd_disorder<br>#10: symptoms_severity_scores |

*Table S12. Top ten associations of consensus maps, for positive and negative values, and meta-analytical terms for the prediction of the summary score Naming.*

| Task and contrast | Positive | Negative |
|---|---|---|
| Task: PhonLex, contrast: pseudohomophone | #1: taste_swallowing_gustatory<br>#2: anterior_insula_cortex<br>#3: pain_painful_chronic<br>#4: empathy_social_empathic<br>#5: placebo_mg_blind<br>#6: likelihood_consistency_estimation<br>#7: acc_cingulate_anterior<br>#8: insula_disgust_insular<br>#9: thalamus_insula_putamen<br>#10: ibs_visceral_rectal | #1: blind_sighted_global<br>#2: visual_occipital_early<br>#3: gyrus_fusiform_parahippocampal<br>#4: gaze_eye_eyes<br>#5: navigation_virtual_gait<br>#6: visual_auditory_modality<br>#7: spatial_location_locations<br>#8: scenes_scene_ppa<br>#9: presented_visual_presentation<br>#10: motion_mt_visual |
| Task: CharProc, contrast: FFnew | #1: empathy_social_empathic<br>#2: familiar_unfamiliar_familiarity<br>#3: likelihood_consistency_estimation<br>#4: moral_psychopathy_harm<br>#5: sexual_love_romantic<br>#6: dacc_dorsal_cingulate<br>#7: salience_network_sn<br>#8: group_control_individuals<br>#9: limbic_amygdala_paralimbic<br>#10: gestures_gesture_communicative | #1: visual_occipital_early<br>#2: gyrus_fusiform_parahippocampal<br>#3: blind_sighted_global<br>#4: response_hemodynamic_time<br>#5: surface_colour_thinning<br>#6: frequency_hz_slow<br>#7: reference_frame_relations<br>#8: dimensions_horizontal_ap<br>#9: presented_visual_presentation<br>#10: gaze_eye_eyes |
| Task: Learn, contrast: incorrect | #1: frontal_inferior_gyrus<br>#2: likelihood_consistency_estimation<br>#3: task_relevant_irrelevant<br>#4: conflict_response_monitoring<br>#5: prefrontal_cortex_dorsolateral<br>#6: category_categories_categorization<br>#7: rule_rules_abstraction<br>#8: reading_phonological_dyslexia<br>#9: semantic_word_temporal<br>#10: cortex_prefrontal_orbitofrontal | #1: stimulation_somatosensory_contralateral<br>#2: touch_tactile_somatosensory<br>#3: stimulation_somatosensory_representation<br>#4: movements_movement_motor<br>#5: pmc_cord_sci<br>#6: motor_cortex_sensorimotor<br>#7: stroke_recovery_acute<br>#8: motor_cortex_supplementary<br>#9: pd_disease_parkinson<br>#10: cortex_primary_somatosensory |
